# Diffusion barriers and adaptive carbon uptake strategies enhance the modeled performance of the algal CO_2_-concentrating mechanism

**DOI:** 10.1101/2021.03.04.433933

**Authors:** Chenyi Fei, Alexandra T. Wilson, Niall M. Mangan, Ned S. Wingreen, Martin C. Jonikas

**Affiliations:** Department of Molecular Biology, Princeton University, Princeton, NJ 08544; Lewis-Sigler Institute for Integrative Genomics, Princeton University, Princeton, NJ 08544; Department of Engineering Sciences and Applied Mathematics, Northwestern University, Evanston, IL 60208

**Keywords:** CO_2_-concentrating mechanism, *Chlamydomonas reinhardtii*, Rubisco, carbon fixation, algal photosynthesis

## Abstract

Many photosynthetic organisms enhance the performance of their CO_2_-fixing enzyme Rubisco by operating a CO_2_-concentrating mechanism (CCM). Most CCMs in eukaryotic algae supply concentrated CO_2_ to Rubisco in an organelle called the pyrenoid. Ongoing efforts seek to engineer an algal CCM into crops that lack a CCM to increase yields. To advance our basic understanding of the algal CCM, we develop a chloroplast-scale reaction-diffusion model to analyze the efficacy and the energy efficiency of the CCM in the green alga *Chlamydomonas reinhardtii*. We show that achieving an effective and energetically efficient CCM requires a physical barrier such as thylakoid stacks or a starch sheath to reduce CO_2_ leakage out of the pyrenoid matrix. Our model provides insights into the relative performance of two distinct inorganic carbon uptake strategies: at air-level CO_2_, a CCM can operate effectively by taking up passively diffusing external CO_2_ and catalyzing its conversion to HCO_3_^−^, which is then trapped in the chloroplast; however, at lower external CO_2_ levels, effective CO_2_ concentration requires active import of HCO_3_^−^. We also find that proper localization of carbonic anhydrases can reduce futile carbon cycling between CO_2_ and HCO_3_^−^, thus enhancing CCM performance. We propose a four-step engineering path that increases predicted CO_2_ saturation of Rubisco up to seven-fold at a theoretical cost of only 1.5 ATP per CO_2_ fixed. Our system-level analysis establishes biophysical principles underlying the CCM that are broadly applicable to other algae and provides a framework to guide efforts to engineer an algal CCM into land plants.

**Significance Statement:** Eukaryotic algae mediate approximately one-third of CO_2_ fixation in the global carbon cycle. Many algae enhance their CO_2_-fixing ability by operating a CO_2_-concentrating mechanism (CCM). Our model of the algal CCM lays a solid biophysical groundwork for understanding its operation. The model’s consistency with experimental observations supports existing hypotheses about the operating principles of the algal CCM and the functions of its component proteins. We provide a quantitative estimate of the CCM’s energy efficiency and compare the performance of two distinct CO_2_ assimilation strategies under varied conditions. The model offers a quantitative framework to guide the engineering of an algal CCM into land plants and supports the feasibility of this endeavor.

## Introduction

The CO_2_-fixing enzyme Rubisco mediates the entry of roughly 10^14^ kilograms of carbon into the biosphere each year (1–3). However, Rubisco is rather inefficient at performing this essential task (4), fixing CO_2_ at just 10% of its maximum rate under atmospheric levels of CO_2_ (*SI Appendix*, Fig. S1). Moreover, O_2_ competes with CO_2_ for the active site of Rubisco (5), resulting in the loss of fixed carbon and nitrogen through a process known as photorespiration (6). To overcome Rubisco’s inefficiency, many photosynthetic organisms, including cyanobacteria, eukaryotic algae, and some land plants, have evolved CO_2_-concentrating mechanisms (CCMs) (7–10). Such mechanisms elevate CO_2_ levels in the vicinity of Rubisco, thus enhancing CO_2_ fixation and decreasing photorespiration.

Eukaryotic algal CCMs mediate approximately one-third of global CO_2_ fixation (11), yet they remain poorly characterized at a molecular and functional level. In addition to their importance for global biogeochemistry, there is growing interest in engineering an algal CCM into C_3_ crops such as wheat and rice to improve yields and nitrogen- and water-use efficiency (12, 13). Here, we advance our understanding of the eukaryotic algal CCM by developing a reaction-diffusion model of this mechanism based on the molecularly best-characterized alga, *Chlamydomonas reinhardtii* (Chlamydomonas hereafter). As is commonly the case among algae (14), the Chlamydomonas CCM is built around a structure called the pyrenoid, which consists of three elements: (i) a spheroidal matrix, comprised of phase-separated Rubisco (11, 15, 16), (ii) traversing thylakoid tubules (17), which are thought to deliver CO_2_ to the Rubisco (18), and (iii) a surrounding starch sheath, which has been proposed to serve as a diffusion barrier to slow CO_2_ escape from the pyrenoid (19, 20) (Fig. 1A).

**Fig. 1.**
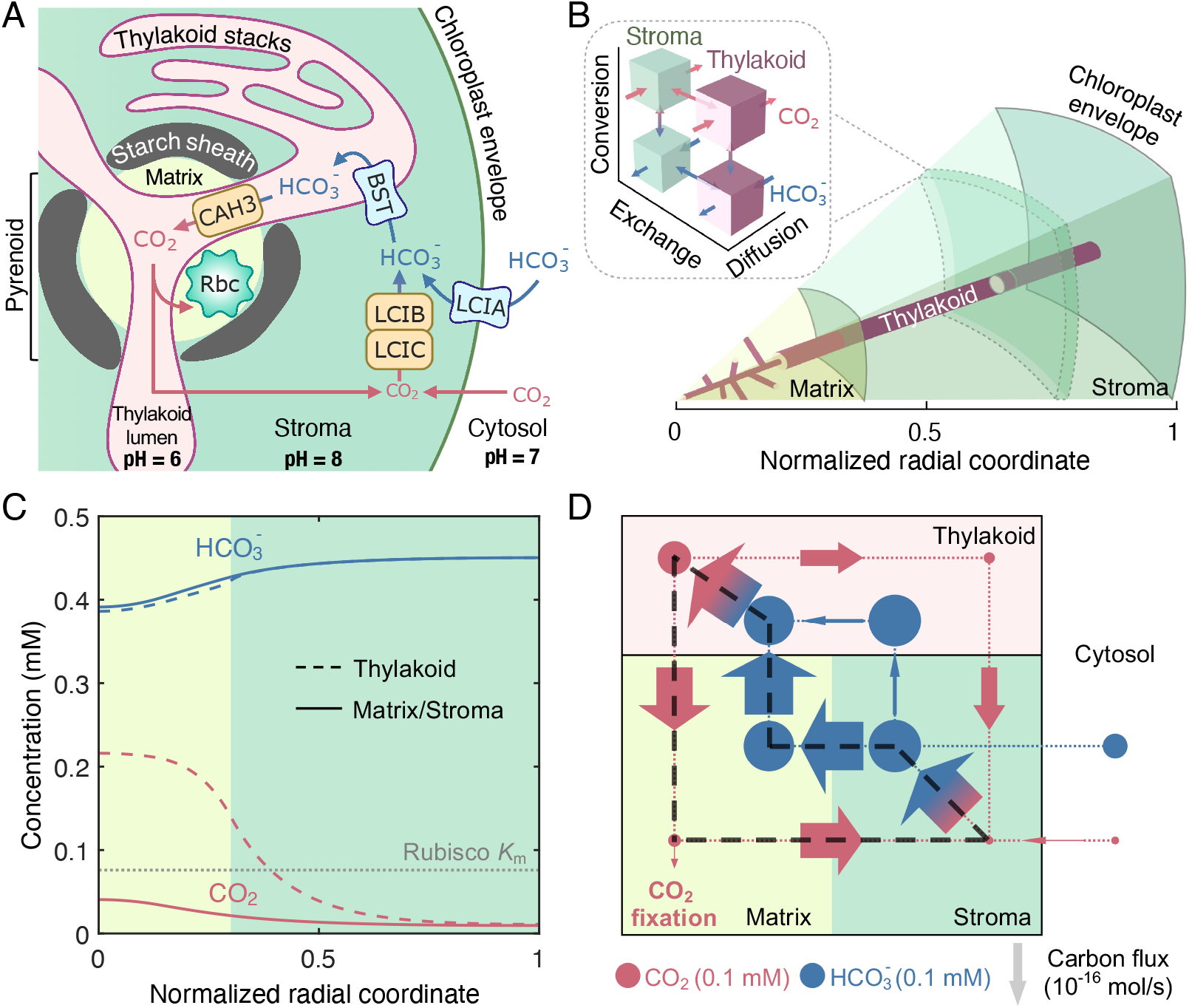
A multi-compartment reaction-diffusion model describes the *Chlamydomonas* CCM. (*A*) Cartoon of a *Chlamydomonas* chloroplast with known CCM components. HCO_3_^−^ is transported across the chloroplast membrane by LCIA and across the thylakoid membranes by BST1–3 (referred to as BST henceforth for simplicity). In the acidic thylakoid lumen, a carbonic anhydrase CAH3 converts HCO_3_^−^ into CO_2_, which then diffuses into the pyrenoid matrix, where the CO_2_-fixing enzyme Rubisco (Rbc) is localized. CO_2_ leakage out of the matrix and out of the chloroplast can be impeded by potential diffusion barriers — a starch sheath and stacks of thylakoids — and by conversion to HCO_3_^−^ by a CO_2_ recapturing complex LCIB/LCIC (referred to as LCIB henceforth for simplicity) in the basic chloroplast stroma. (*B*) A schematic of the modeled CCM, which considers intra-compartment diffusion and inter-compartment exchange of CO_2_ and HCO_3_^−^, as well as their interconversion, as indicated in the *Inset*. Color code as in *A.* The model is spherically symmetric and consists of a central pyrenoid matrix surrounded by a stroma. Thylakoids run through the matrix and stroma; their volume and surface area correspond to a reticulated network at the center of the matrix extended by cylinders running radially outward. (*C*) Concentration profiles of CO_2_ and HCO_3_^−^ in the thylakoid (dashed curves) and in the matrix/stroma (solid curves) for the baseline CCM model that lacks LCIA activity and diffusion barriers. Dotted gray line indicates the effective Rubisco *K*m for CO_2_. Color code as in *A.* (*D*) Net fluxes of inorganic carbon between the indicated CCM compartments. The width of arrows is proportional to flux; area of circles is proportional to the average molecular concentration in the corresponding regions. The black dashed loop denotes the major futile cycle of inorganic carbon in the chloroplast. Color code as in *A*. For *C* and *D*, LCIA-mediated chloroplast membrane HCO_3_^−^ channel transport rate 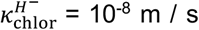, BST-mediated thylakoid membrane HCO_3_^−^ channel transport rate 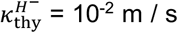, LCIB rate *V*_LCIB_ = 10^3^ / s, and CAH3 rate = 10^4^ / s (see *Materials and Methods* for details). Other parameters of the model are estimated from experiments (Table 1 and *SI Appendix*, Table S1).

**Table 1.** Summary of key parameter values used in the reaction-diffusion model of the algal CCM. See attached pdf.

| Symbols | Descriptions | Values | Ref. |
| --- | --- | --- | --- |
| **Chloroplast geometry** |  |  |  |
| $R_{\text{chlor}}$ | Radius of the chloroplast | 3.14 $\mu\text{m}$ | (17, 81) |
| $R_{\text{pyr}}$ | Radius of the pyrenoid | 0.3 $R_{\text{chlor}}$ | (17, 81) |
| $N_{\text{tub}}$ | Number of thylakoid tubules | 40 | (17) |
| $R_{\text{mesh}}$ | Radius of the tubule meshwork | 0.4 $\mu\text{m}$ | (17) |
| $a_{\text{tub}}$ | Cylindrical radius of the thylakoid tubules | 50 nm | (17) |
| **Kinetic parameters** |  |  |  |
| $V_{\text{max}}^C/K_{\text{m}}^C$ | First-order rate of $\text{CO}_2 + \text{H}_2\text{O} \rightarrow \text{HCO}_3^- + \text{H}^+$ | $10^4 \text{ s}^{-1}$ for CAH3<br>$10^3 \text{ s}^{-1}$ for LCIB | *<br>* |
| $\kappa_{\text{thy}}^{H^-}$ | Rate of $\text{HCO}_3^-$ channels at the thylakoid membranes | $10^{-2} \text{ m/s}$ | * |
| $\kappa_{\text{chlor}}^{H^-}$ | Rate of chloroplast envelope $\text{HCO}_3^-$ transporters | $10^{-8} \text{ m/s}$ | * |

\* Values used in the baseline model. See Fig. 2 and *SI Appendix*, Fig. S4 for a thorough parameter scan. See *SI Appendix*, Table S1 for a full list of model parameters.

CO_2_ is supplied to the pyrenoid by the concerted action of carbonic anhydrases and HCO_3_^−^ transporters (Fig. 1A) (13, 21, 22). External inorganic carbon (Ci: CO_2_ and HCO_3_^−^) is transported across the plasma membrane by LCI1 and HLA3 (23–25), and accumulates in the chloroplast stroma in the form of HCO_3_^−^, either via direct transport across the chloroplast membrane by the HCO_3_^−^ transporter LCIA (24, 26) or via conversion of CO_2_ to HCO_3_^−^ by the putative stromal carbonic anhydrase LCIB/LCIC complex (LCIB hereafter) (27–29). Once in the stroma, HCO_3_^−^ is proposed to travel via the putative HCO_3_^−^ channels BST1–3 (30) into the thylakoid lumen, where the carbonic anhydrase CAH3 (31–33) converts HCO_3_^−^ into CO_2_. This CO_2_ can then diffuse from the thylakoid lumen into the pyrenoid, where Rubisco catalyzes fixation.

Ci fluxes in this system are thought to be powered by two distinct mechanisms: one using pH differences between compartments, and the other using active pumping of HCO_3_^−^ across membranes. Proton pumping during the light reactions of photosynthesis results in a more acidic pH in the thylakoid lumen (pH 6) compared to the stroma (pH 8) (34–36). Since the pH of a compartment dictates the equilibrium ratio of CO_2_ to HCO_3_^−^, the intercompartmental pH differences can drive intercompartmental Ci concentration gradients, and hence Ci fluxes (18, 21). Additionally, any of the transmembrane HCO_3_^−^ transporters could be active pumps that would drive Ci fluxes in the system.

Previous modeling works assuming active HCO_3_^−^ import (37–39) have supported the above mechanisms, but many unanswered questions remain: Does the CCM require active import of Ci? What is the energetic cost of operating the CCM? Do effective CCM strategies change with environmental CO_2_ concentrations? What function does the starch sheath play in the CCM? And, how do the localization patterns of carbonic anhydrases benefit the CCM?

Our reaction-diffusion model suggests that diffusion barriers preventing CO_2_ efflux from the pyrenoid matrix are essential to an effective and energetically efficient CCM. At air-level CO_2_, these diffusion barriers enable a “passive” CCM that lacks any form of active Ci transport and is driven solely by intercompartmental pH differences. Our model of the CCM further reveals two distinct Ci uptake strategies, namely a passive CO_2_ uptake strategy that employs a stromal carbonic anhydrase (LCIB) to convert a diffusive influx of CO_2_ into HCO_3_^−^, and an active HCO_3_^−^ uptake strategy that employs an active pump (LCIA) to directly import external HCO_3_^−^. Moreover, the feasible Ci uptake strategies to support an effective CCM vary with external CO_2_ levels: both strategies function at air-level CO_2_, while active HCO_3_^−^ uptake is necessary under lower CO_2_ conditions. We also demonstrate that proper spatial localization of carbonic anhydrases reduces futile carbon cycling, thereby enhancing CCM performance. Thus, our model illustrates the key biophysical principles necessary to build an effective and energetically efficient algal CCM. Based on these principles, we propose a stepwise engineering path to install an algal CCM into land plants.

## Results

### A multi-compartment reaction-diffusion model of the Chlamydomonas CCM

When adapted to air-level CO_2_ conditions, chloroplasts isolated from Chlamydomonas show an ability to concentrate Ci similar to that of whole cells (40, 41). Thus, we model the chloroplast (Fig. 1A), with constant cytosolic CO_2_ and HCO_3_^−^ concentrations representing external Ci conditions and presumably maintained by the diffusion or import of Ci into the cytosol (see *SI Appendix*).

For simplicity, our reaction-diffusion model is spherically symmetric while taking into account the essential spatial organization of the Chlamydomonas CCM (17). Specifically, we model the chloroplast as a sphere comprised of three compartments: a spherical pyrenoid matrix in the center, a surrounding stroma bounded by a chloroplast envelope, and thylakoids that traverse both the matrix and stroma (Fig. 1B). In Chlamydomonas, the thylakoids enter the pyrenoid matrix in the form of roughly cylindrical membrane tubules, and near the center of the matrix they become interconnected to form a reticulated meshwork (17). In our spherically symmetric model, we account for this geometry implicitly by varying the local volume fraction and surface-to-volume ratio of the thylakoids (Fig. 1B, *Materials and Methods*, and *SI Appendix*, Section IC and Fig. S2).

To understand how Ci is delivered from the cytosol outside the chloroplast to the pyrenoid matrix, we track Ci in each compartment in the form of CO_2_, HCO_3_^−^, and H_2_CO_3_ (with concentrations denoted by *C*, *H*^−^, and *H*^0^, respectively), assuming that H_2_CO_3_ is always in equilibrium with HCO_3_^−^ (42, 43). For simplicity, we assume that the ratio of CO_2_ to HCO_3_^−^ is fixed by *H*^−^ = 10*C* in the cytosol (the consequences of different cytosolic Ci levels are discussed in *SI Appendix*). The first step in Ci concentration is the entry of Ci into the chloroplast. We assume that CO_2_ and H_2_CO_3_ diffuse across the chloroplast envelope at velocities of *κ*^*c*^ = 300 μm/s and 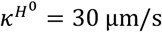, respectively (44, 45). By contrast, HCO_3_^−^ is negatively charged and thus has a much lower baseline velocity 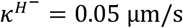 (44). Below, we will refer to the velocity of membrane permeation *κ* as “permeability”. Previous experiments suggest that LCIA, a formate/nitrite transporter homolog, is involved in HCO_3_^−^ transport across the chloroplast envelope (24, 46, 47). We model the action of LCIA with a tunable rate 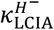. It is presently unclear whether LCIA is an active pump or a passive channel; thus, we model LCIA with a tunable reversibility of inward HCO_3_^−^ transport across the chloroplast envelope (*Materials and Methods*).

Once Ci species enter the chloroplast stroma, the CO_2_ to HCO_3_^−^ ratio is adjusted by the stromal carbonic anhydrase (CA) LCIB, which catalyzes the reaction CO_2_ + H_2_O ↔ HCO_3_^−^ + H^+^ (27–29). Since the interconversion of CO_2_ and HCO_3_^−^ involves a proton, the equilibrium ratio of CO_2_ to HCO_3_^−^, *K*^eq^ = 10^6.1-pH^, depends on the pH in each compartment (48) (*Materials and Methods*). Based on previous measurements and estimates, we set a pH of 8 in the pyrenoid matrix and stroma (34, 35), strongly favoring Ci in the form of HCO_3_^−^ with an equilibrium ratio of approximately *H*^−^ = 80*C*. In our model, the CA-mediated interconversion between CO_2_ and HCO_3_^−^ is described by reversible Michaelis-Menten kinetics (*Materials and Methods* and *SI Appendix*, Section IB). Below, we will refer to the first-order rate constant of CO_2_ to HCO_3_^−^ conversion of a given CA as the “rate” of that enzyme.

All Ci species in the chloroplast can diffuse radially inside compartments and exchange between compartments (Fig. 1B and *Materials and Methods*). In particular, we assume that all Ci species diffuse across the thylakoid membranes, with a permeability of *κ^c^* for CO_2_, 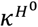 for H_2_CO_3_, and 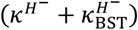 for HCO_3_^−^. Here, the tunable parameter 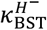 represents the additional permeability to HCO_3_^−^ allowed by bestrophin-like channels that traverse the thylakoid membranes (30).

The lumen of the thylakoids has a pH of 6 (35, 36), favoring a roughly equal partition between CO_2_ and HCO_3_^−^, which is mediated by the luminal carbonic anhydrase CAH3 (31–33). Due to the pH differences, HCO_3_^−^ in the stroma can diffuse into the thylakoid lumen and be converted to CO_2_ by CAH3. This CO_2_ can then diffuse into the pyrenoid matrix and be fixed by Rubisco. We assume that CO_2_ fixation follows Michaelis-Menten kinetics with a maximum rate 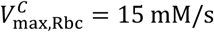, and an effective *K_m_*, 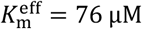, taking into account competitive inhibition by O_2_ (*Materials and Methods*, and *SI Appendix*, Table S1 and Section IB). CO_2_ within the pyrenoid matrix can also diffuse back out into the stroma. Two chloroplast structures have been suggested to act as barriers preventing this diffusion, namely the starch sheath that surrounds the pyrenoid matrix and the stacks of thylakoid membranes present in the chloroplast stroma (Fig. 1A).

In summary, the flux balance of intra-compartment reaction and diffusion and intercompartment exchange sets the steady-state concentration profiles of Ci species in all compartments (Fig. 1B; see *Materials and Methods* for details). All of the model parameters were estimated from the literature (Table 1, and *SI Appendix*, Table S1) except for the enzymatic rates of CAH3 and LCIB and the kinetic parameters of BST and LCIA transporters, for which we performed a systematic scan within a range of reasonable values.

We first present results for our baseline model, with LCIB diffuse throughout the stroma, BST channels uniformly distributed across the thylakoid membranes, CAH3 localized to the thylakoid lumen within the pyrenoid, and Rubisco condensed within the pyrenoid matrix (Fig. 1C and D). The baseline model lacks LCIA and the potential diffusion barriers to Ci mentioned above; we introduce these elements in later sections.

### The baseline CCM model suffers from CO_2_ leakage out of the pyrenoid matrix

A functioning CCM must be out of thermodynamic equilibrium: it elevates the CO_2_ concentration locally around Rubisco. This nonequilibrium system is powered in part by the influx of light energy collected by photosystems I and II, which is used to pump protons into the thylakoid lumen and thereby maintain a pH differential between compartments (49). Thus, we first ask: how effectively can this pH gradient drive CCM function?

In our baseline model, CO_2_ diffusing into the chloroplast is converted to HCO_3_^−^ in the high-pH stroma. Since passive diffusion of HCO_3_^−^ back out across the chloroplast envelope is slow, HCO_3_^−^ becomes trapped in the chloroplast, resulting in a high level of HCO_3_^−^ in the stroma and the thylakoids that traverse it (Fig. 1C). The lower pH in the thylakoid lumen is then leveraged to convert HCO_3_^−^ back to CO_2_ within the pyrenoid radius (Fig. 1D). This CO_2_ can enter the pyrenoid matrix, leading to an enhanced concentration of CO_2_ at the location of Rubisco.

Despite enhancing CO_2_ concentration around Rubisco, our baseline CCM model suffers from significant CO_2_ leakage out of the matrix (Fig. 1D). The first-order rate constant of Rubisco-catalyzed CO_2_ fixation is 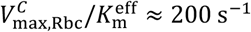. Thus, the average time required for a free CO_2_ molecule to be fixed by Rubisco in the pyrenoid matrix is 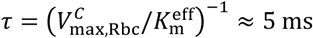. Over the same period of time, that molecule can diffuse over a typical distance (*D*^*C*^τ)^1/2^ ≈ 3 μm, larger than the radius of the pyrenoid *R*_pyr_ ≈ 1 μm. As a result, ~95% of CO_2_ molecules that diffuse into the pyrenoid matrix from the thylakoids leave the matrix without being fixed by Rubisco (*SI Appendix*, Section IH and Fig. S3). Once this CO_2_ reaches the stroma, it is recycled back into HCO_3_^−^ by LCIB. While this HCO_3_^−^ can re-enter the thylakoid lumen where it is again converted into CO_2_ by CAH3 (Fig. 1D, black dashed loop), this recycling of Ci does not enhance the efficacy of the CCM, since the vast majority of CO_2_ released in the pyrenoid does not remain there long enough to be fixed by Rubisco. One might think that increasing the rate at which HCO_3_^−^ diffuses into the chloroplast could overcome this deficit of the system. However, adding LCIA as a passive channel for HCO_3_^−^ at the chloroplast envelope does not solve the intrinsic deficiency caused by CO_2_ leakage. In fact, there is no combination of LCIB catalytic rate and LCIA and BST channel rates that can achieve half-saturation of Rubisco with CO_2_ (Fig. 2 A and B, and *SI Appendix*, Fig. S4 A and B). Thus, in the absence of a diffusion barrier, pH differences alone are not enough to concentrate sufficient CO_2_ in the pyrenoid matrix to half-saturate Rubisco.

**Fig. 2.**
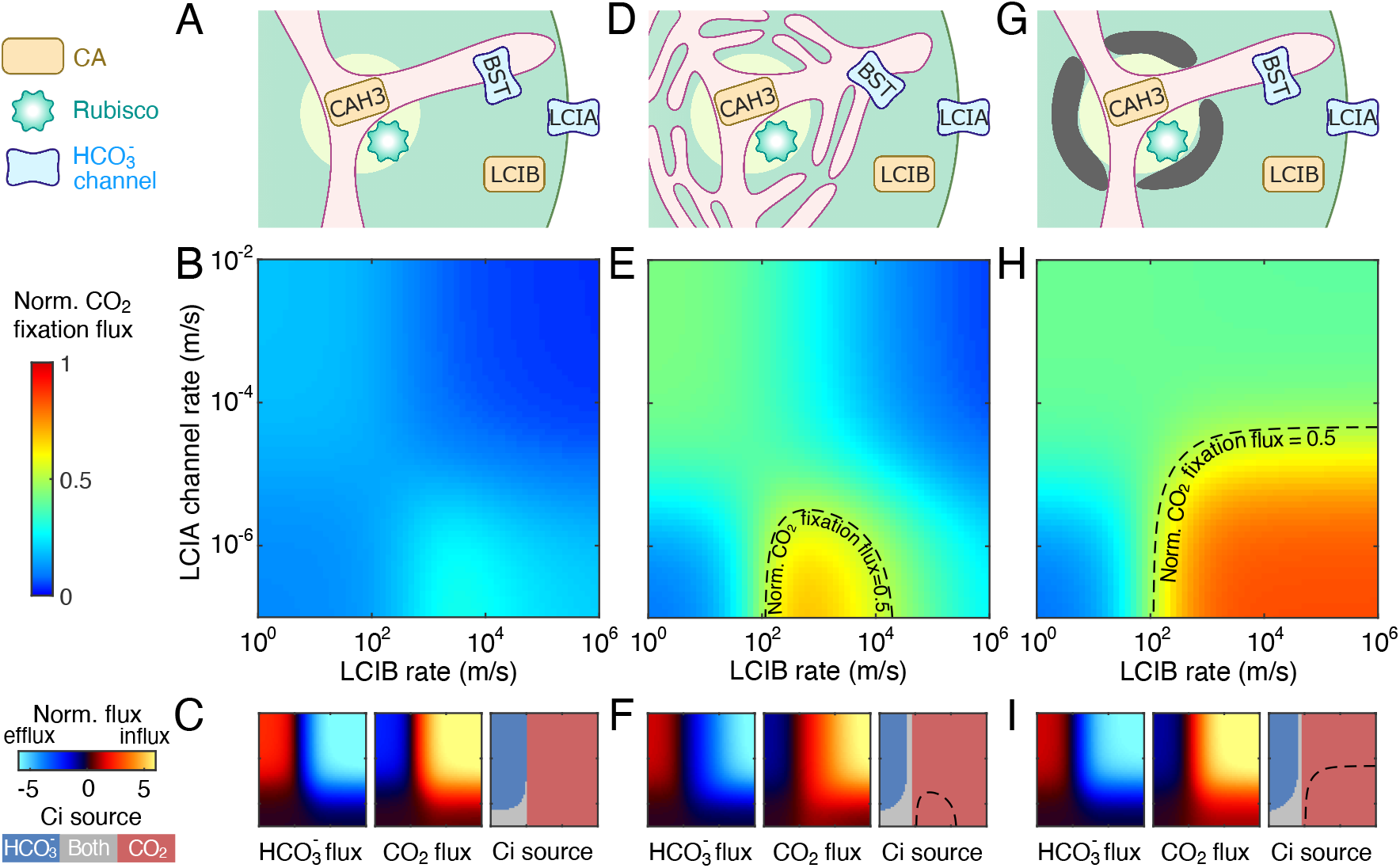
Structural barriers to CO_2_ diffusion out of the pyrenoid matrix enable an effective CCM driven only by intercompartmental pH differences. A model with no barrier to CO_2_ diffusion out of the pyrenoid matrix (*A–C*) is compared to a model with thylakoid stacks slowing inorganic carbon diffusion in the stroma (*D–F*, see *SI Appendix*, Sec. IG) and a model with an impermeable starch sheath (*G–I*) under air-level CO_2_ (10 μM cytosolic). (*A*, *D*, and *G*) Schematics of the modeled chloroplast. (*B*, *E*, and *H*) Heatmaps of normalized CO_2_ fixation flux, defined as the ratio of the total Rubisco carboxylation flux to its maximum if Rubisco were saturated, at varying HCO_3_^−^ channel transport rates across the chloroplast envelope and varying LCIB rates. The HCO_3_^−^ channel transport rate across the thylakoid membrane is the same as in Fig. 1C and D. For *E* and *H*, dashed black curves indicate a normalized CO_2_ fixation flux of 0.5. (*C*, *F*, and *I*) Overall fluxes of HCO_3_^−^ (*Left subpanels*) and CO_2_ (*Middle subpanels*) into the chloroplast, normalized by the maximum CO_2_ fixation flux if Rubisco were saturated, at varying HCO_3_^−^ channel transport rates across the chloroplast envelope and varying LCIB rates. Negative values denote efflux out of the chloroplast. The inorganic carbon (Ci) species with a positive influx is defined as the Ci source, shown in *Right subpanels*. Axes are the same as *B*, *E*, and *H*.

### Structural barriers to CO_2_ diffusion out of the pyrenoid matrix enable an effective CCM driven only by intercompartmental pH differences

In order to operate a more effective CCM, the cell must reduce CO_2_ leakage from the pyrenoid matrix. We examine whether adding a diffusion barrier to the baseline CCM model would be sufficient to concentrate substantially more CO_2_ around Rubisco. We consider two *in vivo* structures of the Chlamydomonas chloroplast as potential barriers for Ci molecules leaving the matrix. The first is the thylakoid stacks, which comprise layers of membranes surrounding the pyrenoid that could effectively slow the diffusion of Ci in the stroma (17, 38) (Fig. 1A). In our spherically symmetric model, we treat the stroma traversed by thylakoid stacks as a homogeneous compartment where diffusion of each Ci species is slowed, as molecules must diffuse between or through the interdigitated thylakoid stacks (*Materials and Methods*). Indeed, a realistic geometry of the thylakoid stacks increases the diffusion path length of Ci in the stroma, reducing effective diffusion coefficients to as low as 3% of their unrestricted values (*SI Appendix*, Section IG and Fig. S5).

Another potential barrier is the starch sheath (Fig. 1A). This sheath forms around the pyrenoid matrix under the same environmental conditions that induce CCM activity in Chlamydomonas and has been suggested to reduce Ci efflux out of the matrix (19, 20). For simplicity, we model the starch sheath as a thin semi-permeable barrier around the matrix, with the same permeability *κ*_starch_ to all Ci species (*Materials and Methods*). Adding a starch sheath to the baseline model creates a discontinuity in carbon concentration across the matrix-stroma interface (*SI Appendix*, Fig. S6). We find that even a relatively high permeability *κ*_starch_~100 μm/s, a value similar to the permeability of a single lipid membrane bilayer to CO_2_, renders the starch barrier effective enough that more than half of CO_2_ leakage from the matrix occurs instead via the thylakoid tubules, which provide passages through the starch barrier (*SI Appendix*, Fig. S7). A somewhat lower starch sheath permeability, equivalent to 10 lipid bilayers, yields a CO_2_ fixation flux almost identical to that of a totally impermeable starch sheath. Since starch consists of many lamellae of crystalline amylopectin (50–52), we hypothesize that its permeability to Ci is low enough that it can be neglected. Thus, we focus below on the case of impermeable starch, i.e., *κ*_starch_ = 0.

We find that adding either thylakoid stacks or a starch sheath to the baseline CCM model drastically reduces CO_2_ leakage from the pyrenoid matrix to the stroma (*SI Appendix*, Figs. S6 and S7). As a result, the addition of either barrier under air-level CO_2_ (10 μM cytosolic) leads to a highly effective CCM that raises CO_2_ concentrations in the matrix above the Rubisco *K*_m_ using only the nonequilibrium pH differential and passive Ci uptake (Fig. 2 E and H). The performance of a model including both barriers closely resembles the case with only an impermeable starch sheath (*SI Appendix*, Fig. S8); thus, we omit such a combined model from further discussion.

### The optimal passive Ci uptake strategy utilizes cytosolic CO_2_, not HCO_3_^−^

In addition to the requirement for a diffusion barrier, the efficacy of the CCM depends on enzyme and HCO_3_^−^ channel rates (Fig. 2). What is the best strategy to passively transport and uptake Ci when the CCM is powered only by the pH differences between compartments?

Since the algal CCM relies on CAH3 to convert HCO_3_^−^ to CO_2_ for fixation by Rubisco, it is important for stromal HCO_3_^−^ to enter the thylakoid lumen and reach CAH3. Indeed, our model shows that Rubisco CO_2_ fixation flux increases with BST passive HCO_3_^−^ channel rates across the thylakoid membranes under all conditions explored in Fig. 2 (*SI Appendix*, Fig. S4).

To achieve an effective CCM, it is equally important to maintain a high level of HCO_3_^−^ in the chloroplast stroma. Depending on LCIB activity, there are two possible passive Ci uptake strategies to achieve this goal. If LCIB activity is low, CO_2_ fixation flux increases with higher rates of the LCIA HCO_3_^−^ channels (Fig. 2 B, E, and H), which facilitates the diffusion of cytosolic HCO_3_^−^ into the chloroplast stroma (Fig. 2 C, F, and I). In contrast, if LCIB activity is high, CO_2_ fixation flux is maximized when LCIA channel rates are low (Fig. 2 B, E, and H); in this case, a diffusive influx of CO_2_ into the chloroplast is converted by LCIB into HCO_3_^−^, which becomes trapped and concentrated in the chloroplast (Fig. 2 C, F, and I). Under this scenario, fast HCO_3_^−^ channel transport across the chloroplast envelope is detrimental, since it allows HCO_3_^−^ converted by LCIB to diffuse immediately back out into the cytosol (Fig. 2 B, E, and H).

Interestingly, for both modeled diffusion barriers we find that the highest CO_2_ fixation flux is achieved by employing LCIB for passive CO_2_ uptake, not by employing LCIA channels for passive HCO_3_^−^ uptake (Fig. 2), even though HCO_3_^−^ is more abundant than CO_2_ in the cytosol. Key to this result is our assumption that the stroma (at pH 8) is more basic than the cytosol (at pH 7.1), which allows LCIB to create a pool of HCO_3_^−^ in the chloroplast stroma at a concentration higher than in the cytosol.

We note that LCIB activity is not always beneficial to CCM function, even when LCIA HCO_3_^−^channel rates are low. Specifically, without a starch sheath, high LCIB activity in the stroma draws more CO_2_ efflux out of the pyrenoid matrix by rapidly converting to HCO_3_^−^ some CO_2_ molecules which would otherwise diffuse back into the matrix (*SI Appendix*, Section IV and Fig. S9). In this case, the highest CO_2_ concentration in the pyrenoid occurs at an intermediate LCIB activity (Fig. 2B and E, and *SI Appendix*, Fig. S9D). As an alternative to blocking CO_2_ escape from the matrix with a starch sheath, this detrimental efflux of CO_2_ can be reduced by localizing LCIB away from the pyrenoid, which leads to a monotonic increase of CO_2_ fixation flux with LCIB activity (*SI Appendix*, Fig. S11).

### Feasible Ci uptake strategies depend on the environmental level of CO_2_

While the passive CO_2_ uptake strategy employing LCIB activity and pH differences can power the CCM under air-level CO_2_ (10 μM cytosolic), the Ci uptake rate is ultimately limited by the diffusion of CO_2_ across the chloroplast envelope. Thus, we questioned whether this strategy is feasible at even lower environmental CO_2_ concentrations. The largest possible flux of CO_2_ diffusing into the chloroplast is *κ*^*c*^*C*_cyt_, which is proportional to the cytosolic CO_2_ concentration *C*_cyt_. Consequently, when *C*_cyt_ is lower than 2 μM, this diffusive influx becomes insufficient to achieve half-saturation of Rubisco with CO_2_ (*SI Appendix*, Section IIID). Indeed, our simulations show that under very low CO_2_ conditions (1 μM cytosolic) (53), a chloroplast using the passive CO_2_ uptake strategy can achieve at most 25% of its maximum CO_2_ fixation flux, even in the presence of barriers to Ci diffusion (Fig. 3).

**Fig. 3.**
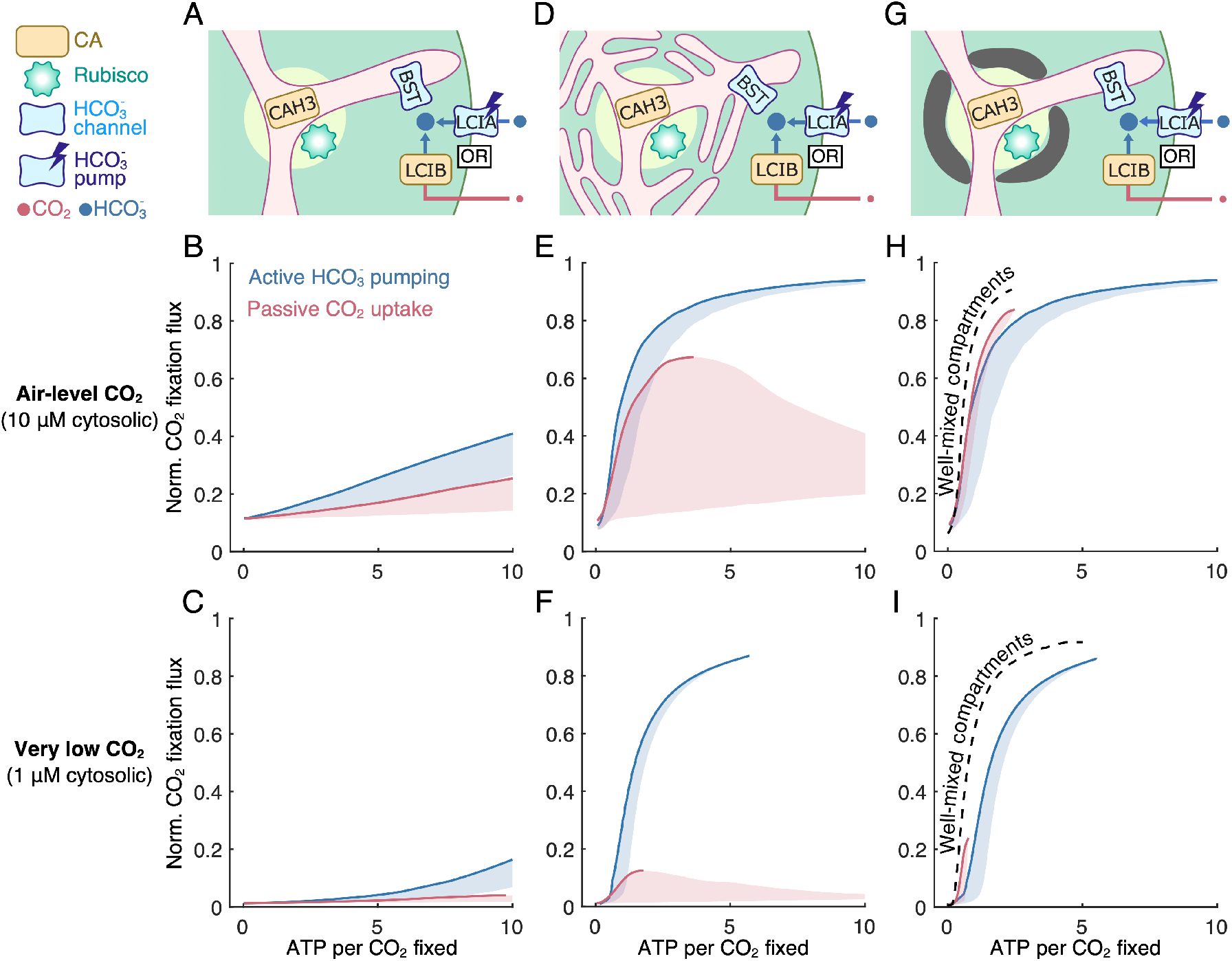
Feasible inorganic carbon uptake strategies for the chloroplast depend on the environmental level of CO_2_. Results are shown for (*A–C*) a model with no barrier to CO_2_ diffusion out of the pyrenoid matrix, (*D–F*) a model with thylakoid stacks serving as diffusion barriers, and (*G–I*) a model with an impermeable starch sheath. (*A*, *D*, and *G*) Schematics of the modeled chloroplast employing LCIB for passive CO_2_ uptake (red), or employing active HCO_3_^−^ pumping across the chloroplast envelope and no LCIB activity (blue). (*B–C*, *E–F*, and *H–I*) CCM performance under (*B*, *E*, and *H*) air-level CO_2_ (10 μM cytosolic) and under (*C*, *F*, and *I*) very low CO_2_ (1 μM cytosolic), measured by normalized CO_2_ fixation flux versus ATP spent per CO_2_ fixed, for the two inorganic carbon uptake strategies in *A*, *D*, and *G*. Solid curves indicate the minimum energy cost necessary to achieve a certain normalized CO_2_ fixation flux. Shaded regions represent the range of possible performances found by varying HCO_3_^−^ transport rates and LCIB rates. Color code as in *A*. For *H* and *I*, dashed black curves indicate the optimal CCM performance of a simplified model that assumes fast intra-compartmental diffusion, fast HCO_3_^−^ diffusion across the thylakoid membranes, and fast equilibrium between CO_2_ and HCO_3_^−^ catalyzed by CAH3 in the thylakoid tubules inside the pyrenoid (see *Materials and Methods* and *SI Appendix*, Sec. III).

What strategies could Chlamydomonas use to continue growing well under very low CO_2_ conditions? Previous research has shown that at very low CO_2_ an LCIB single mutant is viable while an LCIB-LCIA double mutant fails to grow (46), suggesting that HCO_3_^−^ uptake by LCIA is crucial to growth under this condition. Since passive HCO_3_^−^ uptake cannot concentrate more Ci than passive CO_2_ uptake (Fig. 2), we hypothesize that LCIA could actively pump HCO_3_^−^ into the chloroplast under very low CO_2_ conditions. To test whether active HCO_3_^−^ uptake can function as an alternative import mechanism to create a high concentration of stromal HCO_3_^−^, we considered a model employing active LCIA HCO_3_^−^ pumps without LCIB activity (Fig. 3 A, D, and G). We find that, indeed, HCO_3_^−^ pumping enables saturating CO_2_ fixation by Rubisco under both air-level CO_2_ and very low CO_2_ conditions (Fig. 3 and *SI Appendix*, Fig. S12).

While light energy drives the passive CO_2_ uptake strategy by pumping protons across the thylakoid membranes to establish pH differences between compartments, active pumping of HCO_3_^−^ requires additional forms of energy expenditure. What is the energy cost of a CCM that employs active HCO_3_^−^ uptake, and how does this cost compare to that of the passive CO_2_ uptake strategy? To answer these questions, we follow the theoretical framework of nonequilibrium thermodynamics to compute the energy cost of different Ci uptake strategies (*SI Appendix*, Section IIB) (54). Our calculation shows that futile-cycle fluxes, including Ci recycling flux through LCIB and Ci leakage flux out of the chloroplast, increase the energy cost of the CCM (*SI Appendix*, Figs. S13 and S14). Indeed, a chloroplast without diffusion barriers suffers from a very large futile-cycle flux (Fig. 1D) and hence runs a much more energetically expensive CCM than a chloroplast with diffusion barriers (Fig. 3). We note that free energy is also dissipated by nonequilibrium diffusion processes. Thus, a well-mixed compartment model assuming fast intra-compartment Ci diffusion and fast BST-mediated HCO_3_^−^ diffusion across the thylakoid membrane (*Materials and Methods* and *SI Appendix*, Section III and Fig. S15) yields a higher energetic efficiency than the full model with finite rates of diffusion (Fig. 3 H and I, black dashed curves).

Interestingly, the most energy-efficient Ci uptake strategy depends on both the type of diffusion barrier employed and the environmental CO_2_ conditions. At air-level CO_2_ with a starch sheath diffusion barrier, the passive CO_2_ uptake strategy has a slightly higher energy efficiency than the active HCO_3_^−^ uptake strategy (Fig. 3H). Without a starch sheath, however, the active HCO_3_^−^ uptake strategy is energetically less expensive (Fig. 3 B and E). Additionally, we find under very low CO_2_ conditions that no amount of energy input can power CO_2_ concentration to a level higher than the Rubisco *K*_m_ using passive CO_2_ uptake. Therefore, HCO_3_^−^ pumping is required for an effective CCM at very low CO_2_ (Fig. 3 F and I, and *SI Appendix*, Fig. S16). Our results may thus provide mechanistic insights into the observation that Chlamydomonas employs different CO_2_- concentrating strategies depending on the external concentration of CO_2_ (46). Specifically, for a modeled chloroplast with a starch sheath, at air-level external CO_2_ the least costly strategy that allows for half-saturation of Rubisco is activating LCIB for passive CO_2_ uptake. At lower levels of external CO_2_, the least costly strategy is active HCO_3_^−^ pumping across the chloroplast envelope combined with LCIB activity to recapture CO_2_ that leaks from the pyrenoid matrix (*SI Appendix*, Figs. S17 and S18).

### Localization of carbonic anhydrases alters Ci fluxes in the chloroplast

In response to varying external CO_2_ conditions, changes occur not only in the Ci uptake strategy but also in the enzyme localization patterns in the Chlamydomonas CCM. In particular, LCIB localization varies, changing from diffuse throughout the stroma under air-level CO_2_ to localized at the pyrenoid periphery under very low CO_2_ (46, 55). It has also been suggested that CAH3 localizes toward the intra-pyrenoid portion of the thylakoid tubules under air-level CO_2_ (33). These findings prompted us to wonder whether CA localization could impact the performance of the modeled CCM.

To explore this question, we vary the start radius 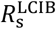 of LCIB, i.e., how close to the chloroplast center LCIB localization starts, and the end radius 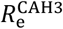 of CAH3, i.e., how far CAH3 extends through the thylakoid tubules (Fig. 4A). We explore this in our spherically symmetric model while maintaining the total number of molecules of each CA. Our simulations reveal three CA localization patterns that would compromise CCM performance. First, when LCIB extends into the pyrenoid matrix, i.e., when 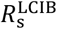 is smaller than the pyrenoid radius *R*_pyr_, LCIB converts Rubisco’s substrate, CO_2_, into HCO_3_^−^. Since HCO_3_^−^ cannot be fixed by Rubisco, this localization of LCIB decreases CO_2_ fixation (Fig. 4 B, D, and F, region i). Second, when CAH3 is distributed in the thylakoids outside the pyrenoid, CO_2_ molecules produced by this CAH3 may diffuse directly into the stroma, where they can be converted to HCO_3_^−^ by LCIB. While this HCO_3_^−^ can then diffuse back into the thylakoid lumen and undergo conversion to CO_2_ again, such futile cycling decreases both the efficacy and energy efficiency of the CCM (Fig. 4 B–F, region ii, and *SI Appendix*, Fig. S19). Finally, concentrating CAH3 to a small region of thylakoid lumen in the center of the pyrenoid increases the distance over which HCO_3_^−^ needs to diffuse before it is converted to CO_2_, thus lowering the CO_2_ production flux by CAH3 (Fig. 4 B, D, and F, region 3). All these results hold true both at air-level CO_2_ employing passive CO_2_ uptake (Fig. 4) and at very low CO_2_ employing active HCO_3_^−^ uptake (*SI Appendix*, Fig. S20). Thus, our model shows that proper CA localization is crucial to overall CCM performance.

**Fig. 4.**
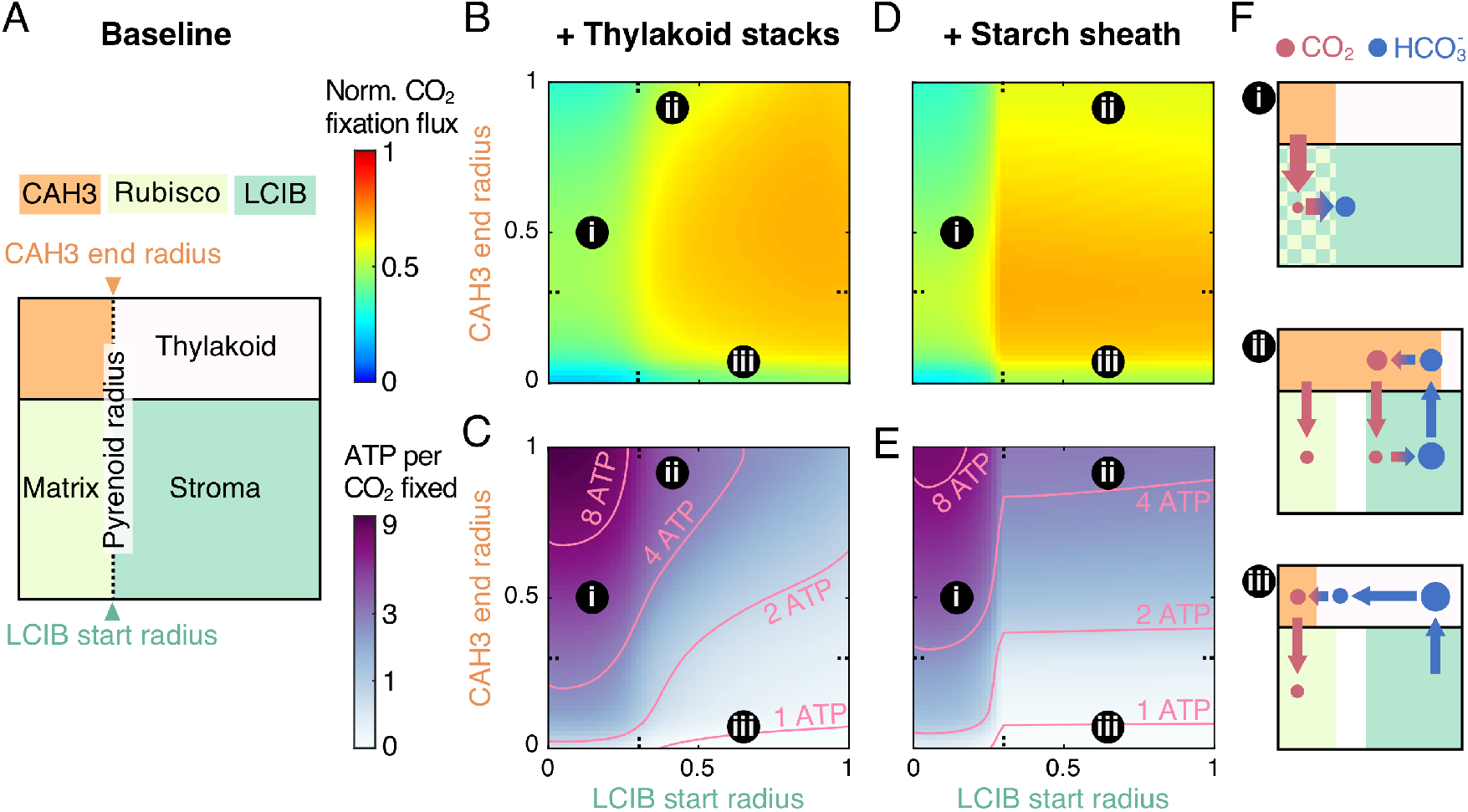
Localization of carbonic anhydrases alters Ci fluxes in the chloroplast and enhances CCM performance. (*A*) Schematics of varying localization of carbonic anhydrases. The CAH3 domain starts in the center of the intra-pyrenoid tubules (radius *r* = 0), and the LCIB domain ends at the chloroplast envelope. CAH3 end radius and LCIB start radius are varied in a modeled chloroplast employing the passive CO_2_ uptake strategy under air-level CO_2_, (*B–C*) with thylakoid stacks slowing inorganic carbon diffusion in the stroma or (*D–E*) with an impermeable starch sheath. Color code is the same as in Fig. 1D. Orange denotes region occupied by CAH3. (*B*, *D*) Normalized CO_2_ fixation flux and (*C*, *E*) ATP spent per CO_2_ fixed when the localizations of carbonic anhydrases are varied. (*F*) Schematics of inorganic carbon fluxes for the localization patterns indicated in *B–E*. Color code is the same as in *A* and Fig. 1D. Simulation parameters are the same as in Fig. 1C and D.

### Activity and localization of LCIB could reduce Ci leakage out of the chloroplast

To better understand the role of LCIB and its *in vivo* localization pattern at very low CO_2_, we next consider a model employing HCO_3_^−^ pumping across the chloroplast envelope. Here, we fix 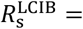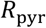 and vary both the end radius of LCIB, 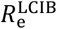, which defines how far LCIB extends toward the chloroplast envelope, and the total number of LCIB molecules (Fig. 5A). In the absence of a starch sheath, localizing LCIB to the pyrenoid periphery harms the CCM; such localization results in a large CO_2_ efflux out of the matrix due to rapid conversion to HCO_3_^−^ (*SI Appendix*, Fig. S21). Thus, we focus on a model employing a starch sheath barrier. Since actively accumulating Ci in the form of HCO_3_^−^ costs energy, it is energetically wasteful if any Ci molecules diffuse out of the chloroplast without being fixed (Fig. 5 B–C, region iii). Consequently, localizing LCIB near the starch sheath increases energy efficiency by recapturing CO_2_ molecules that diffuse out of the matrix and trapping them as HCO_3_^−^ in the chloroplast (Fig. 5 B–C, region i). In contrast, diffuse LCIB is suboptimal because LCIB near the chloroplast envelope could rapidly convert HCO_3_^−^ pumped into the chloroplast into CO_2_, which can then immediately diffuse back out into the cytosol (Fig. 5 B–C, region ii). This futile cycle occurs when HCO_3_^−^ pumping across the chloroplast envelope is fast and irreversible (*SI Appendix*, Fig. S22). Our model thus suggests that under very low CO_2_ and in the presence of a strong CO_2_ diffusion barrier around the pyrenoid, localizing LCIB at the pyrenoid periphery allows for efficient Ci recycling, therefore enhancing CCM performance.

**Fig. 5.**
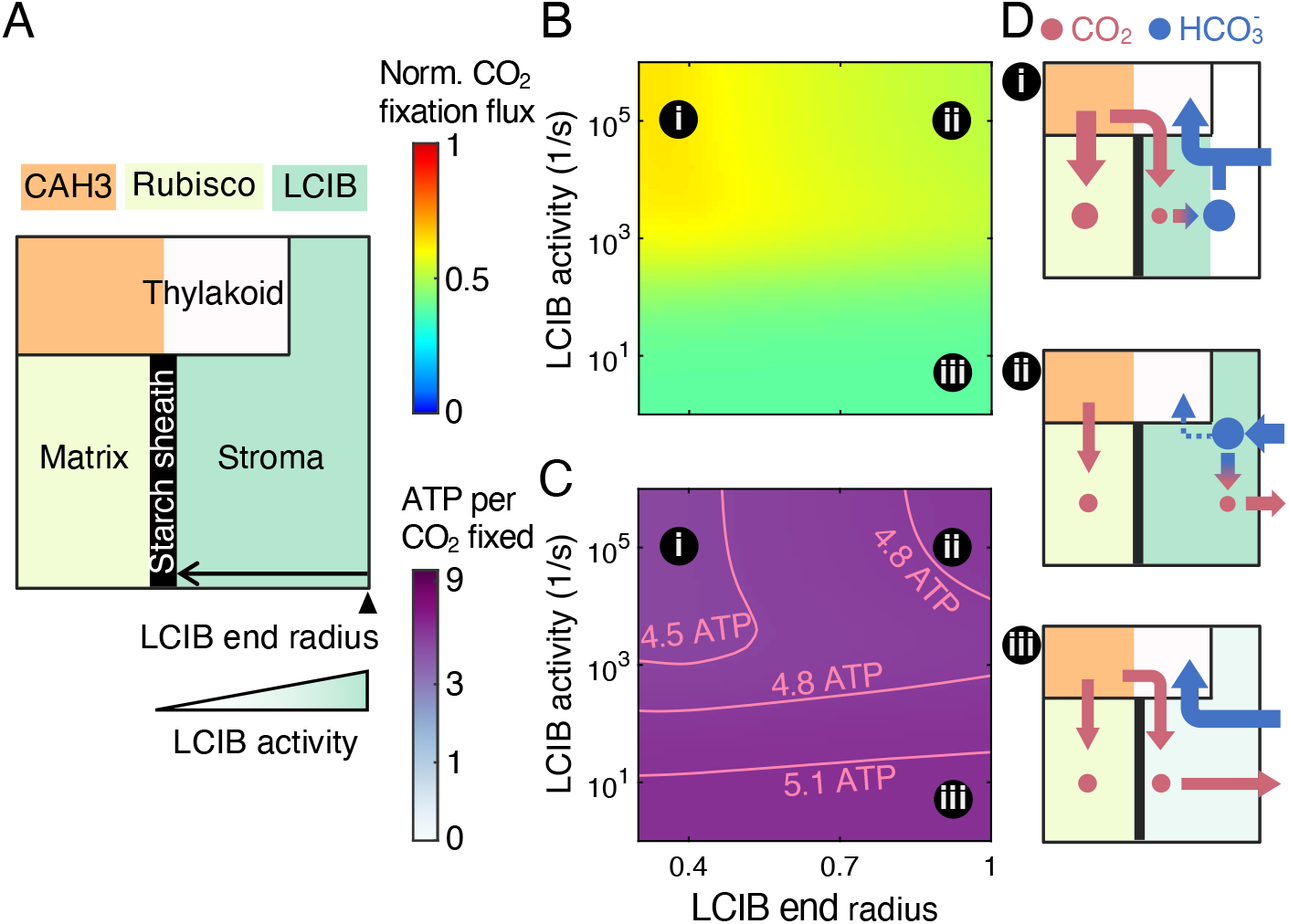
Localization of LCIB around the pyrenoid periphery reduces Ci leakage out of the chloroplast. (*A*) Schematics of varying activity and end radius of LCIB in a modeled chloroplast employing an impermeable starch sheath and active HCO_3_^−^ pumping across the chloroplast envelope under very low CO_2_. Color code as in Fig. 4A. The LCIB domain starts at the pyrenoid radius (0.3 on the *x* axis in *B* and *C*). (*B*) Normalized CO_2_ fixation flux and (*C*) ATP spent per CO_2_ fixed when the designated characteristics of LCIB are varied. (*D*) Schematics of inorganic carbon fluxes for the LCIB states indicated in *B* and *C*. Color code as in Fig. 4F. Simulation parameters as in Fig. 4. Active HCO_3_^−^ pumping is described by the rate *κ*_chlor_ = 10^−4^ m / s and the reversibility γ = 10^−4^. In order to show a notable variation in normalized CO_2_ fixation flux, a model with shortened thylakoid tubules is simulated (see *Materials and Methods*). The qualitative results hold true independent of this specific choice.

### Possible stepwise engineering strategies for transferring algal CCM components to land plants

Many land plants, including most crop plants, are thought to lack any form of CCM. Engineering an algal CCM into land plants has emerged as a promising strategy to potentially increase crop yields through enhanced CO_2_ fixation (12, 13). Despite early engineering advances (47, 56), it remains to be determined what minimal set of engineering steps is needed and in what order these steps should be implemented to establish an effective algal CCM in a plant chloroplast.

To address this question within our model, we measured the efficacy and energetic efficiency of 216 configurations of chloroplast-based CCM, varying the presence and localization of Rubisco, thylakoid and stromal CAs, HCO_3_^−^ channels on the thylakoid membranes and the chloroplast envelope, and diffusion barriers (*SI Appendix*, Fig. S23). Note that we restrict our focus to modeled CCMs that employ passive Ci uptake strategies at 10 μM cytosolic CO_2_, which is close to the CO_2_ levels experienced by plant chloroplasts (57). The use of the passive Ci uptake strategy simplifies the engineering problem by eliminating the needs to engineer active HCO_3_^−^ transport at the chloroplast envelope and to decrease carbonic anhydrase activity in the stroma.

To the best of our knowledge, the typical land plant chloroplast contains diffuse CA and diffuse Rubisco in the stroma, and lacks HCO_3_^−^ channels and diffusion barriers (58) (Fig. 6A). Studies have also suggested the presence of native plant CAs diffuse in the thylakoid lumen (59), so we have included these CAs in our modeled plant chloroplast configuration. This configuration supports only 10% of the maximum CO_2_ fixation flux through Rubisco, and its efficacy is identical to that of the same configuration without thylakoid CAs (*SI Appendix*, Table S4). By contrast, the configuration that achieves the highest CO_2_ fixation flux, >70% of the maximum, corresponds to a Chlamydomonas chloroplast employing passive CO_2_ uptake and a strong diffusion barrier around the pyrenoid (Figs. 2D and 6A).

**Fig. 6.**
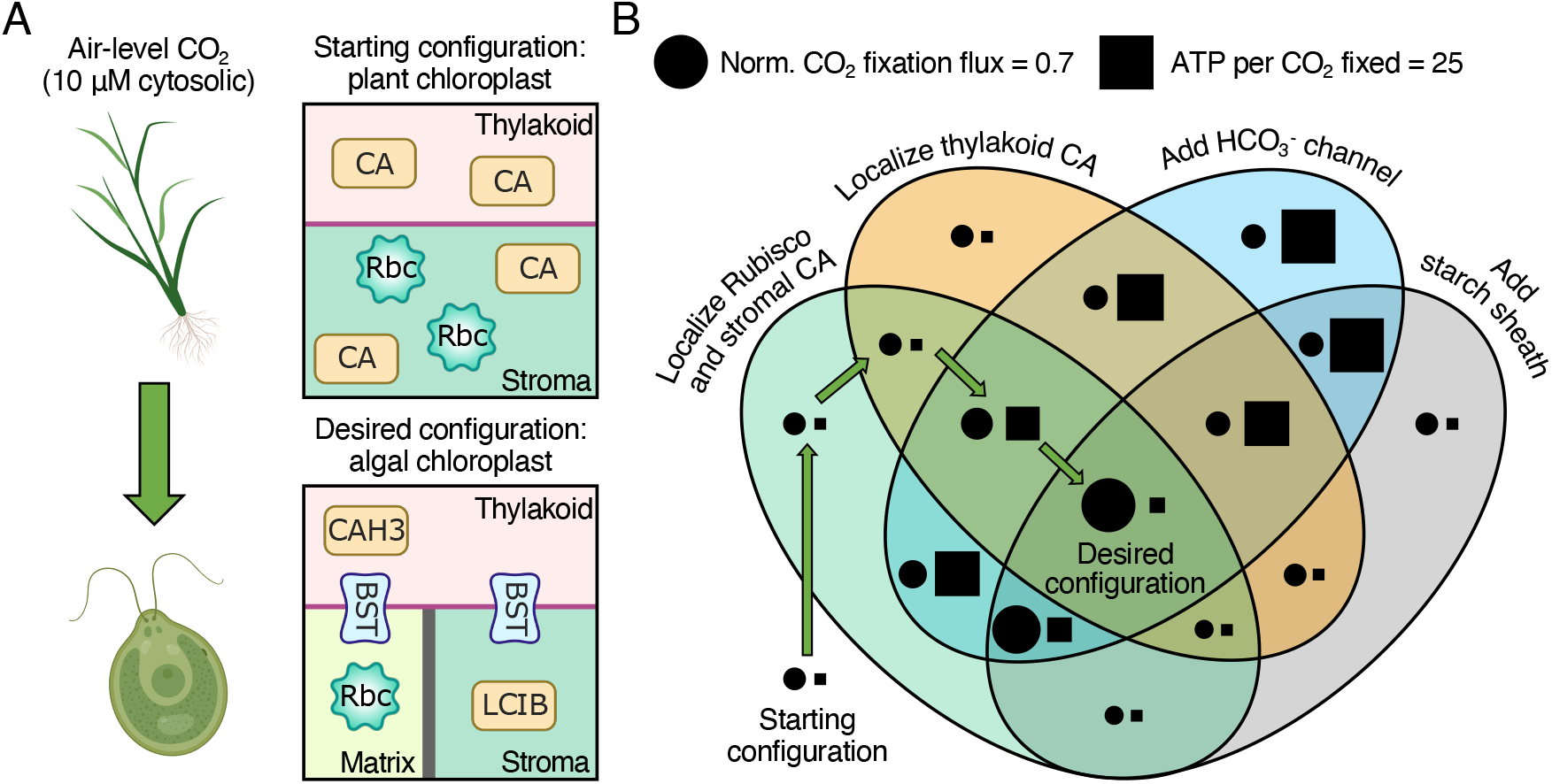
Proposed engineering path for installing an algal CCM into land plants. (*A*) Schematics of (*Top*) the starting configuration representing a typical plant chloroplast that contains diffuse thylakoid CA, diffuse stromal CA, and diffuse Rubisco, and lacks HCO_3_^−^ transporters and diffusion barriers, and (*Bottom*) the desired configuration representing a Chlamydomonas chloroplast that employs the passive CO_2_ uptake strategy and a starch sheath (as in Fig. 2G). (*B*) Venn diagram showing the normalized CO_2_ fixation flux (circle, area in proportion to magnitude) and ATP spent per CO_2_ fixed (square, area in proportion to magnitude) of various configurations after implementing the designated changes. Green arrows denote the proposed sequential steps to transform the starting configuration into the desired configuration (see text). The starting configuration has a normalized CO_2_ fixation flux of 0.11 and negligible ATP cost. All costs below 0.8 ATP per CO_2_ fixed are represented by a square of the minimal size.

We next chart an engineering path from the configuration representing a plant chloroplast (Fig. 6, starting configuration) to the Chlamydomonas configuration that maximizes CO_2_ fixation flux (Fig. 6, desired configuration). While forming the pyrenoid matrix through the condensation of Rubisco and excluding LCIB from this matrix could be considered two separate engineering steps, previous research suggests that the matrix might inherently exclude proteins larger than ~78 kDa (55). Since the plant stromal CA is thought to form complexes with a molecular weight larger than 78 kDa (60), we assume that localizing Rubisco into a matrix and localizing the plant stromal CA outside that matrix can be achieved in a single engineering step. Thus, the four necessary engineering steps are to localize Rubisco and the stromal CA, to localize the thylakoid CA, to add HCO_3_^−^ channels spanning the thylakoid membranes, and to add a starch sheath around the newly created pyrenoid matrix (Fig. 6B). (Note that for simplicity we consider adding HCO_3_^−^ channels uniformly to both the matrix/thylakoid interface and the stroma/thylakoid interface.) In addition, the spatial proximity between the pyrenoid matrix and the thylakoids is important for the engineered CCM, which we address in the Discussion.

We find that the order in which these engineering steps are implemented matters for the efficacy and efficiency of the CCM in intermediate stages. Notably, adding HCO_3_^−^ channels on the thylakoid membranes before the stromal and thylakoid CAs are localized leads to an energetically inefficient configuration (Fig. 6B, blue oval) due to the futile cycling generated by overlapping CAs (Fig. 4, region ii). Additionally, adding a starch sheath before HCO_3_^−^ channels are added to the thylakoids does not increase CO_2_ fixation (Fig. 6B, gray oval), because without channels HCO_3_^−^ cannot readily diffuse to the thylakoid CA to produce CO_2_, and the diffusion of CO_2_ to Rubisco from the stroma is impeded by a starch sheath.

Finally, we suggest a four-step engineering path that avoids intermediate configurations with decreased efficacy or extreme energetic inefficiency (Fig. 6B, green arrows): The first two steps are the localization of Rubisco and the stromal CA and the localization of the thylakoid CA to the thylakoids inside the newly formed pyrenoid matrix. These steps do not yield notable changes to either the efficacy or the efficiency of the CCM, and they could be implemented in either order. The next step is to introduce HCO_3_^−^ channels to the thylakoid membranes, which increases the CO_2_ fixation flux by ~100%. This step also increases the cost of the CCM to around 10 ATP per CO_2_ fixed; such a high-cost step cannot be avoided, and all other possible paths with increasing efficacy at each step have more costly intermediate configurations (Fig. 6B and Table S4). The final step of the suggested path is to add a starch sheath, which drastically increases the efficacy and energy efficiency of the CCM by blocking CO_2_ leakage from the pyrenoid matrix.

One additional benefit of this path is that it provides opportunities for assessing the success of introducing HCO_3_^−^ channels spanning the thylakoid membranes. The increased CO_2_ fixation flux resulting from this step in the proposed path would provide evidence that the installed channels are functional, and could also be used to apply a selective pressure to aid engineering in the event that merely transforming BST channels into plants does not yield HCO_3_^−^ transport across the thylakoid membranes.

### An effective CCM requires Ci uptake, transport, and trapping

What are the essential building blocks of an effective pyrenoid-based CCM? Investigating the performances of the various CCM configurations described in the previous section reveals three central modules of an effective CCM (Fig. 7A): (i) an effective Ci uptake strategy that employs either a carbonic anhydrase (LCIB) to convert a diffusive influx of CO_2_ into HCO_3_^−^ or an active pump (LCIA) to import external HCO_3_^−^ into the chloroplast (Fig. 3), (ii) a system consisting of an HCO_3_^−^ channel (BST) in the thylakoid membranes and another carbonic anhydrase (CAH3) that together transport HCO_3_^−^ to near Rubisco and then convert the HCO_3_^−^ to CO_2_, and (iii) a pyrenoid matrix that houses Rubisco, surrounded by diffusion barriers that trap CO_2_ inside the matrix. We find that CCM configurations lacking any one of these modules show a compromised ability to concentrate CO_2_ (Fig. 7B). Thus, our characterization illustrates the minimal functional modules for an algal CCM.

**Fig. 7.**
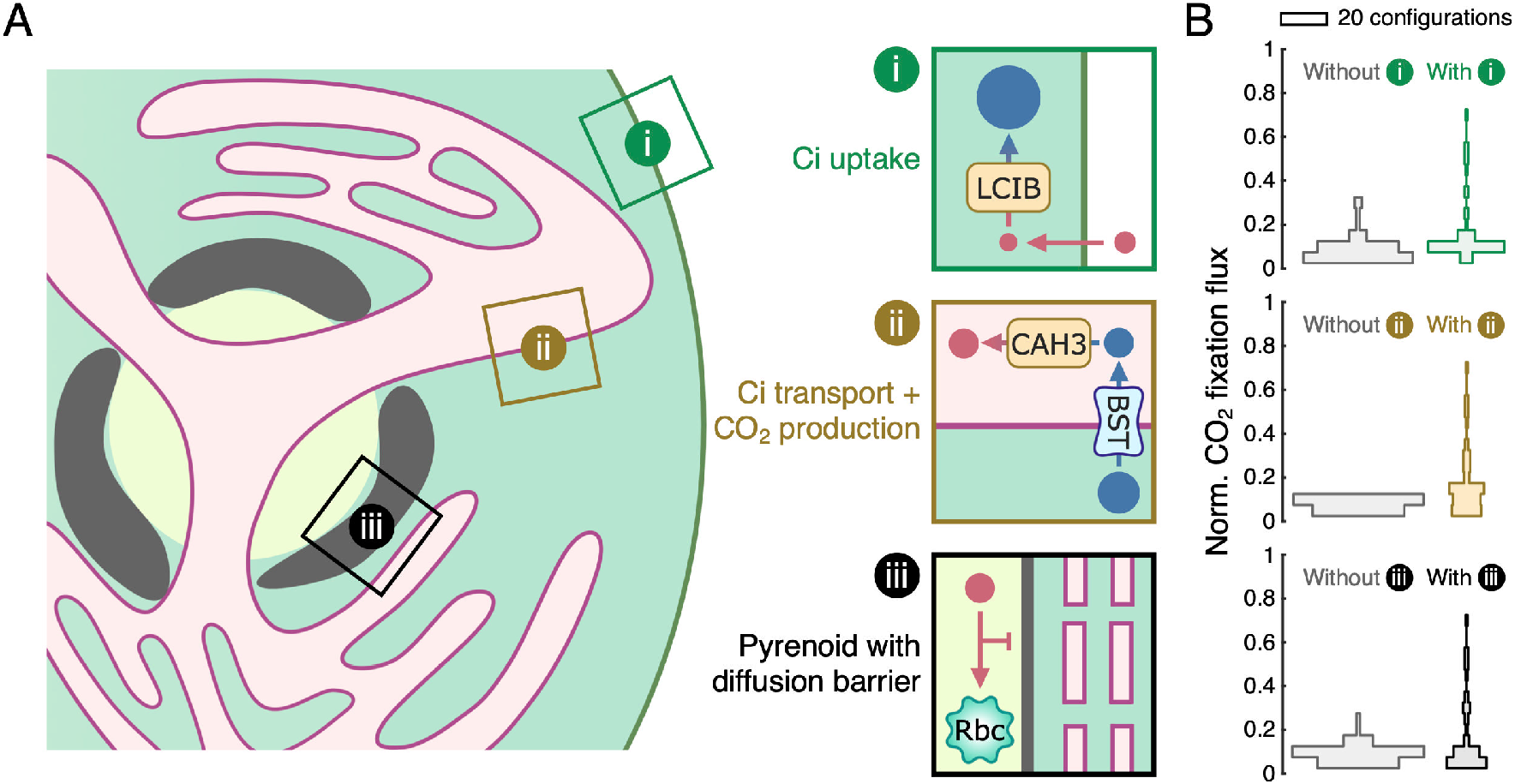
An effective CCM is comprised of three essential modules. (*A*) Schematics of the three essential modules with designated functions. Same style as in Fig. 1A. In Chlamydomonas, LCIB can be used for passive uptake of CO_2_ which is then trapped in the stroma as HCO_3_^−^ (module i); BST allows stromal HCO_3_^−^ to diffuse into the thylakoid lumen where CAH3 converts HCO_3_^−^ into CO_2_ (module ii); and a starch sheath and thylakoid stacks could act as diffusion barriers to slow CO_2_ escape out of the pyrenoid matrix (module iii). (*B*) Histograms of normalized CO_2_ fixation flux for CCM configurations without (*left*, gray) or with (*right*, colored) the respective module. We tested 216 CCM configurations by varying the presence and/or localization of enzymes, HCO_3_^−^ channels, and diffusion barriers in the model (see *SI Appendix*, Fig. S23).

## Discussion

The algal CCM elevates CO_2_ around Rubisco, thereby enhancing photosynthesis. This CCM has the potential to be transferred into crop plants to increase their photosynthetic efficiency. To better understand how the algal CCM works, we develop a multi-compartment reaction-diffusion model based on the Chlamydomonas chloroplast. We provide a quantitative framework to evaluate overall CCM performance, considering both the efficacy and the energetic efficiency of the CCM (*SI Appendix*, Sec. IIC and Fig. S24). While previous works have suggested the operational principles underlying an effective CCM, our analysis lays a quantitative and biophysical groundwork for these principles as discussed below.

According to our model, a diffusion barrier that blocks CO_2_ leakage out of the pyrenoid matrix is essential to an effective CCM in Chlamydomonas. Indeed, previous modeling works have demonstrated the necessity of diffusion barriers for the cyanobacterial CCM (61, 62). Recent experiments in Chlamydomonas showed that mutants either lacking the starch sheath or having a thinner starch sheath have decreased CO_2_-concentrating activity, suggesting that the starch sheath is required for an effective CCM (20). Further experiments are needed to clarify the role of starch sheath in the CCM, and physical properties of the starch sheath such as its permeability to inorganic carbon need to be defined. Our modeling results provide additional testable predictions for experiments to examine a role for starch as a barrier to CO_2_ escape. For example, if indeed the starch sheath functions as an effective diffusion barrier, our model predicts that overexpressing LCIB in wildtype cells will not affect their growth in air (Fig. 2H). By contrast, we predict that overexpressing LCIB in a starchless mutant of Chlamydomonas will lead to a growth defect under air-level CO_2_ — the LCIB will effectively “pull” CO_2_ from the pyrenoid matrix (Fig. 2E). Testing these mutant phenotypes will shed light on the nature of the diffusion barriers present in the algal CCM.

Our results demonstrate two distinct Ci uptake strategies, i.e., employing LCIB for passive CO_2_ uptake and employing LCIA for active HCO_3_^−^ pumping. Our modeling shows that an effective CCM must use the active HCO_3_^−^ uptake strategy to function under very low CO_2_. Thus, we suggest that the algal CCM may switch from passive CO_2_ uptake under air-level CO_2_ to active HCO_3_^−^ uptake under very low CO_2_. This proposal is consistent with and provides a mechanistic explanation for previous experiments studying LCIA and LCIB mutants (27, 46, 63): compared to wildtype cells, the *lcia* mutant shows a significantly decreased photosynthetic activity across varying external Ci levels at pH 9.0 where almost all Ci is in the form of HCO_3_^−^, presumably due to the lack of a functional HCO_3_^−^ uptake system (46). The photosynthetic activity of the *lcia* mutant is raised at pH 7.3, where more Ci is in the form of CO_2_, consistent with our model that LCIB facilitates CO_2_ uptake in this mutant. The *lcib* mutant fails to grow in air, presumably due to the lack of a functional CO_2_ uptake system, but recovers growth under very low CO_2_ — an effect we attribute to the activation of an HCO_3_^−^ uptake system under this condition (24, 46). Indeed, further knockdowns of genes encoding putative HCO_3_^−^ transporters LCIA or HLA3 in the *lcib* mutant result in dramatic decreases in Ci uptake and growth under very low CO_2_ (23, 46). Our model predicts that there must be an active HCO_3_^−^ pump in Chlamydomonas, possibly LCIA, which has yet to be shown experimentally. At this time, only the diatom and bacterial homologs of LCIB have been explicitly shown to have carbonic anhydrase activity (29), but our model and the observed *lcib* mutant phenotypes together strongly suggest that LCIB has carbonic anhydrase activity. Future functional characterizations of Chlamydomonas LCIA and LCIB will be crucial to verify their proposed roles in Ci uptake. Interestingly, the additional mutation of *cah3* in the *lcib* mutant rescues the air-dier phenotype of the latter (63), while our modeled configurations corresponding to these two mutants show similarly low CO_2_ concentrating activity (Table S4), which would predict that they both have growth defects. Further experimentation is thus required to shed light on how Ci is taken up by the chloroplast and how sufficient CO_2_ reaches the pyrenoid in the *cah3-lcib* double mutant.

Compared to the active HCO_3_^−^ uptake strategy, our model suggests that the passive CO_2_ uptake strategy has a similar performance under air-level CO_2_ and has a much lower efficacy under very low CO_2_ (Fig. 3). One may ask: what are the potential benefits of employing passive CO_2_ uptake? One possibility is that, by employing both Ci uptake strategies, the algal CCM can remain effective under environments with various Ci compositions (*SI Appendix*, Fig. S25). Another possibility regards the feasibility of maintaining cytosolic Ci levels. Since there is no known carbonic anhydrase in the Chlamydomonas cytosol, maintaining HCO_3_^−^ levels requires expressing HCO_3_^−^ transporters at the cell membrane, while cytosolic CO_2_ can be replenished by external CO_2_ diffusing across the cell membrane.

In addition to the Ci uptake strategy employed, the localization of LCIB also changes *in vivo* in response to different external Ci levels. Specifically, under air-level CO_2_ LCIB is distributed throughout the stroma, while the enzyme is localized around the starch sheath under very low CO_2_ (28, 46, 55). Our model suggests that such differences in localization could improve performance under the corresponding Ci uptake strategies: when the passive CO_2_ uptake strategy is used at air-level CO_2_, varying LCIB localization only minimally affects CCM performance as long as the enzyme is confined to the stroma (*SI Appendix*, Fig. S21). In contrast, when HCO_3_^−^ is actively pumped into the chloroplast, localizing LCIB near the pyrenoid periphery and away from the chloroplast envelope is advantageous to avoid the conversion of HCO_3_^−^ near the chloroplast envelope into CO_2_, which can immediately leak back out into the cytosol (Fig. 5). Thus, our model predicts that diffuse LCIB in the chloroplast *in vivo* at very low external CO_2_ could lead to a CCM-deficient phenotype — similar to the phenotype observed when an exogenous CA was introduced diffusely into the cytosol of cyanobacteria that employ active HCO_3_^−^ transport across the cell membrane (7). Understanding experimentally how LCIB localization impacts Ci fluxes in the chloroplast would advance our understanding of the Chlamydomonas CCM.

The analysis of Ci fluxes in our model supports the long-held view that the thylakoid tubules traversing the pyrenoid and converging in the pyrenoid center are capable of delivering stromal HCO_3_^−^ to the pyrenoid, where it can be converted to CO_2_ by CAH3 (18, 21). However, this unique architecture of thylakoid tubules is not essential for transporting HCO_3_^−^ and producing CO_2_. Indeed, algae display a variety of thylakoid tubule morphologies, such as multiple non-connecting parallel thylakoid stacks passing through the pyrenoid, a single disc of thylakoids bisecting the pyrenoid matrix, or thylakoid sheets surrounding but not traversing the pyrenoid (64–67). Our calculations support the idea that different morphologies could in principle allow the functioning of an effective CCM, as long as HCO_3_^−^ can diffuse into the low-pH thylakoid lumen and the thylakoid CA is localized near the pyrenoid to convert HCO_3_^−^ to CO_2_ (*SI Appendix*, Fig. S26).

In our model, thylakoid tubules traversing the starch sheath are the main route for CO_2_ escape from the pyrenoid, which is detrimental to CO_2_ concentration. Additionally, this particular geometry is not required for HCO_3_^−^ delivery to the pyrenoid (*SI Appendix*, Fig. S26 D*–*F). Nevertheless, Chlamydomonas cells appear to maintain these tubule structures *in vivo*. It is possible that, by connecting to the thylakoid network outside of the pyrenoid, thylakoid tubules can also serve as a route for protons to diffuse in, which helps to maintain the acidic pH in the lumen of the intra-pyrenoid tubules. Future experimental studies will be important to investigate the trade-off between proton supply and CO_2_ leakage in employing thylakoid tubules traversing the pyrenoid matrix.

A key driver of the algal CCM is the pH difference maintained across different compartments. However, our model does not explicitly consider the reaction-diffusion kinetics of protons, but rather assumes a uniform pH in each compartment. Previous *in vivo* measurements of the pH biosensor pHluorin in Chlamydomonas suggest that the pyrenoid matrix has a relatively uniform basic pH (34), yet it remains unclear how the uniformity is achieved. Protons are produced and consumed in the reactions catalyzed by CA. In addition, Rubisco CO_2_ fixation yields two protons for every CO_2_ fixed (5). Our calculations suggest that the concentrations of free protons at measured physiological pH values are too low to account for the corresponding fluxes without creating noticeable pH gradients (*SI Appendix*, Sec. VI). Thus, efficient transport of protons must involve proton carriers. A recent modeling work suggests that two metabolites, RuBP and 3-PGA, could play an important role in buffering the pH of CO_2_-fixing Rubisco condensates (68) — these metabolites have pKa values of 6.7 and 6.5, respectively, and are present at millimolar concentrations in the pyrenoid and stroma (69). Understanding the molecular mechanisms underlying proton transport will be an important topic for future studies.

Based on the known molecular machinery of the Chlamydomonas CCM, we proposed a minimal set of engineering steps to install an effective CCM in plant chloroplasts (Fig. 6). In our favored engineering path, Rubisco first needs to be assembled into a CA-free pyrenoid-like condensate *in vivo*. A phase-separated pyrenoid matrix has been successfully reconstructed in Arabidopsis chloroplasts (56), but it remains to be verified whether the native stromal CA is excluded from the engineered pyrenoid. Then, the newly formed matrix needs to be positioned proximal to low-pH thylakoids, and the thylakoid CA needs to be localized to the pyrenoid-proximal thylakoid lumen. Recent work in Chlamydomonas has shown that a Rubisco-binding motif targets proteins to the pyrenoid, and appears to link the matrix to the intra-pyrenoid tubules and starch sheath (70), thus providing a framework for manipulating the organization of an engineered pyrenoid. For example, constructing a fusion protein of the plant thylakoid CA and a membrane protein containing the Rubisco-binding motif may promote the desired colocalization of the pyrenoid matrix, thylakoids, and thylakoid CA. The next step in our favored engineering path is to insert HCO_3_^−^ channels through the thylakoid membranes, which could double the CO_2_ fixation flux according to our calculation. Particularly important is targeting HCO_3_^−^ channels specifically to the thylakoid membranes, but not to the chloroplast membrane, since adding HCO_3_^−^ channels to the latter will lead to severe HCO_3_^−^ leakage out of the chloroplast. Native thylakoids may naturally form a CO_2_ diffusion barrier, which is expected to increase the performance of the CCM. Further studies of the pyrenoid starch sheath in Chlamydomonas will enable its reconstitution around the engineered pyrenoid, which we expect will maximize the efficacy and energetic efficiency of the CCM.

We hope that our model provides practical information for engineers aiming to transfer algal machinery into plants, and that it will serve as a useful quantitative tool to guide basic CCM studies in the future.

## Materials and Methods

See attached pdf.

## Acknowledgments

We thank members of the Jonikas and Wingreen groups for insightful discussions. This work was supported by the National Science Foundation, through grant MCB-1935444 and through the Center for the Physics of Biological Function (PHY-1734030). Schematics for a subset of figures were created with BioRender.com.

## Materials and Methods

### Reaction-diffusion model

To better understand the operation of the Chlamydomonas CO_2_-concentrating mechanism (CCM), we developed a multi-compartment reaction-diffusion model that takes into account the key CCM enzymes and transporters and the relevant architecture of the Chlamydomonas chloroplast (17). For simplicity, our model assumes spherical symmetry and considers a spherical chloroplast of radius *R*_chlor_ in an infinite cytosol. Thus, all model quantities can be expressed as functions of the radial distance *r* from the center of the chloroplast (Fig. 1B). The modeled chloroplast consists of three compartments: a spherical pyrenoid matrix of radius *R*_pyr_ (pH 8) in the center, surrounded by a stroma (pH 8), with thylakoids (luminal pH 6) traversing both the matrix and stroma (Fig. 1) (34–36). At steady state, flux-balance equations set the spatially-dependent concentrations of CO_2_, 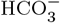, and H_2_CO_3_ in their respective compartments (indicated by subscripts; see *SI Appendix*, Sec. I and Table S1):

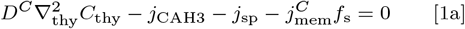

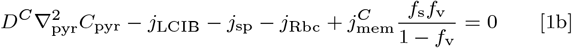

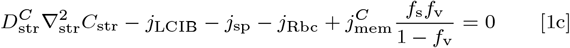

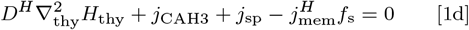

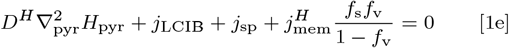

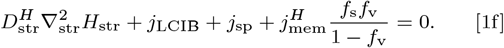

Here, *C* denotes the concentration of CO_2_, and *H* denotes the combined concentration of 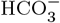 and H_2_CO_3_, which are assumed to be in fast equilibrium (43). Thus, their respective concentrations are given by 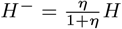 for 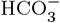 and 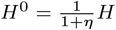 for H_2_CO_3_, where 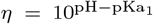 is a pH-dependent partition factor and pKa_1_ = 3.4 is the first pKa of H_2_CO_3_ (71). The first terms in Eqs. (1a-1f) describe the diffusive fluxes of inorganic carbon (Ci) within compartments. *D*^*C*^ and *D*^*H*^ denote, respectively, the diffusion coefficients of CO_2_, and 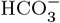 and H_2_CO_3_ combined, in aqueous solution. In a model with thylakoid stacks slowing Ci diffusion in the stroma, the effective diffusion coefficients 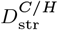 are obtained using a standard homogenization approach (see *SI Appendix*, Sec. IG and Fig. S5); 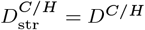 otherwise. The other flux terms (*j*_X_) in Eqs. (1a-1f) describe enzymatic reactions and inter-compartment Ci transport. Their expressions are provided in subsequent sections.

The boundary conditions at *r* = *R*_pyr_ are determined by the diffusive flux of Ci across the starch sheath at the matrix-stroma interface, i.e.,

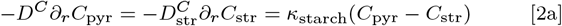

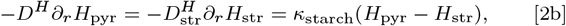

where the starch sheath is assumed to have the same permeability *k*_starch_ for all Ci species. *k*_starch_ → ∞ when there is no starch sheath and Ci can diffuse freely out of the matrix. *k*_starch_ = 0 describes an impermeable starch sheath (see *SI Appendix*, Sec. IF). Similarly, Ci transport flux across the chloroplast envelope yields the boundary conditions at *r* = *R*_chlor_, i.e.,

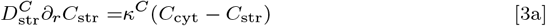

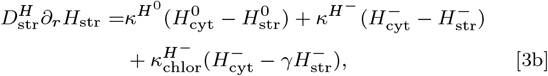

where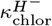 and *γ* denote the rate and reversibility of inward 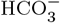 transport from the cytosol, representing the action of the uncharacterized chloroplast envelope 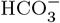 transporter LCIA (24, 26); *γ* = 1 corresponds to a passive bidirectional channel and *γ* < 1 corresponds to an active pump. The external CO_2_ conditions are specified by *C*_cyt_ and all cytosolic Ci species are assumed to be in equilibrium at pH 7.1 (see *SI Appendix*, Sec. VIB) (72). We set *C*_cyt_ = 10 μM for air-level CO_2_ conditions, and *C*_cyt_ = 1 μM for very low CO_2_ conditions.

### Thylakoid geometry

The thylakoid geometry has been characterized by cryo-electron tomography in Chlamydomonas (17). In our model, we account for this geometry by varying the local volume fraction *f*_v_ and surface-to-volume ratio *f*_s_ of the thylakoids. These fractions describe a tubule meshwork at the center of the pyrenoid (*r* ≤ *R*_mesh_), extended radially by *N*_tub_ cylindrical tubules each of radius *a*_tub_ (see *SI Appendix*, Sec. IC), i.e.,

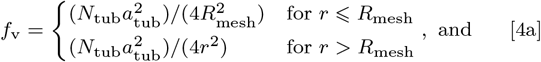

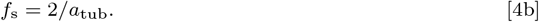

In the baseline model the thylakoid tubules are assumed to extend to the chloroplast envelope, i.e., the outer radius of tubules *R*_tub_ = *R*_chlor_. In a model with shorter tubules, we choose *R*_tub_ = 0.4 *R*_chlor_, and set *f*_v_ = 0 and *f*_s_ = 0 for *r* > *R*_tub_. Thus, the Laplace–Beltrami operator is given by 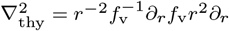 for the thylakoid tubules, and by 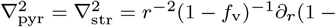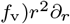 for the matrix and stroma.

### Enzyme kinetics

The model considers three key CCM enzymes, i.e., the carbonic anhydrases (CAs) CAH3 and LCIB and the CO_2_-fixing enzyme Rubisco. The interconversion between CO_2_ and 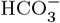 is catalyzed by both CAs and follows reversible Michaelis-Menten kinetics (73). The rate of CA-mediated 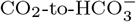 conversion is given by

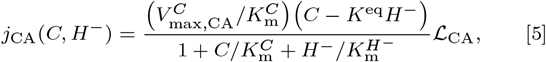

where 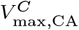 denotes the maximum rate of CA, 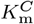 and 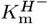 denote, respectively, the half-saturation concentrations for CO_2_ and 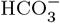, and 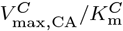 denotes the first-order rate constant which we refer to as the “rate” of the CA (Fig. 2). Finally, 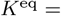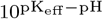 denotes the equilibrium ratio of CO_2_ to 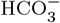, with the effective pKa pK_eff_ = 6.1 (42, 48). The localization function 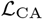 is equal to one for *r* where CA is present and zero elsewhere. The uncatalyzed spontaneous rate of 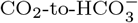 conversion, with a first-order rate constant 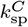, is given by 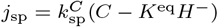 (74). Note that negative values of *j*_CA_ and *j*_sp_ denote fluxes of 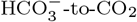 conversion.

The rate of CO_2_ fixation catalyzed by Rubisco is calculated from

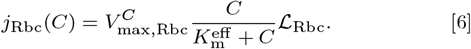

Here, 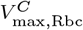 denotes the maximum rate, and the effective *K*_m_ (Rubisco *K*_m_ in Fig. 1) is given by 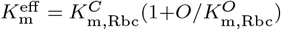 to account for competitive inhibition by O_2_ (75, 76), where *O* denotes the concentration of O_2_, and 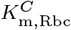 and 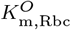 denote the half-saturation substrate concentrations for CO_2_ and O_2_, respectively. 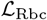 is equal to one where Rubisco is localized, and zero elsewhere.

In our baseline model, we assume that CAH3 is localized in the thylakoid tubules traversing the pyrenoid (33), LCIB is distributed diffusely in the stroma (46), and Rubisco is localized in the pyrenoid matrix (11). To explore the effect of enzyme localization, we vary the start and end radii of the enzymes while maintaining a constant number of molecules (see Figs. 4 and 5 and *SI Appendix*, Sec. V).

### Transport of Ci across thylakoid membranes

The flux of CO_2_ diffusing across the thylakoid membrane from the thylakoid lumen to the matrix or stroma is given by

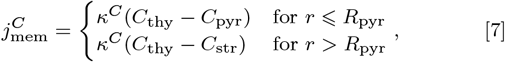

where *k^C^* denotes the permeability of thylakoid membranes to CO_2_. Similarly, the cross-membrane diffusive flux of 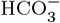 and H_2_CO_3_, 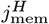, is given by

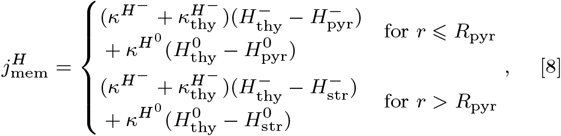

where 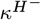 and 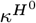 denote, respectively, the baseline membrane permeability to 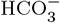 and H_2_CO_3_, and 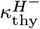 denotes the additional permeability of thylakoid membranes to 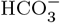 due to bestrophin-like channels (30). Note that the final terms of Eq. (1a) and Eqs. (1b,1c) differ by a factor of 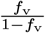 because the cross-membrane fluxes have a larger impact on the concentrations in the thylakoid compartment, which has a smaller volume fraction.

### Choice of parameters and numerical simulations

The model parameters are estimated from experiment (see *SI Appendix*, Table S1 and references therein), except for the rates of LCIB and CAH3 and the kinetic parameters of the 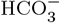 transporters, which are not known. We performed a systematic scan for these unknown parameters within a range of reasonable values (Fig. 2 and *SI Appendix*, Fig. S4). The numerical solutions of Eq. (1) were obtained by performing simulations using a finite element method. Partial differential equations were converted to their equivalent weak forms, computationally discretized by first-order elements (77), and implemented in the open-source computing platform FEniCS (78). A parameter sensitivity analysis was performed to verify the robustness of the model results (*SI Appendix*, Fig. S28). A convergence study was performed to ensure sufficient spatial discretization (*SI Appendix*, Fig. S29).

### Energetic cost of the CCM

We compute the energetic cost using the framework of nonequilibrium thermodynamics (54) (see *SI Appendix*, Sec. IIB, for details). In brief, the free-energy cost of any nonequilibrium process (reaction, diffusion, or transport) is given by (*j*_+_ − *j*_−_) ln(*j*_+_/*j*_−_) (in units of thermal energy *RT*), where *j*_+_ and *j*_−_ denote the forward and backward flux, respectively. Summing the energetic cost of nonequilibrium processes described in Eq. (1), we show that the total energy required to operate the CCM can be approximated (in units of *RT*) by

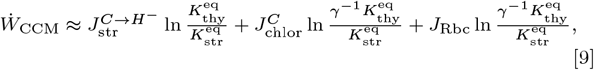

Here, 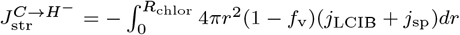 *dr* integrates the flux of LCIB-mediated and spontaneous conversion from CO_2_ to 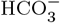 in the stroma, with 4π*r*^2^(1 − *f*_v_)*dr* being the geometric factor. 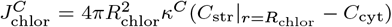 denotes the flux of CO_2_ diffusing from the stroma back out into the cytosol. 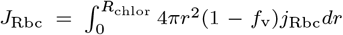 integrates the flux of CO_2_ fixation by Rubisco. The ln *γ*^−1^ and 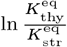 terms denote the free-energy cost of pumping 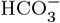 across the chloroplast envelope and pumping protons across the thylakoid membranes, respectively. Using ATP hydrolysis energy |Δ*G*_ATP_| = 51.5 *RT* (79), we compute the equivalent ATP spent per CO_2_ fixed as 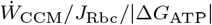.

### Well-mixed compartment model

To better understand the biophysical limit of the CCM, we consider a well-mixed compartment simplification of the full model. Specifically, we assume that (i) the diffusion of Ci is fast in the matrix and stroma, and therefore the concentrations of CO_2_ and 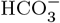 are constant across radii in each of the two compartments, taking values denoted by *C*_pyr_*, C*_str_,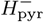, and 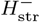; (ii) 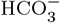 transport across the thylakoid membranes is fast, and thus the thylakoid tubule concentration of 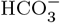 inside the pyrenoid is equal to 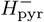, while the thylakoid tubule concentration outside the pyrenoid is equal to 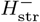; (iii) 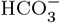 and CO_2_ are in equilibrium (catalyzed by CAH3) in the thylakoid tubules inside the pyrenoid, and thus the CO_2_ concentration therein is given by 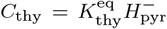; and (iv) the concentration of CO_2_ in the thylakoid tubules approaches *C*_str_ toward the chloroplast envelope. Thus, the flux-balance conditions are described by a set of algebraic equations of 4 variables, *C*_pyr_*, C*_thy_*, C*_str_ and 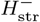 (see *SI Appendix*, Sec. III). The algebraic equations are solved using the Python-based computing library SciPy (80). The energetic cost of the well-mixed compartment model is computed similarly as above.

### Engineering paths

We are interested in how adding and removing individual components affects the overall functioning of the CCM. We thus measure the efficacy and energy efficiency of 216 CCM configurations, modulating the presence and localization of enzymes, 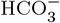 channels, and diffusion barriers. Each configuration is simulated using the reaction-diffusion model above with the appropriate parameters for that strategy (*SI Appendix*, Fig. S23).

To find all possible engineering paths between these configurations, we consider a graph on which each possible configuration is a node. Nodes are considered to be connected by an undirected edge if they are separated by one engineering step. Thus, by taking steps on the graph, we can search all possible engineering paths given a start node with poor CCM performance and a target node with good performance. A single engineering step could be the addition or removal of an enzyme, a channel, or a diffusion barrier, as well as the localization of a single enzyme. The exception is the localization of Rubisco, which we assume can exclude LCIB from the matrix as it forms a phase-separated condensate (55). We do not consider strategies employing both a starch sheath and thylakoid stacks as diffusion barriers. We use a custom depth-first search algorithm in MATLAB to identify all shortest engineering paths between a start and a target node.

### Data and software availability

All data and simulation code required to reproduce the results in this manuscript are available to the readers on GitHub: https://github.com/f-chenyi/Chlamydomonas-CCM.

