## Supplementary material for "Diffusion barriers and adaptive carbon uptake strategies enhance the modeled performance of the algal CO_2_-concentrating mechanism": SI Appendix

#### TABLE OF CONTENTS

List of SI Figures 2

List of SI Tables 2

I. A multi-compartment reaction-diffusion model of the Chlamydomonas CCM 4
- A. General formulation 4
- B. Enzyme kinetics 5
- C. Thylakoid geometry 6
- D. Diffusion and transport of inorganic carbon across the thylakoid membranes 7
- E. Diffusion and transport of inorganic carbon across the chloroplast envelope 7
- F. Starch sheath: a potential barrier to diffusion of inorganic carbon out of the pyrenoid matrix 8
- G. Thylakoid stacks: a potential barrier to slow diffusion of inorganic carbon in the stroma 8
- H. Model discussion: $CO_2$ leakage 9

II. Evaluating the performance of the CCM 10
- A. Normalized $CO_2$ fixation flux and net carbon fixation flux 10
- B. Energetic cost of the CCM 10
- C. A combined measure of CCM performance 14

III. Well-mixed compartment model with an impermeable starch sheath 17
- A. Governing equations 17
- B. Two extreme CCM strategies 18
- C. Feasible CCM strategies under varying $CO_2$ conditions 18
- D. Limits of the passive $CO_2$ uptake strategy 19

IV. Well-mixed compartment model with thylakoid stacks 20
- A. Diffusion length scale in the stroma 20
- B. Flux-balance equations 20
- C. Results and discussion 22

V. Localization and regulation of key CCM enzymes 24
- A. Localization and regulation of CAH3 24
- B. Localization of LCIB 24

VI. Supplementary discussion 26
- A. Proton transport 26
- B. Effect of different cytosolic Ci compositions 27
- C. Thylakoid morphologies 27
- D. Effect of LCIB-Rubisco complexes 28
- E. Import of bicarbonate into the cell 29

Supplementary figures and tables 31

References 63

**LIST OF SI FIGURES**


S1 $CO_2$ fixation by Rubisco is inhibited by oxygen at air-level $CO_2$. . . . . 31
S2 Volume fraction of the modeled chloroplast compartments. . . . . 31
S3 Low fixation efficiency at air-level $CO_2$ (10 μM cytosolic) in the absence of a diffusion barrier is due to severe $CO_2$ leakage from the pyrenoid matrix. . . . . 32
S4 $CO_2$ fixation flux increases with BST $HCO_3^-$ channel rate. . . . . 33
S5 Model for the effect of thylakoid stacks on diffusion in the stroma. . . . . 34
S6 Comparison between two types of diffusion barriers under air-level $CO_2$ (10 μM cytosolic). . . . . 35
S7 $CO_2$ concentrating performance is compromised by weaker barriers to the diffusion of inorganic carbon. 36
S8 A model including both thylakoid stacks and an impermeable starch sheath behaves similarly to a model with only the latter. . . . . 37
S9 In a modeled chloroplast employing thylakoid stacks and no starch sheath, LCIB activity increases intra-chloroplast $HCO_3^-$ concentration by converting external $CO_2$ but also effectively "steals" $CO_2$ from the pyrenoid matrix, leading to a nonmonotonic impact of LCIB rate on $CO_2$ concentration inside the pyrenoid matrix. . . . . 38
S10 A trade-off between $CO_2$ transport and $CO_2$ leakage leads to the existence of an intermediate membrane permeability to $CO_2$ that optimizes $CO_2$-concentrating performance. . . . . 39
S11 Localizing LCIB close to the chloroplast envelope prevents excessive $CO_2$ efflux drawn by LCIB from the pyrenoid matrix in a modeled chloroplast with thylakoid stacks slowing diffusion in the stroma and no starch sheath. . . . . 40
S12 Active $HCO_3^-$ transport across the chloroplast envelope enables an effective CCM under very low $CO_2$. 41
S13 Illustrations of the nonequilibrium processes that cost free energy. . . . . 42
S14 The least costly CCM strategies. . . . . 43
S15 A well-mixed compartment model of the CCM. . . . . 44
S16 Two extreme CCM strategies are compared under various $CO_2$ conditions for a model employing an impermeable starch sheath. . . . . 45
S17 Feasible CCM strategies depend on external $CO_2$ levels in the well-mixed compartment model. . . . . 46
S18 Feasible CCM strategies depend on external $CO_2$ levels in the full reaction-diffusion model. . . . . 47
S19 CAH3 localization and activity that enable an effective CCM depend on environmental $CO_2$ levels. . . . 49
S20 Localization of carbonic anhydrases enhances CCM performance under very low $CO_2$. . . . . 50
S21 A strong diffusion barrier limits the dependence of CCM activity on LCIB localization for a passive $CO_2$ uptake strategy. . . . . 51
S22 Localization of LCIB around the pyrenoid in a model with a starch sheath allows for $CO_2$ recycling without inducing carbon leakage from the chloroplast when the active $HCO_3^-$ pumping strategy is employed. . . . . 52
S23 Characterizing various CCM configurations yields feasible engineering paths for installing a CCM into plant chloroplast. . . . . 53
S24 Modeled cell growth rate is determined jointly by the efficacy and energy cost of the CCM. . . . . 54
S25 The composition of cytosolic Ci impacts the efficacy of Ci import strategies. . . . . 55
S26 Alternative thylakoid morphologies also support an effective CCM. . . . . 56
S27 Protons derived from Rubisco $CO_2$ fixation lead to a notable increase in local pH and $CO_2$ availability only in 10 μm CA-Rubisco condensates. . . . . 57
S28 Parameter sensitivity analysis of the multi-compartment reaction-diffusion model. . . . . 58
S29 Mesh convergence of the finite-element simulations. . . . . 59


**LIST OF SI TABLES**


S1 Summary of the notations and parameter values used in the reaction-diffusion model of the CCM. . . . . 60
S2 Summary of the notations and parameter values used in the combined metric calculation (Sec. II C). . . 61
S3 Summary of the parameters and their values used in the well-mixed compartment model. . . . . 62

S4 Efficacy and energetic efficiency of the 216 engineering configurations varying the presence and localization of enzymes, $HCO_3^-$ channels, and diffusion barriers. . . . . . 62

#### I. A MULTI-COMPARTMENT REACTION-DIFFUSION MODEL OF THE CHLAMYDOMONAS CCM

Chlamydomonas cells operate a $CO_2$-concentrating mechanism (CCM) to supply the central carbon-fixing enzyme, Rubisco, with a high concentration of its substrate, $CO_2$. To better understand how the Chlamydomonas CCM works, we build a multi-compartment reaction-diffusion model that consists of the essential mechanics of the CCM (see Fig. 1A), including the relevant architecture of the Chlamydomonas chloroplast. In brief, we model the chloroplast as a sphere of radius $R_{\rm chlor}$, comprised of three compartments — a spherical pyrenoid matrix of radius $R_{\rm pyr}$ in the center, surrounded by a stroma, with thylakoids traversing both matrix and stroma [1]. We assume a constant pH in each compartment, with the matrix and stroma at pH 8 [2] and the thylakoids at pH 6 [3]. In the model, we consider intra-compartment diffusion and inter-compartment exchange of inorganic carbon in the forms of $CO_2$, $H_2CO_3$, and $HCO_3^-$, as well as their interconversion catalyzed by CAH3 in the thylakoids and by LCIB in the stroma (see Figs. 1A and B). Carbonate ($CO_3^{2-}$) can be neglected at the pH values we consider [4].

Below we describe our model in full: in Sec. I A we derive the steady-state reaction-diffusion kinetics of inorganic carbon; in Sec. I B we describe the reaction kinetics of key CCM enzymes; in Sec. I C we specify the modeled geometry of the thylakoids; in Secs. I D and I E we detail the inter-compartment diffusion and transport of inorganic carbon; in Secs. I F and I G we introduce the presence of potential barriers to inorganic carbon diffusion; and in Sec. I H we discuss the issue of $CO_2$ leakage from the pyrenoid matrix. Notations and parameter values used in our model are provided in Table S1.

##### A. General formulation

We denote by $C$ the concentration of $CO_2$, by $H^-$ the concentration of $HCO_3^-$, by $H^0$ the concentration of $H_2CO_3$, and by $H$ the total concentration of $HCO_3^-$ and $H_2CO_3$, with respective compartments indicated by subscripts. For simplicity, our model assumes radial symmetry, i.e., concentrations of inorganic carbon are functions of radial coordinate $r$ only, $C = C(r)$ and $H = H(r)$. Thus, at steady state, the mass conservation of thylakoid $CO_2$ between $r$ and $r + dr$ is given by

$$\left[(4\pi r^2 f_{\rm v}) D^C \frac{\partial C_{\rm thy}}{\partial r}\right]_r^{r+dr} - (j_{\rm sp} + j_{\rm CAH3})(4\pi r^2 f_{\rm v} dr) - j^C_{\rm mem} ds(r, dr) = 0, \tag{S1}$$

where $f_{\rm v}(r)$ denotes the volume fraction of the thyalkoids, $ds(r, dr)$ denotes the surface area of thylakoid membrane between $r$ and $r+dr$, $D^C$ denotes the diffusion coefficient of $CO_2$, $j^C_{\rm mem}$ denotes the transport flux across the thylakoid membrane, $j_{\rm sp}$ denotes the flux of spontaneous conversion of $CO_2$ to $HCO_3^-$,and $j_{\rm CAH3}$ denotes the flux of conversion of $CO_2$ to $HCO_3^-$ catalyzed by CAH3. $j_{\rm mem}$ is positive for a flux from the thylakoid lumen into the matrix or stroma. Defining $f_{\rm s} \equiv \frac{ds(r,dr)}{4\pi r^2 f_{\rm v} dr}$, we obtain

$$D^C \frac{1}{r^2 f_{\rm v}} \frac{\partial}{\partial r}\left(r^2 f_{\rm v} \frac{\partial C_{\rm thy}}{\partial r}\right) - j_{\rm sp} - j_{\rm CAH3} - j^C_{\rm mem} f_{\rm s} = 0. \tag{S2}$$

We provide the expressions for $j_{\rm sp}$ and $j_{\rm CAH3}$ in Sec. I B, the expressions for $f_{\rm v}$ and $f_{\rm s}$ in Sec. I C, and the expression for $j^C_{\rm mem}$ in Sec. I D. Similarly, the steady-state flux-balance conditions for $CO_2$ in the pyrenoid matrix ($r \leqslant R_{\rm pyr}$) and in the stroma ($R_{\rm pyr} \leqslant r \leqslant R_{\rm chlor}$) yield, respectively,

$$D^C \frac{1}{(1-f_{\rm v})r^2} \frac{\partial}{\partial r}\left((1-f_{\rm v}) r^2 \frac{\partial C_{\rm pyr}}{\partial r}\right) - j_{\rm sp} - j_{\rm LCIB} - j_{\rm Rbc} + j^C_{\rm mem} \frac{f_{\rm s} f_{\rm v}}{1-f_{\rm v}} = 0, \text{ and} \tag{S3}$$

$$D^C_{\rm str} \frac{1}{(1-f_{\rm v})r^2} \frac{\partial}{\partial r}\left((1-f_{\rm v}) r^2 \frac{\partial C_{\rm str}}{\partial r}\right) - j_{\rm sp} - j_{\rm LCIB} - j_{\rm Rbc} + j^C_{\rm mem} \frac{f_{\rm s} f_{\rm v}}{1-f_{\rm v}} = 0, \tag{S4}$$

where $j_{\rm LCIB}$ denotes the flux of conversion of $CO_2$ to $HCO_3^-$ catalyzed by LCIB and $j_{\rm Rbc}$ denotes the flux of $CO_2$ fixation by Rubisco. Their expressions are given in Sec. I B. $D^C_{\rm str}$ denotes the diffusion coefficient of $CO_2$ in the stroma (see **Choice of parameters** below).

Steady-state flux-balance conditions for $HCO_3^-$ and $H_2CO_3$ in the thylakoids are given, respectively, by

$$D^{H^-} \frac{1}{r^2 f_{\rm v}} \frac{\partial}{\partial r}\left(r^2 f_{\rm v} \frac{\partial H^-_{\rm thy}}{\partial r}\right) + j_{\rm sp} + j_{\rm CAH3} - j^{H^-}_{\rm mem} f_{\rm s} + j^H_{\rm int} = 0, \text{ and} \tag{S5}$$

$$D^{H^0} \frac{1}{r^2 f_{\rm v}} \frac{\partial}{\partial r}\left(r^2 f_{\rm v} \frac{\partial H^0_{\rm thy}}{\partial r}\right) - j^{H^0}_{\rm mem} f_{\rm s} - j^H_{\rm int} = 0. \tag{S6}$$

Here, $D^{H^-}$ and $D^{H^0}$ denote the diffusion coefficients of $HCO_3^-$ and $H_2CO_3$, respectively. $j_{\rm int}^H$ denotes the flux of interconversion between $HCO_3^-$ and $H_2CO_3$. Since this reaction is very fast [4, 5], we set all the other terms to zero in Eqs. (S5) and (S6) and solve for the relation between $H^-$ and $H^0$, which yields $H^- = 10^{\rm pH-pKa_1} H^0 \equiv \eta H^0$, i.e., $H^- = \frac{\eta}{1+\eta} H$ and $H^0 = \frac{1}{1+\eta} H$, where $\rm pKa_1 = 3.4$ is the first pKa of $H_2CO_3$ and $\eta$ is a constant factor depending on the compartmental pH. Summing Eqs. (S5) and (S6) yields

$$D^H \frac{1}{r^2 f_{\rm v}} \frac{\partial}{\partial r}\left(r^2 f_{\rm v} \frac{\partial H_{\rm thy}}{\partial r}\right) + j_{\rm sp} + j_{\rm CAH3} - j_{\rm mem}^H = 0, \tag{S7}$$

where $D^H = \frac{\eta}{1+\eta} D^{H^-} + \frac{1}{1+\eta} D^{H^0}$ is the effective diffusion coefficient of the two species, and $j_{\rm mem}^H = j_{\rm mem}^{H^-} + j_{\rm mem}^{H^0}$ denotes combined transport flux whose expression is provided in Sec. I D. Likewise, we obtain the following steady-state flux-balance conditions in the matrix and stroma:

$$D^H \frac{1}{(1-f_{\rm v}) r^2} \frac{\partial}{\partial r}\left((1-f_{\rm v}) r^2 \frac{\partial H_{\rm pyr}}{\partial r}\right) + j_{\rm sp} + j_{\rm LCIB} + j_{\rm mem}^H \frac{f_{\rm s} f_{\rm v}}{1-f_{\rm v}} = 0, \text{ and} \tag{S8}$$

$$D_{\rm str}^H \frac{1}{(1-f_{\rm v}) r^2} \frac{\partial}{\partial r}\left((1-f_{\rm v}) r^2 \frac{\partial H_{\rm str}}{\partial r}\right) + j_{\rm sp} + j_{\rm LCIB} + j_{\rm mem}^H \frac{f_{\rm s} f_{\rm v}}{1-f_{\rm v}} = 0. \tag{S9}$$

Here, $D_{\rm str}^H$ denotes the diffusion coefficient of $HCO_3^-$ and $H_2CO_3$ in the stroma (see **Choice of parameters** below). Note that one can use the relations $H^- = \frac{\eta}{1+\eta} H$ and $H^0 = \frac{1}{1+\eta} H$ to express Eqs. (S7 - S9) as closed equations for $H$.

**Choice of parameters:** The sizes of the chloroplast and the pyrenoid matrix are estimated to be $R_{\rm chlor} = 3.42$ μm and $R_{\rm pyr} = 0.3 R_{\rm chlor}$ from the experimental images in [6]. Diffusion coefficients of $CO_2$ and $HCO_3^-$ in aqueous solution are measured to be $D^C = 1.88 \times 10^3$ μm$^2$/s [7] and $D^{H^-} = 1.15 \times 10^3$ μm$^2$/s [8]. We approximate the diffusion coefficient of $H_2CO_3$ by that of acetic acid due to their identical charge and similar size, i.e., $D^{H^0} \approx 1.2 \times 10^3$ μm$^2$/s [9]. Thus, for simplicity, we set $D^{H^0} = D^{H^-}$, which results in $D^H \equiv D^{H^-}$. In the presence of thylakoid stacks slowing inorganic carbon diffusion in the stroma, the effective diffusion coefficients $D_{\rm str}^{C/H} = D_{\rm eff}^{C/H}$ are computed in Sec. I G; $D_{\rm str}^{C/H} = D^{C/H}$ otherwise.

##### B. Enzyme kinetics

**Carbonic anhydrases:** The two carbonic anhydrases (CAs) CAH3 and LCIB catalyze the interconversion between $CO_2$ and $HCO_3^-$ [10, 11] and are known to be key to proper functioning of the CCM [12, 13]. Since the full reaction $CO_2 + H_2O \leftrightarrow HCO_3^- + H^+$ involves a proton, the equilibrium $CO_2$-to-$HCO_3^-$ ratio $K^{\rm eq} \equiv 10^{(\rm pK_{eff} - pH)}$ depends on the compartmental pH, where $\rm pK_{eff} = 6.1$ is the effective $\rm pK_a$, defined as the pH at which $CO_2$ and $HCO_3^-$ have equal equilibrium concentrations [4]. We assume the reactions catalyzed by CAs follow Michaelis–Menten kinetics, so that

$$j_{\rm CA}(C, H^-) = \frac{\left(V_{\rm max,CA}^C / K_{\rm m}^C\right)\left(C - K^{\rm eq} H^-\right)}{1 + C/K_{\rm m}^C + H^-/K_{\rm m}^{H^-}} \mathcal{L}_{\rm CA}, \tag{S10}$$

where $V_{\rm max,CA}^C$ denotes the maximum rate at which $CO_2$ is converted to $HCO_3^-$, $K_{\rm m}^C$ and $K_{\rm m}^{H^-}$ denote the half-saturation substrate concentrations for $CO_2$ and $HCO_3^-$, respectively, and $V_{\rm max,CA}^C / K_{\rm m}^C$ denotes the first-order rate constant which we refer to as the "rate" of that enzyme (see for example Fig. 2). Here, $\mathcal{L}_{\rm CA}$ denotes the localization of CAs (see **Enzyme localization** below). Note that the uncatalyzed spontaneous conversion of $CO_2$ to $HCO_3^-$ is much slower than the reactions catalyzed by CA. Nevertheless, we include it in our model for completeness, with a flux of $j_{\rm sp} = k_{\rm sp}^C (C - K^{\rm eq} H^-)$.

**Rubisco:** Rubisco not only catalyzes the reaction of $CO_2$ fixation but also catalyzes the oxygenation reaction in which $O_2$ competes with $CO_2$ for the active sites of Rubisco [14]. Thus, the flux of $CO_2$ fixation by Rubisco $j_{\rm Rbc}$ depends on the concentrations of both $CO_2$ and $O_2$. The Michaelis-Menten kinetics of competitive inhibition reads (Fig. S1) [14]

$$j_{\rm Rbc} = \frac{V_{\rm max,Rbc}^C C}{K_{\rm m,Rbc}^C (1 + O/K_{\rm m,Rbc}^O) + C} \mathcal{L}_{\rm Rbc}, \tag{S11}$$

where $V^C_{\text{max,Rbc}}$ denotes the maximum rate of $CO_2$ fixation, $O$ denotes the concentration of $O_2$, and $K^C_{\text{m,Rbc}}$ and $K^O_{\text{m,Rbc}}$ denote the half-saturation substrate concentrations for $CO_2$ and $O_2$, respectively. We term $K^C_{\text{m,Rbc}}(1 + \frac{O}{K^O_{\text{m,Rbc}}})$ the effective $K_\text{m}$ of $CO_2$ fixation (denoted by $K^{\text{eff}}_\text{m}$) or simply Rubisco $K_\text{m}$ (see Figs. 1 and S1). Similar to above, $\mathcal{L}_{\text{Rbc}}$ denotes the localization of Rubisco (see **Enzyme localization** below).

**Enzyme localization:** In our spherically symmetric model, an enzyme ranging from $R_\text{s}$ to $R_\text{e}$ is described by a localization function $\mathcal{L}_{\text{enzyme}} = \ell(r; R_\text{s}, R_\text{e}) \equiv \Theta(r - R_\text{s})\Theta(R_\text{e} - r)$ which is equal to one between its start radius $R_\text{s}$ and its end radius $R_\text{e}$ and zero elsewhere. Here, $\Theta(x)$ denotes the Heaviside step function. Under air-level $CO_2$ (10 μM cytosolic), Rubisco is localized to the pyrenoid matrix [15, 16], CAH3 is localized in the thylakoid tubules traversing the matrix [17], and LCIB is diffusely distributed in the stroma [18]. Thus, for the baseline model, we set $\mathcal{L}_{\text{Rbc}} = \ell(r; 0, R_{\text{pyr}})$, $\mathcal{L}_{\text{CAH3}} = \ell(r; 0, R_{\text{pyr}})$ and $\mathcal{L}_{\text{LCIB}} = \ell(r; R_{\text{pyr}}, R_{\text{chlor}})$. To explore the effect of enzyme localization on CCM performance (see Sec. V), we vary the start and end radii of the enzymes while maintaining a constant number of molecules.

**Choice of parameters:** For both CAH3 and LCIB, we estimate $K^C_\text{m} = K^{H^-}_\text{m} = 5$ mM as the values for a typical CA [19, 20]. The first-order rate constant of CAH3 can be estimated from experiments [21] to be $10^4$ s$^{-1}$. To our knowledge, the enzyme kinetics of LCIB have not been measured. Thus, we vary the $V^C_{\text{max}}$ of LCIB in the model. The rate constant for spontaneous conversion of $CO_2$ to $HCO_3^-$ is set to be $k^C_{\text{sp}} = 0.036$ s$^{-1}$ [22].

For Rubisco, the half-saturation substrate concentrations are measured to be $K^C_{\text{m,Rbc}} = 30$ μM and $K^O_{\text{m,Rbc}} = 150$ μM [23, 24]. The oxygen concentration is assumed to be the same as that in aqueous solution in contact with air, $O = 230$ μM [25], since oxygen evolution only minimally affects the concentration of oxygen in the chloroplast. $V^C_{\text{max,Rbc}}$ can be estimated either from the measured $k_{\text{cat}}$ [23] and concentration [26] of Rubisco or from saturating oxygen evolution measurements [16]. Both yield similar estimates of $V^C_{\text{max,Rbc}} = 15$ mM/s.

##### C. Thylakoid geometry

*In vivo*, the thylakoids enter the pyrenoid matrix through gaps in the starch sheath in the form of roughly cylindrical membrane tubules, and as they reach the center of the matrix (when $r \leqslant R_{\text{mesh}} \approx 0.4$ μm) they become connected and form a reticulated meshwork [1]. These two geometries are modeled as follows:

**Cylindrical tubules:** For $r > R_{\text{mesh}}$, we approximate the geometry of the thylakoids as cylindrical tubules exntending in the radial direction. We consider $N_{\text{tub}}$ tubules, each having a constant radius of $a_{\text{tub}}$. The volume fraction of these tubules at $r$ is given by $f_\text{v} = \frac{N_{\text{tub}}\pi a^2_{\text{tub}} dr}{4\pi r^2 dr} = \frac{N_{\text{tub}} a^2_{\text{tub}}}{4r^2}$. The infinitesimal surface area at radius $r$ is given by $ds(r, dr) = N_{\text{tub}}(2\pi a_{\text{tub}})dr$, and we define $f_\text{s} \equiv \frac{ds(r,dr)}{4\pi r^2 f_\text{v} dr} = \frac{2}{a_{\text{tub}}}$. We employ these functions $f_\text{v}$ and $f_\text{s}$ in our radially symmetric model for the Chlamydomonas CCM.

**Reticulated meshwork:** For $r \leqslant R_{\text{mesh}}$, we approximate the geometry of the thylakoids as a reticulated meshwork. To establish the necessary parameters for our radially symmetric model, we consider a cubic volume of $L^3$ containing a cubic mesh of thin tubules whose radius $a_{\text{tub}}$ is much smaller than the intertubule mesh size $\Delta L$. The volume fraction of the tubules in the meshwork can be estimated by $f_\text{v} \approx 3\big(\frac{L^2}{\Delta L^2}\big)\pi a^2_{\text{tub}} L/L^3 = 3\pi a^2_{\text{tub}}/(\Delta L)^2$, which is constant. The total surface area of tubules in the meshwork can be calculated as $3\big(\frac{L^2}{\Delta L^2}\big)2\pi a_{\text{tub}} L$. The ratio of this total surface area to the total volume of tubules in the meshwork $f_\text{v} L^3$ yields $f_\text{s} = \frac{2}{a_{\text{tub}}}$.

Taken together, the geometric factors in Eqs. (S2 - S4) and (S7 - S9) are given by (Fig. S2)

$$f_\text{v}(r) = \begin{cases} \frac{N_{\text{tub}} a^2_{\text{tub}}}{4R^2_{\text{mesh}}} & r \leqslant R_{\text{mesh}} \\ \frac{N_{\text{tub}} a^2_{\text{tub}}}{4r^2} & r > R_{\text{mesh}} \end{cases}, \text{ and } f_\text{s}(r) \equiv \frac{2}{a_{\text{tub}}}. \tag{S12}$$

**Choice of parameters:** For simplicity, in the baseline model we assume that the thylakoid tubules extend to the radius of the chloroplast $R_{\text{chlor}}$. When tubules extend to a radius $R_{\text{tub}}$ that is smaller than $R_{\text{chlor}}$, we set $f_\text{v} = 0$ and $f_\text{s} = 0$ for $r > R_{\text{tub}}$. Tubule geometric parameters are estimated from experimental images in [1] to be $N_{\text{tub}} = 40$, and $a_{\text{tub}} = 0.05$ μm.

##### D. Diffusion and transport of inorganic carbon across the thylakoid membranes

The diffusive flux of $CO_2$ across the thylakoid membrane from the thylakoid lumen to the matrix or stroma is calculated from

$$j^C_{\text{mem}} = \begin{cases} \kappa^C(C_{\text{thy}} - C_{\text{pyr}}) & \text{for } r < R_{\text{pyr}}, \text{ and} \\ \kappa^C(C_{\text{thy}} - C_{\text{str}}) & \text{for } r \geqslant R_{\text{pyr}}, \end{cases} \tag{S13}$$

where $\kappa^C$ denotes the permeability of thylakoid membranes to $CO_2$. The calculation of the diffusion flux takes the same form for $H_2CO_3$ and $HCO_3^-$ with permeability coefficients $\kappa^{H^0}$ and $\kappa^{H^-}$, respectively. In addition, Chlamydomonas cells possess thylakoid localized bestrophin-like anion channels that are essential to the CCM [27, 28]. These proteins are the prime candidates for $HCO_3^-$ channels to facilitate $HCO_3^-$ diffusion across thylakoid membranes. Thus, we obtain the cross-membrane exchange flux of $H_2CO_3$ and $HCO_3^-$ to be

$$j^H_{\text{mem}} = \begin{cases} \kappa^{H^0}\left(\frac{1}{1+\eta_{\text{thy}}}H_{\text{thy}} - \frac{1}{1+\eta_{\text{pyr}}}H_{\text{pyr}}\right) + (\kappa^{H^-} + \kappa^{H^-}_{\text{thy}})\left(\frac{\eta_{\text{thy}}}{1+\eta_{\text{thy}}}H_{\text{thy}} - \frac{\eta_{\text{pyr}}}{1+\eta_{\text{pyr}}}H_{\text{pyr}}\right) & \text{for } r < R_{\text{pyr}} \\ \kappa^{H^0}\left(\frac{1}{1+\eta_{\text{thy}}}H_{\text{thy}} - \frac{1}{1+\eta_{\text{str}}}H_{\text{str}}\right) + (\kappa^{H^-} + \kappa^{H^-}_{\text{thy}})\left(\frac{\eta_{\text{thy}}}{1+\eta_{\text{thy}}}H_{\text{thy}} - \frac{\eta_{\text{str}}}{1+\eta_{\text{str}}}H_{\text{str}}\right) & \text{for } r \geqslant R_{\text{pyr}} \end{cases}, \tag{S14}$$

where $\kappa^{H^-}_{\text{thy}}$ denotes the additional permeability of thylakoid membranes to $HCO_3^-$ due to bestrophin-like channels.

**Choice of parameters:** We estimate $\kappa^C = 300$ μm/s and $\kappa^{H^-} = 0.05$ μm/s from previous measurements in diatoms [29]. We estimate $\kappa^{H^0} = 30$ μm/s assuming that $H_2CO_3$ has the same membrane permeability as formic acid ($H_2CO_2$) due to their identical charge and similar size [4, 30]. The bestrophin-like channels are not well characterized. Thus, we treat $\kappa^{H^-}_{\text{thy}}$ as a variable. Note that we have assumed that these channels only allow the passage of $HCO_3^-$ but not $CO_2$ and $H_2CO_3$. Such charge selectivity can presumably be achieved through electrostatic interactions of $HCO_3^-$ with the channel [31, 32].

##### E. Diffusion and transport of inorganic carbon across the chloroplast envelope

We assume that the modeled chloroplast sits in an effectively infinite sea of cytosol which serves as a reservoir of inorganic carbon. For simplicity, we assume that the ratio of $HCO_3^-$ concentration (denoted by $H^-_{\text{cyt}}$) to $CO_2$ concentration (denoted by $C_{\text{cyt}}$) is roughly 10 in the cytosol, which corresponds to the equilibrium at pH 7.1, close to the measured cytosolic pH [33].

$CO_2$ enters the chloroplast by diffusing passively through the chloroplast envelope, which sets the boundary condition

$$D^C_{\text{str}} \frac{\partial C_{\text{str}}}{\partial r}\bigg|_{r=R_{\text{chlor}}} = \kappa^C(C_{\text{cyt}} - C_{\text{str}}|_{r=R_{\text{chlor}}}), \tag{S15}$$

where we have assumed that the permeability $\kappa^C$ of the chloroplast membrane to $CO_2$ is the same as that of the thylakoid membranes.

Similarly, both $H_2CO_3$ and $HCO_3^-$ can enter the chloroplast via passive diffusion with permeability coefficients $\kappa^{H^-}$ and $\kappa^{H^0}$, respectively. In addition, Chlamydomonas cells can employ an active $HCO_3^-$ uptake system, which consists of an ATP-binding cassette (ABC) transporter HLA3 at the plasma membrane and a formate/nitrite-transporter (FNT) LCIA at the chloroplast membrane to uptake $HCO_3^-$ [34]. In our model, we introduce a chloroplast envelope $HCO_3^-$ transporter with a rate $\kappa^{H^-}_{\text{chlor}}$ and a reversibility $\gamma$ of inward $HCO_3^-$ transport to represent the action of LCIA. Thus, the total influx of $H_2CO_3$ and $HCO_3^-$ across the chloroplast envelope is

$$\begin{aligned} &\kappa^{H^0}\left(\frac{1}{1+\eta_{\text{cyt}}}H_{\text{cyt}} - \frac{1}{1+\eta_{\text{str}}}H_{\text{str}}|_{r=R_{\text{chlor}}}\right) + \kappa^{H^-}\left(\frac{\eta_{\text{cyt}}}{1+\eta_{\text{cyt}}}H_{\text{cyt}} - \frac{\eta_{\text{str}}}{1+\eta_{\text{str}}}H_{\text{str}}|_{r=R_{\text{chlor}}}\right) \\ &+ \kappa^{H^-}_{\text{chlor}}\left(\frac{\eta_{\text{cyt}}}{1+\eta_{\text{cyt}}}H_{\text{cyt}} - \gamma\frac{\eta_{\text{str}}}{1+\eta_{\text{str}}}H_{\text{str}}|_{r=R_{\text{chlor}}}\right) \equiv D^H_{\text{str}}\frac{\partial H_{\text{str}}}{\partial r}\bigg|_{r=R_{\text{chlor}}}. \end{aligned} \tag{S16}$$

**Choice of parameters:** The environmental $CO_2$ condition is specified by $C_{\text{cyt}}$ in our model. In particular, we consider air-level $CO_2$ (400 ppm) which gives us $C_{\text{cyt}} = 10$ μM and very low $CO_2$ under which $C_{\text{cyt}} = 1$ μM [34]. For the chloroplast envelope $HCO_3^-$ transporter, we vary its rate $\kappa^{H^-}_{\text{chlor}}$ and reversibility $\gamma$ to explore possible CCM performances (Fig. 3). Note that $\gamma = 0$ corresponds to completely unidirectional pumping while $\gamma = 1$ corresponds to passive channel transport.

##### F. Starch sheath: a potential barrier to diffusion of inorganic carbon out of the pyrenoid matrix

In Chlamydomonas, the conditions that induce the CCM also induce the formation of a starch sheath around the pyrenoid matrix [35]. This correlation has led to the speculation that the starch sheath could act as a barrier to slow the efflux of $CO_2$ (and other forms of inorganic carbon) from the pyrenoid matrix [35]. Here, we model the starch sheath as a thin membrane between the matrix and the stroma, with a permeability $\kappa_{\text{starch}}$ to all forms of inorganic carbon. Thus, the boundary conditions at the pyrenoid radius $R_{\text{pyr}}$ are given by

$$-D^C \frac{\partial C_{\text{pyr}}}{\partial r}\Big|_{r=R_{\text{pyr}}} = -D^C_{\text{str}} \frac{\partial C_{\text{str}}}{\partial r}\Big|_{r=R_{\text{pyr}}} = \kappa_{\text{starch}}\big(C_{\text{pyr}}|_{r=R_{\text{pyr}}} - C_{\text{str}}|_{r=R_{\text{pyr}}}\big), \text{ and} \tag{S17a}$$

$$-D^H \frac{\partial H_{\text{pyr}}}{\partial r}\Big|_{r=R_{\text{pyr}}} = -D^H_{\text{str}} \frac{\partial H_{\text{str}}}{\partial r}\Big|_{r=R_{\text{pyr}}} = \kappa_{\text{starch}}\big(H_{\text{pyr}}|_{r=R_{\text{pyr}}} - H_{\text{str}}|_{r=R_{\text{pyr}}}\big). \tag{S17b}$$

**Choice of parameters:** Note that $\kappa_{\text{starch}} \to \infty$ corresponds to the case where the starch sheath plays no role in slowing the diffusion of inorganic carbon out of the pyrenoid matrix, while $\kappa_{\text{starch}} \to 0$ corresponds to the case where the starch sheath is completely impermeable to inorganic carbon, which can then only enter or leave the pyrenoid via the thylakoid tubules. In the absence of other types of diffusion barriers, we term the limit $\kappa_{\text{starch}} \to \infty$ "no diffusion barrier", and the limit $\kappa_{\text{starch}} \to 0$ "starch sheath" (see Figs. 2 and S6). The performance of the CCM for finite starch permeabilites between these two limits is shown in Fig. S7. Note that for $\kappa_{\text{starch}} \lesssim \kappa^C = 3 \times 10^{-4}$ m/s, $CO_2$ leakage from the pyrenoid occurs primarily via thylakoid tubules, and in this regime the $CO_2$ concentration in the pyrenoid is higher than the Rubisco $K_{\text{m}}$ (Fig. S7D). Thus, a starch sheath with a $CO_2$ permeability comparable to that of a typical membrane can be as effective a barrier as an impermeable starch sheath.

##### G. Thylakoid stacks: a potential barrier to slow diffusion of inorganic carbon in the stroma

Despite the proposed role of the starch sheath as a diffusion barrier preventing $CO_2$ efflux, mutants that are unable to synthesize starch still appear to have a fully functional CCM at air-level $CO_2$ (10 μM cytosolic) [36, 37]. According to our model (see Fig. 2), however, the CCM should be compromised in the absence of a barrier to $CO_2$ diffusion due to $CO_2$ leakage from the pyrenoid (see Sec. I H and Fig. S3). To resolve this discrepancy, we speculate that the thylakoid stacks, which comprise layers of membranes surrounding the pyrenoid, could form another diffusion barrier to slow $CO_2$ (as well $HCO_3^-$ and $H_2CO_3$) efflux.

For simplicity, we model the stroma packed with thylakoid stacks as a homogeneous compartment in which the diffusion of inorganic carbon is slowed. The effective diffusion coefficient $D_{\text{eff}}$ depends on the geometry of the thylakoid stacks and the rate $\kappa$ at which inorganic carbon species diffuse across thylakoid membranes. We model a realistic geometry of the thylakoid stacks *in silico* as follows (Fig. S5): Thylakoids of luminal width $d_h$ and membrane thickness $d_t$ are stacked laterally with alternating large and small interthylakoid stroma spaces $d_{\text{l}}$ and $d_{\text{s}}$. Thylakoid sheets possess wide and narrow gaps $\Delta_{\text{w}}$ and $\Delta_{\text{n}}$ through which inorganic carbon molecules can diffuse freely. The gaps are center-aligned and have a longitudinal repeat length of $L_{\text{rep}}$. Given the diffusion coefficient $D$ in the thylakoid lumen and interthylakoid stroma space and the diffusion coefficient $D_{\text{mem}} = \kappa d_t$ through the thylakoid membranes, we numerically simulate 1D lateral diffusion across the *in silico* thylakoid stacks and obtain $D_{\text{eff}}(D, D_{\text{mem}}) = D\phi(D_{\text{mem}}/D)$ where $\phi$ is the dimensionless calibration function shown in Fig. S5.

In a model where we consider the thylakoid stacks as a diffusion barrier, we replace the diffusion coefficients in Eq. (S4, S9, S15, and S16) with $D^C_{\text{eff}} = D^C\phi(\frac{\kappa^C}{D^C/d_t})$ for $CO_2$ and with $D^H_{\text{eff}} = \frac{1}{1+\eta_{\text{str}}}D^{H^0}_{\text{eff}} + \frac{\eta_{\text{str}}}{1+\eta_{\text{str}}}D^{H^-}_{\text{eff}} \approx D^{H^-}\phi(\frac{\kappa^{H^-}+\kappa^{H^-}_{\text{thy}}}{D^{H^-}/d_t})$ for $HCO_3^-$ and $H_2CO_3$.

**Choice of parameters:** We analyzed cryo-electron tomography images from [1, 38] and measured the following geometric parameters: $d_t = 5$ nm, $d_h = 10$ nm, $d_{\text{s}} = 4$ nm, $d_{\text{l}} = 40$ nm, $\Delta_{\text{n}} = 5.6$ nm, $\Delta_{\text{w}} = 50$ nm, and $L_{\text{rep}} = 800$ nm. In addition, we estimated the fraction of narrow gaps to be 40%. Note that the CCM performance for a model with both thylakoid stacks and an impermeable starch sheath is very similar to that of a model with only an impermeable starch sheath (Fig. S8). In the absence of a starch sheath, the model dependence of CCM performance on the strength of the diffusion barrier formed by the thylakoid stacks is shown in Fig. S7.

##### H. Model discussion: $CO_2$ leakage

To illustrate the important issue of $CO_2$ leakage from the pyrenoid matrix in the absence of a diffusion barrier, we consider a simplified problem where $CO_2$ is produced at a flux of $Q_i$ in the center of a sphere of radius $R$ packed with Rubisco. $CO_2$ can either diffuse away or be consumed by Rubsico via linear kinetics with a first-order rate coefficient $\mu$.

**Reaction-diffusion kinetics.** Assume that the diffusion coefficients of $CO_2$ molecules inside and outside the sphere are given by $D_{\text{in}}$ and $D_{\text{out}}$, respectively. The reaction-diffusion kinetics can be described by

$$\begin{cases} D_{\text{in}}\frac{1}{r^2}\frac{\partial}{\partial r}\left(r^2\frac{\partial C}{\partial r}\right) - \mu c = 0 & \text{for } r \leqslant R, \text{ and} \\ D_{\text{out}}\frac{1}{r^2}\frac{\partial}{\partial r}\left(r^2\frac{\partial C}{\partial r}\right) = 0 & \text{for } r > R. \end{cases} \tag{S18}$$

The boundary conditions (B.C.s) are given by ① $C|_{r\to\infty} = C_\infty$, ② $C|_{r=R^+} = C|_{r=R^-}$, ③ $D_{\text{out}}\frac{\partial C}{\partial r}\big|_{r=R^+} = D_{\text{in}}\frac{\partial C}{\partial r}\big|_{r=R^-}$, and ④ $-4\pi r^2 D_{\text{in}}\frac{\partial C}{\partial r}\big|_{r=0} = Q_i$. Here, $C_\infty$ denotes the concentration of $CO_2$ at infinity. Using B.C. ①, we obtain the solution for $r > R$, $C(r) = \mathcal{K}(R/r) + C_\infty$, where $\mathcal{K}$ is an undetermined constant. The general solution for $r \leqslant R$ can be written as $C = \mathcal{A}\sinh(\frac{r}{R_\mu})(R/r) + \mathcal{B}\cosh(\frac{r}{R_\mu})(R/r)$ where $\mathcal{A}$ and $\mathcal{B}$ are two constants, and $R_\mu = (D_{\text{in}}/\mu)^{1/2}$ denotes the typical length over which a $CO_2$ molecule can diffuse before being consumed by Rubisco. Next, we determine $\mathcal{A}, \mathcal{B}, \mathcal{K}$ using B.C.s ② - ④, which yield

$$② \Rightarrow \mathcal{A}\sinh(\tilde{R}) + \mathcal{B}\cosh(\tilde{R}) = c_\infty + \mathcal{K}, \tag{S19}$$

$$③ \Rightarrow \mathcal{A}\big[\cosh(\tilde{R})\tilde{R} - \sinh(\tilde{R})\big] + \mathcal{B}\big[\sinh(\tilde{R})\tilde{R} - \cosh(\tilde{R})\big] = -\epsilon_D\mathcal{K}, \text{ and} \tag{S20}$$

$$④ \Rightarrow \mathcal{B} = \frac{Q_i}{4\pi D_{\text{in}}R}, \tag{S21}$$

where we have defined two dimensionless variable $\epsilon_D = D_{\text{out}}/D_{\text{in}}$ and $\tilde{R} = R/R_\mu$. Equations (S19) and (S20) yield

$$\mathcal{A} = \frac{C_\infty\epsilon_D + \mathcal{B}\big[(1-\epsilon_D)\cosh(\tilde{R}) - \tilde{R}\sinh(\tilde{R})\big]}{\tilde{R}\cosh(\tilde{R}) - (1-\epsilon_D)\sinh(\tilde{R})}. \tag{S22}$$

**Percent leakage.** Within the simple model, how much $CO_2$, out of all the $CO_2$ produced at the center $r = 0$, diffuses away to infinity without being fixed by Rubisco? We compute the total leakage flux as $Q_R = -4\pi R^2 D_{\text{out}}\frac{\partial C}{\partial r}\big|_{r=R^+} = 4\pi D_{\text{out}}R\mathcal{K} = 4\pi D_{\text{out}}R\left[\mathcal{A}\sinh(\tilde{R}) + \mathcal{B}\cosh(\tilde{R}) - C_\infty\right]$. Introducing Eqs. (S21) and (S22) into the expression for $Q_R$, we obtain, in the low $CO_2$ limit $C_\infty \to 0$,

$$\frac{Q_R}{Q_i} \approx \frac{\epsilon_D\tilde{R}}{\tilde{R}\cosh(\tilde{R}) - (1-\epsilon_D)\sinh(\tilde{R})}. \tag{S23}$$

When the inside and outside have equal diffusivity, i.e., when $\epsilon_D = 1$, the right side of Eq. (S23) becomes $[\cosh(\tilde{R})]^{-1}$. When the radius $R$ of the sphere of Rubisco is much smaller then the typical capture length $R_\mu$, $\tilde{R} \ll 1$ and $Q_R/Q_i \approx 1$, i.e., almost all $CO_2$ diffuses away to infinity without being fixed by Rubisco. When $R \gg R_\mu$, $Q_R/Q_i$ approaches zero as $\frac{1}{2}\exp(-\tilde{R})$ and $CO_2$ leakage is minimal. Choosing parameters relavant to the Chlamydomonas pyrenoid, $R = R_{\text{pyr}} \approx 1$ µm, $D_{\text{in}} = D^C = 1.88 \times 10^3$ µm$^2$/s and $\mu \approx 200$/s (Table S1), yields $\tilde{R} \approx 0.326$. Thus, as shown in Fig. S3A, the $CO_2$ leakage percentage is 95% when there is no $CO_2$ diffusion barrier, i.e., when $\epsilon_D = 1$, and it decreases as $\epsilon_D$ drops, e.g., in the presence of thylakoid stacks acting to slow diffusion in the stroma.

Next, we assume $\epsilon_D = 1$ and consider how the percent $CO_2$ leakage varies with the size of the sphere filled with Rubisco. If the concentration of Rubisco is fixed, i.e., $\mu$ does not change with $R$, then the percent leakage decreases with increasing $R$ (Fig. S3B) — since $R_\mu$ is unchanged, $\tilde{R}$, and thus the probability of $CO_2$ being fixed, increases with $R$ when there is more Rubiso. However, if the total number of Rubiscos is fixed, then $\mu \propto R^{-3}$ and therefore $\tilde{R} \propto R^{-1/2}$, so that the percent leakage actually decreases as $R$ becomes smaller (Fig. S3C), indicating that concentrating Rubisco in the pyrenoid matrix could help reduce $CO_2$ leakage.

#### II. EVALUATING THE PERFORMANCE OF THE CCM

In this section, we discuss how to evaluate the functionality of the CCM. In Sec. II A we consider the ability to fix $CO_2$; in Sec. II B we consider the energetic cost to operate the CCM; and in Sec. II C we attempt to integrate both aspects into a combined measure of CCM performance. To simplify the notation, we denote by

$$I\big[R_\mathrm{s}, R_\mathrm{e}, f(r), g(r)\big] \equiv \int_{R_\mathrm{s}}^{R_\mathrm{e}} f(r) 4\pi r^2 g(r) dr \tag{S24}$$

the integral of an arbitrary function $f(r)$ from the starting radius $R_\mathrm{s}$ to the end radius $R_\mathrm{e}$ in spherical coordinates, where $g(r)$ denotes the geometric factor explicitly.

##### A. Normalized $CO_2$ fixation flux and net carbon fixation flux

The ability of the chloroplast to fix $CO_2$ can be quantified by comparing the $CO_2$ fixation flux through Rubisco to the maximal flux if Rubisco were fully saturated, i.e.,

$$\Phi_C \equiv \frac{I[0, R_\mathrm{chlor}, j_\mathrm{Rbc}, 1-f_\mathrm{v}]}{V^C_\mathrm{max,Rbc} I[0, R_\mathrm{chlor}, \mathcal{L}_\mathrm{Rbc}, 1-f_\mathrm{v}]}, \tag{S25}$$

which we term the Normalized $CO_2$ Fixation Flux (henceforth, NCFF). As a criterion for a "working" CCM, we require NCFF $> 0.5$, i.e., local $CO_2$ concentration around Rubisco on average higher than the Rubisco $K_\mathrm{m}$. When Rubisco becomes fully saturated, NCFF approaches 1.

As alluded to in Sec. I B, Rubisco also has oxygenase activity. The oxygenation flux can be calculated as

$$J_\mathrm{oxy} = I[0, R_\mathrm{chlor}, j_\mathrm{oxy}, 1-f_\mathrm{v}], \tag{S26}$$

where the oxygenation reaction is described by the Michaelis-Menten kinetics

$$j_\mathrm{oxy} = \frac{V^O_\mathrm{max,Rbc} O}{O + K^O_\mathrm{m,Rbc}(1 + \frac{C}{K^C_\mathrm{m,Rbc}})} \mathcal{L}_\mathrm{Rbc}. \tag{S27}$$

For every two oxygenation reactions, a carbon is lost in the form of $CO_2$ [39]. Thus, the net carbon fixation flux compared to its maximum is given by

$$\Psi_C \equiv \Phi_C - \Phi_O, \text{ where } \Phi_O \equiv \frac{1}{2}\left(\frac{J_\mathrm{oxy}}{V^C_\mathrm{max,Rbc} I[0, R_\mathrm{chlor}, \mathcal{L}_\mathrm{Rbc}, 1-f_\mathrm{v}]}\right). \tag{S28}$$

We term $\Psi_C$ the Normalized Net Carbon Fixation Flux (henceforce, NNCFF). This metric will be used in Sec. II C when considering net carbon biomass growth.

##### B. Energetic cost of the CCM

**General formula.** We follow the theoretical framework of nonequilibrium thermodynamics [40] to calculate the energetic cost of the CCM. For a biochemical reaction out of equilibrium, the free-energy dissipation rate (per volume) can be calculated from [41–44]

$$\Delta \dot{W}_\mathrm{reaction} = (j_+ - j_-) \times \Big(RT \ln \frac{j_+}{j_-}\Big) \geqslant 0, \tag{S29}$$

where $j_+$ and $j_-$ denote, respectively, the forward and backward reaction fluxes (in units of molar concentration per time), $(j_+ - j_-)$ is the net reaction flux, and $RT \ln \frac{j_+}{j_-}$ is the chemical potential difference between the reactants and products. Equation (S29) is equivalent to Crook's fluctuation theorem at nonequilibrium steady state, and can be applied to compute the free-energy dissipation for a wide range of nonequilibrium processes. For example, given a nonuniform concentration profile $c(r)$ in the radial direction, Eq. (S29) can be generalized to calculate the dissipation rate associated with the diffusion flux [40], which yields

$$\Delta \dot{W}_\mathrm{diffusion} = \Big(D\frac{\partial c}{\partial r}\Big) \times \Big(RT \frac{\partial \ln c}{\partial r}\Big), \tag{S30}$$

where $D$ denotes the diffusion coefficient. Similarly, the dissipation rate (per area) of cross-membrane transport (e.g., Eq. S13) is given by

$$\Delta \dot{W}_{\text{transport}} = \kappa(c_+ - \gamma c_-) \times \Big(RT \ln \frac{c_+}{\gamma c_-}\Big), \tag{S31}$$

where $\kappa$ denotes the membrane transport rate, $c_\pm$ denotes the concentrations on either side of the membrane, and $\gamma$ denotes the reversibility of the transport, so that $\gamma = 1$ corresponds to a bidirectional channel and $\gamma < 1$ corresponds to a pump.

**Detailed derivation.** In order to concentrate $CO_2$ locally around Rubisco at nonequilibrium steady state, free energy must be dissipated through various nonequilibrium processes. Here, we compute the total energetic cost of operating the CCM, fixing $CO_2$, and making biomass. Our goal is to derive a general form for the total energetic cost that applies to an arbitrary localization of the CCM proteins. To simplify notation, we will express the energy cost in units of $RT$ and drop the term $RT$ from the equations below.

*Total energy consumption in the pyrenoid matrix:* The energetic cost of nonequilibrium processes that occur in the pyrenoid matrix is computed from Eqs. (S29–S31), which yields

$$\begin{aligned}\dot{W}_{\text{pyr}} = {} & I\big[0, R_{\text{pyr}}, D^C(\partial_r C_{\text{pyr}})(\partial_r \ln C_{\text{pyr}}), 1-f_{\text{v}}\big] + I\big[0, R_{\text{pyr}}, D^H(\partial_r H^-_{\text{pyr}})(\partial_r \ln H^-_{\text{pyr}}), 1-f_{\text{v}}\big] \\ & + I\big[0, R_{\text{pyr}}, D^H(\partial_r H^0_{\text{pyr}})(\partial_r \ln H^0_{\text{pyr}}), 1-f_{\text{v}}\big] + I\big[0, R_{\text{pyr}}, (j_{\text{sp}} + j_{\text{LCIB}}) \ln \frac{C_{\text{pyr}}}{K^{\text{eq}}_{\text{pyr}} H^-_{\text{pyr}}}, 1-f_{\text{v}}\big], \\ & + CO_2 \text{ fixation cost,}\end{aligned} \tag{S32}$$

where the energy dissipated by diffusion is described by the first three terms and the energy dissipated by converting $CO_2$ to $HCO_3^-$ is described by the fourth term (see Sec. I B for details). The energy cost of fixing $CO_2$ and converting it to biomass, termed "$CO_2$ fixation cost" in Eq. (S32), will be discussed below and further in Sec. II C. Recall that $K^{\text{eq}} \equiv 10^{\text{pK}_{\text{eff}} - \text{pH}}$ is the equilibrium $CO_2$-to-$HCO_3^-$ ratio in each compartment. We assume equal pH in the pyrenoid matrix and stroma, i.e., $K^{\text{eq}}_{\text{pyr}} = K^{\text{eq}}_{\text{str}}$. Integrating Eq. (S32) by parts and using the steady-state conditions Eqs. (S3, S8), we obtain

$$\begin{aligned}\dot{W}_{\text{pyr}} = {} & \big(-J_C \ln C_{\text{pyr}}\big)\big|_{R_{\text{pyr}}} + \big(J_{H^-} \ln H^-_{\text{pyr}}\big)\big|_{R_{\text{pyr}}} + \big(J_{H^0} \ln H^0_{\text{pyr}}\big)\big|_{R_{\text{pyr}}} + I\big[0, R_{\text{pyr}}, \kappa^C (C_{\text{thy}} - C_{\text{pyr}}) \ln C_{\text{pyr}}, f_{\text{s}} f_{\text{v}}\big] \\ & + I\big[0, R_{\text{pyr}}, (\kappa^{H^-} + \kappa^{H^-}_{\text{thy}})(H^-_{\text{thy}} - H^-_{\text{pyr}}) \ln H^-_{\text{pyr}}, f_{\text{s}} f_{\text{v}}\big] + I\big[0, R_{\text{pyr}}, \kappa^{H^0}(H^0_{\text{thy}} - H^0_{\text{pyr}}) \ln H^0_{\text{pyr}}, f_{\text{s}} f_{\text{v}}\big] \\ & - I\big[0, R_{\text{pyr}}, (j_{\text{sp}} + j_{\text{LCIB}}) \ln K^{\text{eq}}_{\text{pyr}}, 1-f_{\text{v}}\big] - I[0, R_{\text{pyr}}, j_{\text{Rbc}} \ln C_{\text{pyr}}, 1-f_{\text{v}}] + CO_2 \text{ fixation cost,}\end{aligned} \tag{S33}$$

where we have defined[1] outward $CO_2$ flux $J_C \equiv -4\pi r^2 (1-f_{\text{v}}) D^C \frac{\partial C}{\partial r}$, inward $HCO_3^-$ flux $J_{H^-} \equiv 4\pi r^2 (1-f_{\text{v}}) D^H \frac{\partial H^-}{\partial r}$ and inward $H_2CO_3$ flux $J_{H^0} \equiv 4\pi r^2 (1-f_{\text{v}}) D^H \frac{\partial H^0}{\partial r}$, and we have used the boundary conditions $\frac{\partial C_{\text{pyr}}}{\partial r}|_{r=0} = \frac{\partial H^0_{\text{pyr}}}{\partial r}|_{r=0} = \frac{\partial H^-_{\text{pyr}}}{\partial r}|_{r=0} = 0$.

*Total energy consumption in the stroma:* Similar to the calculation above, the energetic cost of nonequilibrium processes that occur in the stroma is given by

$$\begin{aligned}\dot{W}_{\text{str}} = {} & I\big[R_{\text{pyr}}, R_{\text{chlor}}, D^C(\partial_r C_{\text{str}})(\partial_r \ln C_{\text{str}}), 1-f_{\text{v}}\big] + I\big[R_{\text{pyr}}, R_{\text{chlor}}, D^H(\partial_r H^-_{\text{str}})(\partial_r \ln H^-_{\text{str}}), 1-f_{\text{v}}\big] \\ & + I\big[R_{\text{pyr}}, R_{\text{chlor}}, D^H(\partial_r H^0_{\text{str}})(\partial_r \ln H^0_{\text{str}}), 1-f_{\text{v}}\big] + I\big[R_{\text{pyr}}, R_{\text{chlor}}, (j_{\text{sp}} + j_{\text{LCIB}}) \ln \frac{C_{\text{str}}}{K^{\text{eq}}_{\text{str}} H^-_{\text{str}}}, 1-f_{\text{v}}\big] \\ & + CO_2 \text{ fixation cost.}\end{aligned} \tag{S34}$$

---

[1] Note that these definitions also apply to the overall inward/outward flux at the chloroplast envelope $r = R_{\text{chlor}}$, with $D^{C,H}$ replaced with $D^{C,H}_{\text{str}}$.

Integrating Eq. (S34) by parts and using the steady-state conditions Eqs. (S4, S9), we obtain

$$\begin{aligned}\dot{W}_{\rm str} = &\left(J_C \ln C_{\rm str}\right)\big|_{R_{\rm pyr}} - \left(J_C \ln C_{\rm str}\right)\big|_{R_{\rm chlor}} - \left(J_{H^-} \ln H^-_{\rm str}\right)\big|_{R_{\rm pyr}} + \left(J_{H^-} \ln H^-_{\rm str}\right)\big|_{R_{\rm chlor}} \\ &- \left(J_{H^0} \ln H^0_{\rm str}\right)\big|_{R_{\rm pyr}} + \left(J_{H^0} \ln H^0_{\rm str}\right)\big|_{R_{\rm chlor}} - I\left[R_{\rm pyr}, R_{\rm chlor}, (j_{\rm sp} + j_{\rm LCIB}) \ln K^{\rm eq}_{\rm str}, 1 - f_{\rm v}\right] \\ &+ I\left[R_{\rm pyr}, R_{\rm chlor}, \kappa^C (C_{\rm thy} - C_{\rm str}) \ln C_{\rm str}, f_{\rm s} f_{\rm v}\right] + I\left[R_{\rm pyr}, R_{\rm chlor}, \kappa^{H^0} (H^0_{\rm thy} - H^0_{\rm str}) \ln H^0_{\rm str}, f_{\rm s} f_{\rm v}\right] \\ &+ I\left[R_{\rm pyr}, R_{\rm chlor}, (\kappa^{H^-} + \kappa^{H^-}_{\rm thy})(H^-_{\rm thy} - H^-_{\rm str}) \ln H^-_{\rm str}, f_{\rm s} f_{\rm v}\right] \\ &- I[R_{\rm pyr}, R_{\rm chlor}, j_{\rm Rbc} \ln C_{\rm str}, 1 - f_{\rm v}] + {\rm CO_2 \ fixation\ cost}. \end{aligned} \tag{S35}$$

*Total energy consumption in the thylakoid lumen:* Similar to the calculations above, the energetic cost of nonequilibrium processes that occur in the thylakoid lumen including the tubules is given by

$$\begin{aligned}\dot{W}_{\rm thy} = &I\left[0, R_{\rm chlor}, D^C (\partial_r C_{\rm thy})(\partial_r \ln C_{\rm thy}), f_{\rm v}\right] + I\left[0, R_{\rm chlor}, D^H (\partial_r H^-_{\rm thy})(\partial_r \ln H^-_{\rm thy}), f_{\rm v}\right] \\ &+ I\left[0, R_{\rm chlor}, D^H (\partial_r H^0_{\rm str})(\partial_r \ln H^0_{\rm str}), f_{\rm v}\right] + I\left[0, R_{\rm chlor}, (j_{\rm sp} + j_{\rm CAH3}) \ln \frac{C_{\rm thy}}{K^{\rm eq}_{\rm thy} H^-_{\rm thy}}, f_{\rm v}\right]. \end{aligned} \tag{S36}$$

Integrating Eq. (S36) by parts and using the steady-state conditions Eqs. (S2, S4), we obtain

$$\begin{aligned}\dot{W}_{\rm thy} = &-I\left[0, R_{\rm pyr}, \kappa^C (C_{\rm thy} - C_{\rm pyr}) \ln C_{\rm thy}, f_{\rm s} f_{\rm v}\right] - I\left[R_{\rm pyr}, R_{\rm chlor}, \kappa^C (C_{\rm thy} - C_{\rm str}) \ln C_{\rm thy}, f_{\rm s} f_{\rm v}\right] \\ &- I\left[0, R_{\rm pyr}, (\kappa^{H^-} + \kappa^{H^-}_{\rm thy})(H^-_{\rm thy} - H^-_{\rm pyr}) \ln H^-_{\rm thy}, f_{\rm s} f_{\rm v}\right] \\ &- I\left[R_{\rm pyr}, R_{\rm chlor}, (\kappa^{H^-} + \kappa^{H^-}_{\rm thy})(H^-_{\rm thy} - H^-_{\rm str}) \ln H^-_{\rm thy}, f_{\rm s} f_{\rm v}\right] \\ &- I\left[0, R_{\rm pyr}, \kappa^{H^0} (H^0_{\rm thy} - H^0_{\rm pyr}) \ln H^0_{\rm thy}, f_{\rm s} f_{\rm v}\right] - I\left[R_{\rm pyr}, R_{\rm chlor}, \kappa^{H^0} (H^0_{\rm thy} - H^0_{\rm str}) \ln H^0_{\rm thy}, f_{\rm s} f_{\rm v}\right] \\ &- I\left[0, R_{\rm chlor}, (j_{\rm sp} + j_{\rm CAH3}) \ln K^{\rm eq}_{\rm thy}, f_{\rm v}\right], \end{aligned} \tag{S37}$$

where we have used the boundary conditions $\frac{\partial C_{\rm thy}}{\partial r}|_{r=0} = \frac{\partial C_{\rm thy}}{\partial r}|_{r=R_{\rm chlor}} = \frac{\partial H^0_{\rm thy}}{\partial r}|_{r=0} = \frac{\partial H^0_{\rm thy}}{\partial r}|_{r=R_{\rm chlor}} = \frac{\partial H^-_{\rm thy}}{\partial r}|_{r=0} = \frac{\partial H^-_{\rm thy}}{\partial r}|_{r=R_{\rm chlor}} = 0$.

*Total energy consumption of intercompartmental exchange:* We consider three intercompartmental transport processes — exchange of carbon molecules (1) between the thylakoid lumen and the pyrenoid matrix/stroma, (2) between the matrix and the stroma, and (3) between the stroma and the cytosol. Their energetic costs are calculated as follows:

$$\begin{aligned}\dot{W}^{(1)}_{\rm intercomp} = &I\left[0, R_{\rm pyr}, \kappa^C (C_{\rm thy} - C_{\rm pyr}) \ln \frac{C_{\rm thy}}{C_{\rm pyr}}, f_{\rm s} f_{\rm v}\right] + I\left[R_{\rm pyr}, R_{\rm chlor}, \kappa^C (C_{\rm thy} - C_{\rm str}) \ln \frac{C_{\rm thy}}{C_{\rm str}}, f_{\rm s} f_{\rm v}\right] \\ &+ I\left[0, R_{\rm pyr}, (\kappa^{H^-} + \kappa^{H^-}_{\rm thy})(H^-_{\rm thy} - H^-_{\rm pyr}) \ln \frac{H^-_{\rm thy}}{H^-_{\rm pyr}}, f_{\rm s} f_{\rm v}\right] \\ &+ I\left[R_{\rm pyr}, R_{\rm chlor}, (\kappa^{H^-} + \kappa^{H^-}_{\rm thy})(H^-_{\rm thy} - H^-_{\rm str}) \ln \frac{H^-_{\rm thy}}{H^-_{\rm str}}, f_{\rm s} f_{\rm v}\right] \\ &+ I\left[0, R_{\rm pyr}, \kappa^{H^0} (H^0_{\rm thy} - H^0_{\rm pyr}) \ln \frac{H^0_{\rm thy}}{H^0_{\rm pyr}}, f_{\rm s} f_{\rm v}\right] + I\left[R_{\rm pyr}, R_{\rm chlor}, \kappa^{H^0} (H^0_{\rm thy} - H^0_{\rm str}) \ln \frac{H^0_{\rm thy}}{H^0_{\rm str}}, f_{\rm s} f_{\rm v}\right], \end{aligned} \tag{S38}$$

$$\dot{W}^{(2)}_{\rm intercomp} = \left(J_C \ln \frac{C_{\rm pyr}}{C_{\rm str}}\right)\Big|_{R_{\rm pyr}} + \left(J_{H^-} \ln \frac{H^-_{\rm str}}{H^-_{\rm pyr}}\right)\Big|_{R_{\rm pyr}} + \left(J_{H^0} \ln \frac{H^0_{\rm str}}{H^0_{\rm pyr}}\right)\Big|_{R_{\rm pyr}}, \text{ and} \tag{S39}$$

$$\begin{aligned}\dot{W}^{(3)}_{\rm intercomp} = &\left(J_C\right)\big|_{R_{\rm chlor}} \ln \frac{(C_{\rm str})|_{R_{\rm chlor}}}{C_{\rm cyt}} + \left(J^{\rm diff}_{H^-}\right)\big|_{R_{\rm chlor}} \ln \frac{H^-_{\rm cyt}}{(H^-_{\rm str})|_{R_{\rm chlor}}} \\ &+ \left(J^{\rm trans}_{H^-}\right)\big|_{R_{\rm chlor}} \ln \frac{H^-_{\rm cyt}}{\gamma (H^-_{\rm str})|_{R_{\rm chlor}}} + \left(J_{H^0}\right)\big|_{R_{\rm chlor}} \ln \frac{H^0_{\rm cyt}}{(H^0_{\rm str})|_{R_{\rm chlor}}}, \end{aligned} \tag{S40}$$

where $J_{H^-}^{\rm diff}$ and $J_{H^-}^{\rm trans}$ denote, respectively, the inward $HCO_3^-$ flux via passive diffusion and via active $HCO_3^-$ transporter (see Sec. I E for details).

*Total energy consumption of reactions and transport of inorganic carbon molecules:* Summing up all the terms in Eqs. (S33, S35, S37 & S38 – S40) yields the energetic cost

$$\begin{aligned}\dot{W}_{\rm carbon} = &-\big(J_C\big)\big|_{R_{\rm chlor}} \ln C_{\rm cyt} + \big(J_{H^-}^{\rm diff}\big)\big|_{R_{\rm chlor}} \ln H_{\rm cyt}^- + \big(J_{H^-}^{\rm trans}\big)\big|_{R_{\rm chlor}} \ln(\gamma^{-1} H_{\rm cyt}^-) + \big(J_{H^0}\big)\big|_{R_{\rm chlor}} \ln H_{\rm cyt}^0 \\ &- J_{\rm str}^{C\to H^-} \ln K_{\rm str}^{\rm eq} + J_{\rm thy}^{H^-\to C} \ln K_{\rm thy}^{\rm eq} + \dot{W}_{\rm fixation} \\ &- I[0, R_{\rm pyr}, j_{\rm Rbc} \ln C_{\rm pyr}, 1-f_{\rm v}] - I[R_{\rm pyr}, R_{\rm chlor}, j_{\rm Rbc} \ln C_{\rm str}, 1-f_{\rm v}],\end{aligned} \tag{S41}$$

where $\dot{W}_{\rm fixation}$ denotes the total cost of $CO_2$ fixation and biomass production from the local $CO_2$ concentration around Rubisco. We have defined the total conversion fluxes from $CO_2$ to $HCO_3^-$ in the stroma and pyrenoid matrix as $J_{\rm str}^{C\to H^-} \equiv I[0, R_{\rm chlor}, j_{\rm sp} + j_{\rm LCIB}, 1-f_{\rm v}]$, and the total conversion fluxes from $HCO_3^-$ to $CO_2$ in the thylakoids as $J_{\rm thy}^{H^-\to C} \equiv -I[0, R_{\rm chlor}, j_{\rm sp} + j_{\rm CAH3}, f_{\rm v}]$, respectively. The total $CO_2$ fixation flux is given by $J_{\rm Rbc} = I[0, R_{\rm chlor}, j_{\rm Rbc}, 1-f_{\rm v}]$. Note that $(J_C)|_{R_{\rm chlor}}$ denotes the total $CO_2$ leakage flux out of the chloroplast and $(J_H)|_{R_{\rm chlor}} \equiv (J_{H^-}^{\rm diff})|_{R_{\rm chlor}} + (J_{H^-}^{\rm trans})|_{R_{\rm chlor}} + (J_{H^0})|_{R_{\rm chlor}}$ denotes the total inward flux of $HCO_3^-$ and $H_2CO_3$. Thus, we obtain two global flux-balance conditions: (1) $(J_H)|_{R_{\rm chlor}} = (J_C)|_{R_{\rm chlor}} + J_{\rm Rbc}$, since the carbon molecules entering the chloroplast will either be fixed by Rubisco or diffuse out again; and (2) $J_{\rm thy}^{H^-\to C} = J_{\rm str}^{C\to H^-} + (J_H)|_{R_{\rm chlor}}$, since the $HCO_3^-$ molecules converted to $CO_2$ in the thylakoids are either imported from outside the chloroplast or are recycling products in the stroma and pyrenoid matrix. Using these two relations to replace $(J_{H^-}^{\rm trans})|_{R_{\rm chlor}}$ and $J_{\rm thy}^{H^-\to C}$ in Eq. (S41), we obtain

$$\begin{aligned}\dot{W}_{\rm carbon} = &J_{\rm str}^{C\to H^-} \ln \frac{K_{\rm thy}^{\rm eq}}{K_{\rm str}^{\rm eq}} + \big(J_C\big)\big|_{R_{\rm chlor}} \ln \frac{\gamma^{-1} K_{\rm thy}^{\rm eq}}{K_{\rm str}^{\rm eq}} - \big(J_{H^-}^{\rm diff}\big)\big|_{R_{\rm chlor}} \ln \gamma^{-1} - \big(J_{H^0}\big)\big|_{R_{\rm chlor}} \ln \frac{\gamma^{-1} H_{\rm cyt}^0}{H_{\rm cyt}^-} + \dot{W}_{\rm fixation} \\ &+ J_{\rm Rbc} \ln \frac{\gamma^{-1} K_{\rm thy}^{\rm eq}}{K_{\rm cyt}^{\rm eq}} + I\Big[0, R_{\rm pyr}, j_{\rm Rbc} \ln \frac{C_{\rm cyt}}{C_{\rm pyr}}, 1-f_{\rm v}\Big] + I\Big[R_{\rm pyr}, R_{\rm chlor}, j_{\rm Rbc} \ln \frac{C_{\rm cyt}}{C_{\rm str}}, 1-f_{\rm v}\Big].\end{aligned} \tag{S42}$$

Note that in deriving Eq. (S42) we have assumed fast equilibrium between $CO_2$ and $HCO_3^-$ in the cytosol, i.e., $C_{\rm cyt}/H_{\rm cyt}^- = K_{\rm cyt}^{\rm eq}$.

*Total energy consumption of proton import:* So far, we have only considered the energy cost associated with nonequilibrium reaction and transport of inorganic carbon molecules. Note that we have assumed constant pHs in different compartments and we have not yet explicitly considered the reaction and transport of protons. Nevertheless, protons are consumed by the reaction $HCO_3^- + H^+ \to CO_2 + H_2O$, and thus equal amounts of protons and $HCO_3^-$ need to be taken into the chloroplast to maintain charge neutrality. Proton import from cytosolic pH to stromal pH yields a dissipation of

$$\dot{W}_{\rm proton} = (J_{H^-})|_{R_{\rm chlor}} \ln 10^{-{\rm pH}_{\rm cyt} + {\rm pH}_{\rm str}} = (J_{H^-})|_{R_{\rm chlor}} \ln \frac{K_{\rm cyt}^{\rm eq}}{K_{\rm str}^{\rm eq}}. \tag{S43}$$

*Total energy consumption of $CO_2$ fixation:* Finally, we consider the energy cost in reactions of $CO_2$ fixation and biomass production. We choose air-level $CO_2$ $C_{\rm air} = 10$ µM as our reference concentration and denote by $\epsilon_{\rm b}$ the free-energy difference between $CO_2$ at $C_{\rm air}$ and in carbon biomass. Thus, the total energy cost of $CO_2$ fixation $\dot{W}_{\rm fixation}$ in Eq. (S42) is given by

$$\dot{W}_{\rm fixation} = J_{\rm Rbc} \epsilon_{\rm b} + I\Big[0, R_{\rm pyr}, j_{\rm Rbc} \ln \frac{C_{\rm pyr}}{C_{\rm air}}, 1-f_{\rm v}\Big] + I\Big[R_{\rm pyr}, R_{\rm chlor}, j_{\rm Rbc} \ln \frac{C_{\rm str}}{C_{\rm air}}, 1-f_{\rm v}\Big], \tag{S44}$$

where the last two terms account for the free-energy difference of $CO_2$ between the local concentration at Rubisco and the reference concentration.

**Summary and discussion.** Combining Eq. (S42), (S43) and (S44), we obtain the total energetic cost (in units of $RT$) to be $\dot{W}_{\rm tot} = \dot{W}_{\rm CCM} + \dot{W}_{\rm biomass}$, where $\dot{W}_{\rm biomass} = J_{\rm Rbc}\epsilon_{\rm b}$ denotes the cost of fixing $CO_2$ and making biomass and $\dot{W}_{\rm CCM}$ denotes the cost of operating the CCM, which is given by

$$\dot{W}_{\rm CCM} = J_{\rm str}^{C\to H^-}\ln\frac{K_{\rm thy}^{\rm eq}}{K_{\rm str}^{\rm eq}} + \big(J_{\rm C}\big)\big|_{R_{\rm chlor}}\ln\frac{\gamma^{-1}K_{\rm thy}^{\rm eq}}{K_{\rm str}^{\rm eq}} + J_{\rm Rbc}\Big(\ln\frac{\gamma^{-1}K_{\rm thy}^{\rm eq}}{K_{\rm str}^{\rm eq}} + \ln\frac{C_{\rm cyt}}{C_{\rm air}}\Big)$$
$$- \big(J_{H^-}^{\rm diff}\big)\big|_{R_{\rm chlor}}\ln\gamma^{-1} - \big(J_{H^0}\big)\big|_{R_{\rm chlor}}\ln\Big(\frac{\gamma^{-1}K_{\rm cyt}^{\rm eq}}{K_{\rm str}^{\rm eq}}10^{\rm pKa_1 - pH_{cyt}}\Big), \qquad \text{(S45)}$$

where $\rm pKa_1 = 3.4$ denotes the first pKa of $H_2CO_3$ [4]. One can readily normalize Eq. (S45) by $J_{\rm Rbc}$ and use ATP hydrolysis energy $|\Delta G_{\rm ATP}| = 20.8RT$ (see [14] Box 13-1) to compute the equivalent ATP cost per $CO_2$ fixed to operate the CCM, i.e., $\epsilon_C \equiv \dot{W}_{\rm CCM}/J_{\rm Rbc}/20.8$.

Note that since the chloroplast membrane is quite impermeable to charged molecules like $HCO_3^-$, the influx of $HCO_3^-$ via passive diffusion across the chloroplast envelope $\big(J_{H^-}^{\rm diff}\big)\big|_{R_{\rm chlor}}$ is negligible. Note also that all of the compartments in our model have pHs much larger than $\rm pKa_1$. Thus, the concentration of $H_2CO_3$ and its transport flux are also negligible compared to those of $HCO_3^-$. Neglecting the last two terms in Eq. (S45), we obtain a simplified (approximate) expression for the energetic cost of the CCM at air-level $CO_2$ (10 μM cytosolic) in units of $RT$:

$$\dot{W}_{\rm CCM} \approx J_{\rm str}^{C\to H^-}\ln\frac{K_{\rm str}^{\rm eq}}{K_{\rm thy}^{\rm eq}} + \big(J_{\rm C}\big)\big|_{R_{\rm chlor}}\ln\frac{\gamma^{-1}K_{\rm str}^{\rm eq}}{K_{\rm thy}^{\rm eq}} + J_{\rm Rbc}\ln\frac{\gamma^{-1}K_{\rm str}^{\rm eq}}{K_{\rm thy}^{\rm eq}}. \qquad \text{(S46)}$$

The three terms in Eq. (S46) have clear physical meanings (Fig. S13). The first term describes the energetic cost of LCIB participating in a futile cycle: $HCO_3^-$ that is transported into the thylakoids is converted to $CO_2$, which then diffuses into the pyrenoid matrix and may diffuse back out into the stroma, where it will be recycled back to $HCO_3^-$ by LCIB. For each cycle, the free energy dissipated $\sim \ln\frac{K_{\rm thy}^{\rm eq}}{K_{\rm str}^{\rm eq}} = \ln 10^{\rm pH_{str} - pH_{thy}}$ is equal to the energy needed to pump a proton from the stromal pH to the thylakoid pH. The second term describes the free energy wasted by $CO_2$ leakage, where the import of one molecule of $HCO_3^-$ costs $\sim \ln\gamma^{-1}$ and pumping a proton into the thylakoid lumen to convert $HCO_3^-$ to $CO_2$ costs $\sim \ln\frac{K_{\rm thy}^{\rm eq}}{K_{\rm str}^{\rm eq}}$. The third term describes the energetic cost of these same processes, but for those $CO_2$ molecules that are eventually fixed by Rubisco.

##### C. A combined measure of CCM performance

As seen in Figs. 3 and 4, different CCM performances can be achieved by varying enzyme activity and localization, as well as $HCO_3^-$ transport rates. In this section, we seek to define a combined measure of CCM performance that takes into account both the efficacy and the cost of concentrating $CO_2$. We then employ this combined measure to compare different CCM strategies.

**Energy and flux balance.** For simplicity, we assume that the Chlamydomonas cell is partitioned into two parts, one with biomass $B_{\rm c}$ performing the CCM and the other with biomass $(B_{\rm tot} - B_{\rm c})$ collecting light energy (Fig. S24). The number of photons absorbed by the cell per unit time is assumed to be $\Gamma_{\rm ph} = \alpha(B_{\rm tot} - B_{\rm c})$ where $\alpha$ is a proportionality constant and depends on the incident light intensity. Part of the absorbed photons $\Gamma_{\rm m} = \nu B_{\rm tot}$ is used for maintenance [45], where $\nu$ denotes the number of maintenance photons per biomass. Thus, the total energy input that can be used for running the CCM and synthesizing biomass is given by $P_{\rm tot} = \epsilon_{\rm ph}(\Gamma_{\rm ph} - \Gamma_{\rm m})$, where $\epsilon_{\rm ph}$ denotes the yield of chemical energy per absorbed photon.

For a particular CCM strategy $\mathcal{S}$, the total $CO_2$ fixation flux by Rubisco is given by $\Gamma_C(\mathcal{S}) = \mu B_{\rm c}\Phi_C(\mathcal{S})$, where $\mu$ deonotes the maximum rate of $CO_2$ fixation per CCM biomass and $\Phi_C$ is the normalized $CO_2$ fixation flux. The total cost of concentrating $CO_2$ is given by $\epsilon_C(\mathcal{S})\Gamma_C(\mathcal{S})$, where $\epsilon_C$ denotes the energy cost per $CO_2$ fixed. However, some of the $CO_2$ fixed is then released through Rubisco's competitive oxygenation reaction. The total $CO_2$ flux lost through this reaction is given by $\Gamma_O(\mathcal{S}) = \mu B_{\rm c}\Phi_O(\mathcal{S})$ in which $\Phi_O$ is the normalized loss of $CO_2$ by oxygenation (Eq. S28), with a cost of $\epsilon_O$ per $CO_2$ lost. After fixing a net $CO_2$ flux of $\Gamma_C - \Gamma_O$ to organic carbon, the cell next converts this to biomass at a constant cost $\epsilon_{\rm b}$ per carbon. The total biomass carbon production flux is denoted by $\Gamma_{\rm b}$. Thus, the balance of energy and carbon flux requires that

$$\Gamma_C(\mathcal{S})\epsilon_C(\mathcal{S}) + \Gamma_{\rm b}\epsilon_{\rm b} + \Gamma_O(\mathcal{S})\epsilon_O \leqslant P_{\rm tot},\ \text{and} \qquad \text{(S47a)}$$

$$\Gamma_{\rm b} \leqslant \Gamma_C - \Gamma_O. \qquad \text{(S47b)}$$

The specific growth rate of the cell $g$ can be computed as $g = \frac{\Gamma_\mathrm{b}}{\chi_C B_\mathrm{tot}}$, where $\chi_C$ is the fraction of carbon in biomass. Below, we discuss the optimal CCM strategy which maximizes $g$.

**Optimal CCM strategy.**

*Constrained optimization:* Consider a cell with a fixed CCM biomass fraction $f_\mathrm{c}^0 = B_\mathrm{c}^0/B_\mathrm{tot}$. The specific growth rate depends on the choice of CCM strategy as follows (Fig. S24A): when $\Gamma_C$ (and/or $\epsilon_C$) is small, growth is limited by $CO_2$ fixation, i.e., $\Gamma_\mathrm{b} = \Gamma_C(\mathcal{S}) - \Gamma_O(\mathcal{S}) = \mu\Psi_C(\mathcal{S})B_\mathrm{tot}$ (Eq. S28), and the excess light energy is dissipated; when $\Gamma_C$ (and/or $\epsilon_C$) is large, growth rate becomes energy limited, which yields $\Gamma_\mathrm{b} = \epsilon_\mathrm{b}^{-1}\big[P_\mathrm{tot} - \Gamma_C(\mathcal{S})\epsilon_C(\mathcal{S}) - \Gamma_O(\mathcal{S})\epsilon_O\big]$. Taken together, we obtain

$$g^0(\mathcal{S})\chi_C = \begin{cases} \mu\Psi_C(\mathcal{S})f_\mathrm{c}^0 & \text{when } \mu f_\mathrm{c}^0\big[\Phi_C(\epsilon_\mathrm{b}+\epsilon_C)+\Phi_O(\epsilon_O-\epsilon_\mathrm{b})\big] \\ & \qquad \le \epsilon_\mathrm{ph}\big[\alpha(1-f_\mathrm{c}^0)-\nu\big], \\ \frac{\epsilon_\mathrm{ph}}{\epsilon_\mathrm{b}}\big[\alpha(1-f_\mathrm{c}^0)-\nu\big] - \frac{\epsilon_C}{\epsilon_\mathrm{b}}\mu\Phi_C(\mathcal{S})f_\mathrm{c}^0 - \frac{\epsilon_O}{\epsilon_\mathrm{b}}\mu\Phi_O(\mathcal{S})f_\mathrm{c}^0 & \text{otherwise.} \end{cases} \tag{S48}$$

The optimal CCM strategy that maximizes the growth rate given a particular biomass partition is given by $\mathcal{S}^{\dagger,0} = \arg\max_{\mathcal{S}}\big(g^0(\mathcal{S})\big)$ (Figs. S24B, F, and I).

*Global optimization:* Next we consider a cell that can adjust its CCM biomass fraction $f_\mathrm{c} = B_\mathrm{c}/B_\mathrm{tot}$ and the CCM strategy to maximize growth. In this case, for a particular $\mathcal{S}$, the growth rate is maximized when both $\Gamma_\mathrm{b} = \Gamma_C - \Gamma_O$ and $P_\mathrm{tot} = \Gamma_\mathrm{b}\epsilon_\mathrm{b} + \Gamma_C\epsilon_C + \Gamma_O\epsilon_O$ are satisfied, which yields an optimal CCM biomass fraction

$$f_\mathrm{c}^*(\mathcal{S}) = \frac{\alpha-\nu}{\alpha+\mu\Big[\Phi_C(\mathcal{S})\frac{\epsilon_\mathrm{b}+\epsilon_C(\mathcal{S})}{\epsilon_\mathrm{ph}} + \Phi_O(\mathcal{S})\frac{\epsilon_O-\epsilon_\mathrm{b}}{\epsilon_\mathrm{ph}}\Big]}. \tag{S49}$$

Thus, the biomass growth rate is given by

$$g(\mathcal{S}) = \mu\Psi_C(\mathcal{S})f_\mathrm{c}^*(\mathcal{S})\chi_C^{-1} = \chi_C^{-1}(\alpha-\nu)\Big(1-\frac{\Phi_O(\mathcal{S})}{\Phi_C(\mathcal{S})}\Big)\Big[\frac{\alpha}{\mu\Phi_C(\mathcal{S})} + \frac{\epsilon_\mathrm{b}+\epsilon_C(\mathcal{S})}{\epsilon_\mathrm{ph}} + \frac{\Phi_O(\mathcal{S})}{\Phi_C(\mathcal{S})}\Big(\frac{\epsilon_O-\epsilon_\mathrm{b}}{\epsilon_\mathrm{ph}}\Big)\Big]^{-1}. \tag{S50}$$

Optimizing $g$ with respect to $\mathcal{S}$ yields the globally optimal CCM strategy $\mathcal{S}^\dagger$ and growth rate $g^\dagger$, i.e., $\mathcal{S}^\dagger = \arg\max_{\mathcal{S}}\big(g(\mathcal{S})\big)$ and $g^\dagger = g(\mathcal{S}^\dagger)$ (Figs. S24C, G, and J).

**Values of parameters.** The reaction cost of biomass synthesis from $CO_2$ is $\epsilon_\mathrm{b} \approx 1000$ kJ/C-mol biomass = 20 ATP/C biomass [46].[2] The energy cost of oxygenation per $CO_2$ lost can be computed to be $\epsilon_O \approx 26.4$ ATP [4, 47]. Note that $\epsilon_O$ is two times the energy cost of a oxygenation reaction due to the stoichiometry. The fraction of carbon in biomass is measured to be $\chi_C = 0.48$ g C/g biomass = 0.04 mol-C/g biomass [48].

The yield of chemical energy per photon $\epsilon_\mathrm{ph}$ and the maintenance photon-flux coefficient $\nu$ are estimated from the chemostat experiments in [45]. Briefly, Chlamydomonas cells were grown in a chemostat at various dilution rates $d$, i.e., specific growth rates, and the corresponding numbers of absorbed photons per biomass per unit time $R_\mathrm{ph}$ were measured. $R_\mathrm{ph}$ was found to increase linearly with $d$, where the slope $N_\mathrm{ph}$ is the number of photons absorbed per synthesized biomass and the non-zero offset is the number of photons needed for maintenance, yielding $\nu = 6$ mmol/g biomass/h = 1.67 µmol/g biomass/s. $N_\mathrm{ph}$ was measured to be 0.8 mol photons/g biomass. Note that the energy requirement per biomass can be calculated by $N_\mathrm{ph}\epsilon_\mathrm{ph} = \epsilon_\mathrm{b}\chi_C$. Plugging in the values of $\chi_C, \epsilon_\mathrm{b}$, and $N_\mathrm{ph}$, we obtain $\epsilon_\mathrm{ph} = 1$ ATP/photon. Note that this number is consistent with an independent calculation based on the light reactions of photosynthesis. For linear electron flux (LEF), 8 photons transport 4 electrons, leading to formation of 2 NADPH (8.7 ATP equivalent energy) and the pumping of 12 protons (2.6 ATP equivalent energy) from the stroma to the thylakoid lumen [49, 50]. This gives a chemical energy yield of 1.4 ATP per photon. For cyclic electron flux (CEF), 1 photon drives the cyclic flux of 1 electron, leading to 2 protons pumped from the stroma to the thylakoid lumen, giving a chemical energy yield of 0.43 ATP per photon. In reality, it is likely that the cell uses a mixture of LEF and CEF to maintain a rapid stoichiometric balance of ATP and NADPH [51, 52], and thus $\epsilon_\mathrm{ph}$ is expected to be within the range of 0.43 − 1.4 ATP per photon.[3]

---

[2] This value is for an electron donor that does not require reverse electron flow. The reaction cost is generally between 20 ATP and 60 ATP per biomass carbon.

[3] Here, we have chosen $\epsilon_\mathrm{b} = 20$ ATP per carbon to obtain a simple estimate of $\epsilon_\mathrm{ph} = 1$ ATP / photon.

As discussed above, $\alpha$ depends on the light intensity. Nevertheless, we can estimate the maximum $\alpha$ from light-saturating measurements of photosynthesis (e.g., [53]). In these experiments, cells were grown in medium supplemented with excess inorganic carbon (e.g., > 10 mM $HCO_3^-$). The measured photosynthesis activity first increases with increasing light intensity and eventually saturates for light intensity larger than $I_S \approx 200$ µmol photons $\cdot$ m$^{-2} \cdot$ s$^{-1}$. Thus, the saturating light absorption by a cell is given by $\Gamma_{\mathrm{ph,S}} = I_S \times \Delta S \times (a^*_{\mathrm{Chl}} C_{\mathrm{Chl}} \ell)$, where $\Delta S$ is the effective illuminated area of a cell, $\ell$ is the (average) path length through the cell, $a^*_{\mathrm{Chl}}$ is the absorption coefficient (cross section) of chlorophyll and $C_{\mathrm{Chl}}$ is the concentration of chlorophyll. $\Gamma_{\mathrm{ph,S}}$ can be rewritten as $\Gamma_{\mathrm{ph,S}} = I_S \times a^*_{\mathrm{Chl}} \times N_{\mathrm{Chl}}$, where $N_{\mathrm{Chl}}$ is the total number of chlorophyll per cell.[4] Plugging in measurements from [54], $a^*_{\mathrm{Chl}} = 10$ m$^2$/(g Chl) and $N_{\mathrm{Chl}} = 1.5$ pg/cell yields $\Gamma_{\mathrm{ph,S}} = 3 \times 10^{-9}$ µmol photons/cell/s. The maximum $\alpha$ can be readily calculated below after we estimate the biomass.

It has been measured that a typical Chlamydomonas cell has a total biomass of $B_{\mathrm{tot}} = 43$ pg [54] and contains $B^{(\mathrm{prot})}_{\mathrm{tot}} \approx 20$ pg protein in total [55]. The masses of the major $CO_2$ fixation proteins were measured to be the following [55]: Rubisco large subunit rbcL $B^{(\mathrm{prot})}_{\mathrm{rbcL}} = 1.32$ pg, Rubisco small subunit RBCS $B^{(\mathrm{prot})}_{\mathrm{RBCS}} = 0.33$ pg, pyrenoid linker protein EPYC1 $B^{(\mathrm{prot})}_{\mathrm{EPYC1}} = 0.05$ pg, and Rubisco activase RCA $B^{(\mathrm{prot})}_{\mathrm{RCA}} = 0.02$ pg.[5] Thus, we estimate the total biomass responsible for $CO_2$ fixation to be $B^{(\mathrm{prot})}_{\mathrm{c}} \approx B^{(\mathrm{prot})}_{\mathrm{rbcL}} + B^{(\mathrm{prot})}_{\mathrm{RBCS}} + B^{(\mathrm{prot})}_{\mathrm{EPYC1}} + B^{(\mathrm{prot})}_{\mathrm{RCA}} = 1.72$ pg, which yields a CCM biomass fraction $f^0_{\mathrm{c}} \approx 0.086$. Introducing the values of $f^0_{\mathrm{c}}$ and $B_{\mathrm{tot}}$ into the relation $\alpha_S = \frac{\Gamma_{\mathrm{ph,S}}}{B_{\mathrm{tot}}(1-f^0_c)}$ yields the maximum rate of photon absorption $\alpha_S \approx 76$ µmol/g biomass/s. Finally, the maximum $CO_2$ fixation flux by Rubisco can be calculated from the turnover rate and the total number of active Rubisco catalytic sites [16, 26], which yields $\Gamma_{C,\mathrm{max}} = 0.75 \times 10^{-16}$ mol C/cell/s. Thus, $\mu = \frac{\Gamma_{C,\mathrm{max}}}{B_{\mathrm{tot}} f^0_{\mathrm{c}}} = 20.3$ µmol/g biomass/s.

All the estimated parameter values from experiments are summarized in Table S2. Guided by our theoretical framework, we wondered whether the cell optimizes its biomass fraction according to Eq. (S49). In particular, we consider the case where both $CO_2$ and light are in excess. In this case, there is no cost to concentrate $CO_2$, i.e., $\epsilon_C = 0$, there is no oxygenation $\Phi_O = 0$, and the normalized $CO_2$ fixation flux is $\Phi_C = 1$. Thus, using the parameter values discussed above, Eq. (S49) yields $f^*_{\mathrm{c}} \approx 0.15$, larger than the estimated $f^0_{\mathrm{c}}$. As shown in Fig. S24B, this suggests that biomass growth of Chlamydomonas under saturating light conditions is predominantly limited by $CO_2$ fixation, and the growth rate is simply proportional to NNCFF (see Eq. (S28)).

---

[4] It can be seen here that the saturating energy input from light is proportional to the total biomass of chlorophyll, which justifies our expression $P_{\mathrm{tot}} \propto (B_{\mathrm{tot}} - B_{\mathrm{c}})$.

[5] Note that cells are grown in TAP medium under non-saturating light in the experiments in [55]. Nevertheless, the relative abundance of the four proteins above are in line with [16], and the absolute abundance of rbcL and RBSC are in line with [26]. Specifically, the total number of Rubisco can be calculated from its measured density in the pyrenoid matrix 0.63 mM in [26] and the volume of the pyrenoid 3.5 µm$^3$, which yields $N_{\mathrm{rbc}} \approx 17.64$ amol assuming each Rubisco holoenzyme contains 8 large subunits and 8 small subunits. This agrees well with the reported value $N_{\mathrm{rbcL}} = 25.2$ amol and $N_{\mathrm{RBCS}} = 16.1$ amol in [55].

#### III. WELL-MIXED COMPARTMENT MODEL WITH AN IMPERMEABLE STARCH SHEATH

To gain more analytical insights into the two CCM strategies illustrated in Fig. 3, we consider a simplification of the full reaction-diffusion model with an impermeable starch sheath. We assume that

1. Diffusion of $CO_2$ and $HCO_3^-$ is fast in the pyrenoid matrix and stroma, and hence each of their concentrations can be described by a single number, $C_{\rm pyr}$, $C_{\rm str}$, $H^-_{\rm pyr}$, and $H^-_{\rm str}$, in the two compartments (Fig. S15A).
2. $HCO_3^-$ transport across the thylakoid membranes is fast. Thus, the thylakoid tubule concentrations of $HCO_3^-$ inside and outside the pyrenoid can be approximated by $H^-_{\rm pyr}$ and $H^-_{\rm str}$, respectively.
3. CAH3 is fast and hence $CO_2$ and $HCO_3^-$ are in fast equilibrium in the thylakoid tubules inside the pyrenoid where CAH3 is localized. Thus, the concentration of $CO_2$ can be described by a single number, $C_{\rm thy} = K^{\rm eq}_{\rm thy} H^-_{\rm pyr}$, in this intra-pyrenoid portion of the thylakoid tubules. Here, $K^{\rm eq}_{\rm thy} = 10^{0.1}$ is defined in Sec. I B.
4. The concentration of $CO_2$ in the thylakoid tubules near the chloroplast envelope approaches $C_{\rm str}$ (see for example Fig. S6D).

We term this model the well-mixed compartment model, which has four unknowns $C_{\rm pyr}, C_{\rm thy}, C_{\rm str}$, and $H^-_{\rm str}$. In Sec. III A we derive the algebraic governing equations. In Sec. III B we consider two extremes of the various CCM strategies, one employing passive uptake of $CO_2$ and one employing active pumping of $HCO_3^-$. In Sec. III C we discuss the full CCM strategy space under various external $CO_2$ conditions. Finally, in Sec. III D we discuss the biophysical limits posed by the passive $CO_2$ uptake CCM strategy.

##### A. Governing equations

The flux-balance conditions of inorganic carbon for the intra-pyrenoid portion of the chloroplast yield

$$\underbrace{2D^H a_{\rm tub} N_{\rm tub}(H^-_{\rm str} - C_{\rm thy}/K^{\rm eq}_{\rm thy})}_{\text{HCO}_3^- \text{ influx via thylakoid tubules}} = \underbrace{D^C(\pi a^2_{\rm tub}/L_{\rm thy}) N_{\rm tub}(C_{\rm thy} - C_{\rm str})}_{\text{CO}_2 \text{ efflux via thylakoid tubules}} + \underbrace{\Delta S_{\rm thy}\kappa^C(C_{\rm thy} - C_{\rm pyr})}_{\substack{\text{CO}_2 \text{ flux from the thylakoid} \\ \text{lumen to the pyrenoid matrix}}}, \text{ and} \tag{S51}$$

$$\Delta S_{\rm thy}\kappa^C(C_{\rm thy} - C_{\rm pyr}) = \underbrace{I[0, R_{\rm pyr}, 1, 1-f_{\rm v}] V^C_{\rm max,Rbc} \frac{C_{\rm pyr}}{C_{\rm pyr} + K^{\rm eff}_{\rm m}}}_{\text{CO}_2 \text{ fixation flux by Rubisco}}. \tag{S52}$$

Here, $L_{\rm thy}$ is the typical diffusion length along the thylakoid tubules, which is obtained by fitting the well-mixed compartment model to the full simulation (see below and Fig. S15). The diffusion flux of $HCO_3^-$ through a circular hole of radius $a_{\rm tub}$ in an otherwise impermeable membrane is approximated by $2D^H a_{\rm tub}\Delta H$ [56]. $\Delta S_{\rm thy} = I[0, R_{\rm pyr}, f_{\rm s}, f_{\rm v}]$ denotes the total surface area of thylakoid membranes in the pyrenoid.

The flux balance of $CO_2$ in the chloroplast stroma is given by

$$D^C(\pi a^2_{\rm tub}/L_{\rm thy}) N_{\rm tub}(C_{\rm thy} - C_{\rm str}) = \underbrace{4\pi R^2_{\rm chlor}\kappa^C(C_{\rm str} - C_{\rm cyt})}_{\text{CO}_2 \text{ flux out of the chloroplast}} + \underbrace{I[R_{\rm pyr}, R_{\rm chlor}, 1, 1-f_{\rm v}] \frac{V^C_{\rm max,LCIB}}{K^C_{\rm m}}(C_{\rm str} - K^{\rm eq}_{\rm str} H^-_{\rm str})}_{\text{conversion from CO}_2 \text{ to HCO}_3^-}. \tag{S53}$$

Here, we assume that LCIB is unsaturated and therefore its reaction is described by first-order kinetics.

Similarly, the flux balance of $HCO_3^-$ in the stroma yields

$$2D^H a_{\rm tub} N_{\rm tub}(H^-_{\rm str} - C_{\rm thy}/K^{\rm eq}_{\rm thy}) = \underbrace{4\pi R^2_{\rm chlor}\kappa^{H^-}_{\rm chlor}(H^-_{\rm cyt} - \gamma H^-_{\rm str})}_{\text{HCO}_3^- \text{ uptake from the cytosol}} + I[R_{\rm pyr}, R_{\rm chlor}, 1, 1-f_{\rm v}] \frac{V^C_{\rm max,LCIB}}{K^C_{\rm m}}(C_{\rm str} - K^{\rm eq}_{\rm str} H^-_{\rm str}), \tag{S54}$$

in which we assume that $CO_2$ and $HCO_3^-$ are in fast equilibrium in the cytosol $H^-_{\rm cyt} = 10\ C_{\rm cyt}$ (see also Sec. I E).

Equations (S51–S54) can be written as

$$\alpha^{H^-}_{\rm thy}(H^-_{\rm str} - C_{\rm thy}/K^{\rm eq}_{\rm thy}) = \alpha^C_{\rm thy}(C_{\rm thy} - C_{\rm str}) + \sigma^C_{\rm thy}(C_{\rm thy} - C_{\rm pyr}), \tag{S55a}$$

$$\sigma^C_{\rm thy}(C_{\rm thy} - C_{\rm pyr}) = \frac{C_{\rm pyr}}{C_{\rm pyr} + K^{\rm eff}_{\rm m}}, \tag{S55b}$$

$$\alpha_{\text{thy}}^{C}(C_{\text{thy}} - C_{\text{str}}) = \sigma_{\text{chl}}^{C}(C_{\text{str}} - C_{\text{cyt}}) + \beta_{\text{LCIB}}(C_{\text{str}} - K_{\text{str}}^{\text{eq}} H_{\text{str}}^{-}), \tag{S55c}$$

$$\alpha_{\text{thy}}^{H^-}(H_{\text{str}}^{-} - C_{\text{thy}}/K_{\text{thy}}^{\text{eq}}) = \sigma_{\text{chl}}^{H^-}(10\ C_{\text{cyt}} - \gamma H_{\text{str}}^{-}) + \beta_{\text{LCIB}}(C_{\text{str}} - K_{\text{str}}^{\text{eq}} H_{\text{str}}^{-}), \tag{S55d}$$

where we have grouped the parameters into the following inverse concentrations: $\alpha_{\text{thy}}^{H^-} = \frac{2D^H a_{\text{tub}} N_{\text{tub}}}{I[0,R_{\text{pyr}},1,1-f_{\text{v}}]V_{\text{max,Rbc}}^{C}} = 93.2\ \text{mM}^{-1}$, $\alpha_{\text{thy}}^{C} = \frac{D^C(\pi a_{\text{tub}}^2/L_{\text{thy}})N_{\text{tub}}}{I[0,R_{\text{pyr}},1,1-f_{\text{v}}]V_{\text{max,Rbc}}^{C}} = 13.0\ \text{mM}^{-1}$, $\sigma_{\text{thy}}^{C} = \frac{\Delta S_{\text{thy}}\kappa^C}{I[0,R_{\text{pyr}},1,1-f_{\text{v}}]V_{\text{max,Rbc}}^{C}} = 51.6\ \text{mM}^{-1}$, $\sigma_{\text{chl}}^{C} = \frac{4\pi R_{\text{chlor}}^2\kappa^C}{I[0,R_{\text{pyr}},1,1-f_{\text{v}}]V_{\text{max,Rbc}}^{C}} = 753\ \text{mM}^{-1}$, $\beta_{\text{LCIB}} = \frac{I[R_{\text{pyr}},R_{\text{chlor}},1,1-f_{\text{v}}]V_{\text{max,LCIB}}^{C}/K_{\text{m}}^{C}}{I[0,R_{\text{pyr}},1,1-f_{\text{v}}]V_{\text{max,Rbc}}^{C}}$, $\sigma_{\text{chl}}^{H^-} = \frac{4\pi R_{\text{chlor}}^2\kappa_{\text{chlor}}^{H^-}}{I[0,R_{\text{pyr}},1,1-f_{\text{v}}]V_{\text{max,Rbc}}^{C}}$. The value of $\alpha_{\text{thy}}^{C}$ is obtained by fitting the well-mixed compartment model to the full simulation (Fig. S15B). All the notations and parameter values are summarized in Table S3.

#### B. Two extreme CCM strategies

Different CCM strategies are parameterized by $\beta_{\text{LCIB}}$, $\sigma_{\text{chl}}^{H^-}$ and $\gamma$ in the well-mixed compartment model. Here, we start by considering the extremes of the two CCM strategies compared in Fig. 3.

**Extreme 1: fast LCIB, no $HCO_3^-$ transport across the chloroplast envelope.** In this limit, $\beta_{\text{LCIB}} \to \infty$ and $\sigma_{\text{chl}}^{H^-} = 0$, so $CO_2$ and $HCO_3^-$ are in fast equilibrium in the stroma, i.e., $C_{\text{str}} = K_{\text{str}}^{\text{eq}} H_{\text{str}}^{-}$. Introducing these expressions into Eq. (S55), we obtain

$$\alpha_{\text{thy}}^{H^-}(C_{\text{str}}/K_{\text{str}}^{\text{eq}} - C_{\text{thy}}/K_{\text{thy}}^{\text{eq}}) = \alpha_{\text{thy}}^{C}(C_{\text{thy}} - C_{\text{str}}) + \sigma_{\text{thy}}^{C}(C_{\text{thy}} - C_{\text{pyr}}),\ \text{and} \tag{S56a}$$

$$\sigma_{\text{thy}}^{C}(C_{\text{thy}} - C_{\text{pyr}}) = \frac{C_{\text{pyr}}}{C_{\text{pyr}} + K_{\text{m}}^{\text{eff}}} = \sigma_{\text{chl}}^{C}(C_{\text{cyt}} - C_{\text{str}}). \tag{S56b}$$

Note that there are no undetermined parameters in Eq. (S56). Thus, the model predicts the ability to concentrate $CO_2$ (e.g., NCFF) under different $CO_2$ conditions if LCIB is the only inorganic carbon uptake system (Fig. S16A). Note also that, consistent with full simulations (Fig. 3), NCFF is indeed larger than 0.5 under air-level $CO_2$ (10 μM cytosolic) and below 0.5 under very low $CO_2$ (1 μM cytosolic).

**Extreme 2: no LCIB, fast $HCO_3^-$ transport across the chloroplast envelope.** In this limit, $\beta_{\text{LCIB}} = 0$ and $\sigma_{\text{chl}}^{H^-} \to \infty$, and thus stroma $HCO_3^-$ level is determined by $H_{\text{str}}^{-} = \gamma^{-1} H_{\text{cyt}}^{-} = \gamma^{-1} 10\ C_{\text{cyt}}$. Introducing these expressions into Eq. (S55), we obtain

$$\alpha_{\text{thy}}^{H^-}(\gamma^{-1}10\ C_{\text{cyt}} - C_{\text{thy}}/K_{\text{thy}}^{\text{eq}}) = \alpha_{\text{thy}}^{C}(C_{\text{thy}} - C_{\text{str}}) + \sigma_{\text{thy}}^{C}(C_{\text{thy}} - C_{\text{pyr}}), \tag{S57a}$$

$$\sigma_{\text{thy}}^{C}(C_{\text{thy}} - C_{\text{pyr}}) = \frac{C_{\text{pyr}}}{C_{\text{pyr}} + K_{\text{m}}^{\text{eff}}},\ \text{and} \tag{S57b}$$

$$\alpha_{\text{thy}}^{C}(C_{\text{thy}} - C_{\text{str}}) = \sigma_{\text{chl}}^{C}(C_{\text{str}} - C_{\text{cyt}}). \tag{S57c}$$

As shown in Fig. S16B, active $HCO_3^-$ transporters of reversibility $\gamma = 0.2$ behave similarly to the passive $CO_2$ uptake strategy exmploying fast LCIB (see **Extreme 1** above), and active $HCO_3^-$ transporters of smaller reversibility (e.g., $\gamma \leqslant 0.05$) enable NCFF $> 0.5$ under both air-level $CO_2$ and very low $CO_2$ conditions.

#### C. Feasible CCM strategies under varying $CO_2$ conditions

We next consider an arbitrary CCM strategy in the well-mixed compartment model. Similar to the full reaction-diffusion model, we can compute the overall reaction fluxes of the CCM enzymes as well as the fluxes of inorganic carbon transport between compartments to assess the performance, i.e., NCFF and energy cost, of a given CCM strategy (Figs. S17A and B). The results of the two CCM strategies are in line with Fig. 3. In addition, Figs. S17A and B show that, in the presence of fast $HCO_3^-$ pumping across the chloroplast transport, increasing LCIB activity only minimally affects NCFF but reduces the energy cost by recycling outward diffusing $CO_2$.

Solving Eq. (S55) under different external $CO_2$ levels yields the lower bound $C_{\text{cyt}}^{*}$ below which NCFF $< 0.5$ (Fig. S17C and D). For channel transport of $HCO_3^-$ across the chloroplast envelope ($\gamma = 1$), the lowest $C_{\text{cyt}}^{*}$ is achieved by employing LCIB to catalyze passive $CO_2$ uptake with no investment in $HCO_3^-$ transport across the chloroplast envelope (termed “the passive $CO_2$ uptake strategy”, see Sec. III D). For the CCM to function at even lower $CO_2$

levels, active $HCO_3^-$ pumping ($\gamma < 0.2$) across the chloroplast envelope needs to be turned on. This suggests that the cell might activate different sets of CCM components depending on the environmental $CO_2$ conditions. The least costly functional CCM strategy of the well-mixed compartment model under varying external $CO_2$ conditions is found to be the sequential activation of LCIB followed by chloroplast envelope $HCO_3^-$ pumps (Fig. S17E), which is also the case in the full simulation (Fig. S18).

##### D. Limits of the passive $CO_2$ uptake strategy

What are the relevant biophysical limits for the CCM? For the passive $CO_2$ uptake strategy, where there is no $HCO_3^-$ transport across the chloroplast envelope, the efficacy of the CCM is limited by the influx of $CO_2$ which depends on the external $CO_2$ concentration and the size of the chloroplast. For such strategy, the overall flux balance of carbon takes the same form as Eq. (S56b), since the influx must be in the form of $CO_2$ and must be equal to the total $CO_2$ fixation flux. Since $C_{\rm str}$ cannot be negative, we obtain

$$\mathrm{NCFF} = \frac{C_{\rm pyr}}{C_{\rm pyr} + K_{\rm m}^{\rm eff}} = \sigma_{\rm chl}^{C}(C_{\rm cyt} - C_{\rm str}) \leqslant \sigma_{\rm chl}^{C} C_{\rm cyt}, \tag{S58}$$

which yields the lowest external $CO_2$ concentration, $C_{\rm cyt}^* \sim (\sigma_{\rm chl}^C)^{-1} = 1.5$ μM, above which Rubisco could in principle reach saturation (see Fig. S16A).

In the rough estimation above, we have assumed that the total $CO_2$ influx is not diffusion-limited, but rather limited by the chloroplast membrane permeability to $CO_2$ ($\sigma_{\rm chl}^C$ or $\kappa^C$). However, the passive $CO_2$ uptake strategy will fail at a higher $CO_2$ concentration if the size of the chloroplast is increased — eventually $CO_2$ absorption becomes diffusion-limited with a flux $\propto R_{\rm chlor}$ while the saturating $CO_2$ fixation flux scales as $R_{\rm chlor}^3$. Thus, we wonder whether this diffusion-limited absorption sets a relevant limit to the size of the chloroplast, assuming that the Chlamydomonas cell employs such a CCM strategy under air-level $CO_2$ (10 μM cytosolic).

To calculate $CO_2$ absorption flux by a chloroplast of arbitrary size, we consider free diffusion of $CO_2$ outside the chloroplast, described by $r^{-2}\partial_r(r^2\partial_r C) = 0$ in which $C$ denotes the concentration of $CO_2$ and $r$ denotes the distance from the chloroplast center. Assuming complete absorption, the boundary conditions are given by $D^C(\partial_r C)|_{r=R_{\rm chlor}} = \kappa^C C|_{r=R_{\rm chlor}}$ and $C|_{r=\infty} = C_{\rm cyt}$. Solving the differential equation yields $C(r) = C_{\rm cyt} - \frac{C_{\rm cyt}}{1+D^C/(\kappa^C R_{\rm chlor})}\frac{R_{\rm chlor}}{r}$. Thus, the total $CO_2$ uptake flux is given by $4\pi R_{\rm chlor}^2 \kappa^C C|_{r=R_{\rm chlor}} = \frac{4\pi R_{\rm chlor} D^C C_{\rm cyt}}{1+D^C/(\kappa^C R_{\rm chlor})}$. Setting this equal to the saturating Rubisco fixation flux $\frac{4}{3}\pi R_{\rm pyr}^3 V_{\rm max,Rbc}^C = \frac{4}{3}\pi(\chi R_{\rm chlor})^3 V_{\rm max,Rbc}^C$ yields the upper size limit $R_{\rm chlor}^*$ below which the passive $CO_2$ uptake strategy is functional. Here, we have assumed that the pyrenoid varies its size in proportion to the chloroplast with a proportionality constant $\chi$, while maintaining the same Rubisco density. Plugging in $D^C = 1.88 \times 10^3$ μm/s, $C_{\rm cyt} = 10$ μM, $\kappa^C = 300$ μm/s and $V_{\rm max,Rbc}^C = 15$ mM/s (Table S1), we obtain $R_{\rm chlor}^* = 9.08$ μm for $\chi = 0.3$, $R_{\rm chlor}^* = 5.15$ μm for $\chi = 0.4$, and $R_{\rm chlor}^* = 3.18$ μm for $\chi = 0.5$. These values are consistent with the typical dimensions of algal chloroplasts, which are typically smaller than 5 μm, suggesting a relevant biophysical size limit imposed by $CO_2$ uptake.

#### IV. WELL-MIXED COMPARTMENT MODEL WITH THYLAKOID STACKS

As seen in Figs. 2 and S7, a model with thylakoid stacks slowing inorganic carbon diffusion in the stroma behaves qualitatively differently from a model with a starch sheath. In particular, the former shows a nonmonotonic dependence of NCFF on LCIB activity, while in the latter model NCFF increases monotonically with LCIB activity, plateauing at high LCIB rates. To better understand the differences, we aim here to analyze the diffusion barrier formed by the thylakoid stacks using the framework of the well-mixed compartment model as above (Sec. III A). In Sec. IV A, we derive the typical diffusion length scale of $CO_2$ in the stroma, which depends on LCIB activity. In Sec. IV B, we derive the governing equations of a simplified model. In Sec. IV C, we discuss the consequences of the thylakoid stacks acting as a diffusion barrier.

##### A. Diffusion length scale in the stroma

Consider the reaction-diffusion kinetics of $CO_2$ in the stroma. For simplicity, we ignore the effect of the thylakoid tubules since they have a negligible volume fraction compared to the stroma (Fig. S2). We further approximate the reaction kinetics of LCIB to be linear and ignore the spontaneous interconversion between $CO_2$ and $HCO_3^-$. Thus, Eq. (S4) becomes

$$\frac{a_\beta^2}{r^2}\frac{\partial}{\partial r}\left(r^2\frac{\partial C_{\rm str}}{\partial r}\right) - \left(C_{\rm str} - K^{\rm eq}_{\rm str}H^-_{\rm str}\right) = 0, \tag{S59}$$

where $a_\beta = \left(\frac{D^C_{\rm eff}}{V^C_{\rm max,LCIB}/K^C_{\rm m}}\right)^{1/2}$ denotes the typical distance over which a $CO_2$ molecule can diffuse in the stroma before it is converted to $HCO_3^-$ by LCIB. Assuming that $H^-_{\rm str}$ is a constant in the stroma, we consider the boundary condition problem with $C_{\rm str}(r = R_{\rm pyr}) = C_*$ and $C_{\rm str}(r = R_{\rm chlor}) = C_0$ and compute the diffusive flux of $CO_2$ across $r = R_{\rm pyr}$. In general, one can solve the exact concentration profile from Eq. (S59) and use the expression $-D^C_{\rm eff}\frac{\partial C}{\partial r}$ to compute the flux. Here, for simplicity, we focus on the limit where either the stroma domain is large enough or LCIB is fast enough that $C_0 \approx K^{\rm eq}_{\rm str}H^-_{\rm str}$ (see Sec. IV C for justification). In this limit, we obtain

$$C_{\rm str}(r) = C_0 - (C_* - C_0)\frac{R_{\rm pyr}}{r}\frac{\sinh(r/a_\beta - R_{\rm chlor}/a_\beta)}{\sinh(\Delta R_{\rm str}/a_\beta)}, \tag{S60}$$

where $\Delta R_{\rm str} = R_{\rm chlor} - R_{\rm pyr}$. Thus, the diffusive flux of $CO_2$ across $r = R_{\rm pyr}$ can be written as $-D^C_{\rm eff}\frac{\partial C_{\rm str}}{\partial r}|_{r=R_{\rm pyr}} = D^C_{\rm eff}\frac{C_*-C_0}{L_{\rm str}}$ where the diffusion length scale in the stroma $L_{\rm str}$ is given by

$$L^{-1}_{\rm str} = R^{-1}_{\rm pyr} + a^{-1}_\beta\coth(\Delta R_{\rm str}/a_\beta). \tag{S61}$$

Note that $L_{\rm str}$ is dependent on LCIB activity via $a_\beta$: when LCIB activity is low, i.e., $a_\beta \to \infty$, $L_{\rm str} = \Delta R_{\rm str}R_{\rm pyr}/R_{\rm chlor}$ which is determined solely by the chloroplast geometry; when LCIB activity is high, i.e., $a_\beta \to 0$, $L_{\rm str} \approx a_\beta$ which is determined solely by LCIB activity.

##### B. Flux-balance equations

Here, we derive the algebraic equations of flux-balance conditions in a chloroplast with thylakoid stacks, similar to the derivations of Eq. (S55) in Sec. III A. Because the diffusion coefficient of inorganic carbon in the stroma is smaller than that in the pyrenoid matrix due to the modeled thylakoid stacks, the concentration profiles are flatter in the matrix (Fig. S6A). Thus, we approximate the $CO_2$ concentration in the pyrenoid matrix as constant, denoted by a single number $C_{\rm pyr}$. Similarly, we assume that the $CO_2$ concentration in the thylakoid tubules inside the pyrenoid can be approximated by another constant, $C_{\rm thy}$. $CO_2$ concentrations in the thylakoids outside the pyrenoid and in the stroma decrease from $C_{\rm thy}$ and $C_{\rm pyr}$, respectively, with increasing radius, and they approach the same plateau $C_{\rm str}$ near the chloroplast envelope. For simplicity, we assume that the concentrations of $HCO_3^-$ in all the compartments can be approximated by the same constant $H^-$. [6] Below, we consider the passive $CO_2$ uptake strategy where there

[6] This becomes a good approximation when the $HCO_3^-$ transport rate across the thylakoid membrane $\kappa^{H^-}_{\rm thy}$ is fast. For $\kappa^{H^-}_{\rm thy} \sim \kappa^C$, one could introduce the same set of variables as the $CO_2$ concentrations, i.e., $H^-_{\rm pyr}$, $H^-_{\rm thy}$, $H^-_{\rm str}$, and consider flux-balance equations similar to Eqs. (S64) and (S65). However, the assumptions that $\kappa^{H^-}_{\rm thy}$ is fast is not essential to understanding the nonmonotonic dependence of NCFF on LCIB activity.

is no $HCO_3^-$ transport across the chloroplast envelope.

The overall flux balance of $HCO_3^-$ in the chloroplast is given by

$$I[R_{\rm pyr}, R_{\rm chlor}, 1, 1-f_{\rm v}]\frac{V^C_{\rm max,LCIB}}{K^C_{\rm m}}(C_{\rm str} - K^{\rm eq}_{\rm str}H^-) = I[0, R_{\rm pyr}, 1, f_{\rm v}]\frac{V^C_{\rm max,CAH3}}{K^C_{\rm m}}(K^{\rm eq}_{\rm thy}H^- - C_{\rm thy}), \tag{S62}$$

where the left-hand side and the right-hand side of the equation denote the total $HCO_3^-$ production by LCIB and the total $HCO_3^-$ consumption by CAH3, respectively. Here, we have again assumed linear kinetics of LCIB and CAH3 (see also Sec. III A).

The overall flux balance of $CO_2$ in the chloroplast dictates that the flux of $CO_2$ diffusing into the chloroplast is equal to the flux of $CO_2$ consumed by Rubisco, i.e.,

$$4\pi R^2_{\rm chlor}\kappa^C(C_{\rm cyt} - C_{\rm str}) = I[0, R_{\rm pyr}, 1, 1-f_{\rm v}]V^C_{\rm max,Rbc}\frac{C_{\rm pyr}}{C_{\rm pyr} + K^{\rm eff}_{\rm m}}. \tag{S63}$$

The flux balance of $CO_2$ in the pyrenoid matrix yields

$$\Delta S_{\rm thy}\kappa^C(C_{\rm thy} - C_{\rm pyr}) = I[0, R_{\rm pyr}, 1, 1-f_{\rm v}]V^C_{\rm max,Rbc}\frac{C_{\rm pyr}}{C_{\rm pyr} + K^{\rm eff}_{\rm m}} + 4\pi R^2_{\rm pyr}(1 - f_{\rm v}|_{r=R_{\rm pyr}})D^C_{\rm eff}\frac{C_{\rm pyr} - C_{\rm str}}{L_{\rm str}}, \tag{S64}$$

since the $CO_2$ molecules entering the matrix from the thylakoid lumen (denoted by the left-hand side) must either be fixed by Rubisco (denoted by first term on the right-hand side) or diffuse back into the stroma (denoted by the second term on the right-hand side). Here, we have used results from Sec. IV A to express the flux of $CO_2$ diffusing from the pyrenoid matrix to the stroma. The diffusion length scale in the stroma $L_{\rm str}$ is given by Eq. (S61).

Finally, the flux balance of $CO_2$ in the thylakoid tubules inside the pyrenoid yields

$$I[0, R_{\rm pyr}, 1, f_{\rm v}]\frac{V^C_{\rm max,CAH3}}{K^C_{\rm m}}(K^{\rm eq}_{\rm thy}H^- - C_{\rm thy}) = \Delta S_{\rm thy}\kappa^C(C_{\rm thy} - C_{\rm pyr}) + D^C(\pi a^2_{\rm tub}/L_{\rm thy})N_{\rm tub}(C_{\rm thy} - C_{\rm str}), \tag{S65}$$

since $CO_2$ produced by CAH3 (denoted by the left-hand side) must either diffuse into the pyrenoid matrix from the thylakoid lumen (denoted by first term on the right-hand side) or diffuse away via the thylakoid tubules (denoted by second term on the right-hand side; see also Eq. (S51)).

Equations (S62–S65) can be written as

$$\beta_{\rm LCIB}(C_{\rm str} - K^{\rm eq}_{\rm str}H^-) = \beta_{\rm CAH3}(K^{\rm eq}_{\rm thy}H^- - C_{\rm thy}), \tag{S66a}$$

$$\sigma^C_{\rm chl}(C_{\rm cyt} - C_{\rm str}) = \frac{C_{\rm pyr}}{C_{\rm pyr} + K^{\rm eff}_{\rm m}}, \tag{S66b}$$

$$\sigma^C_{\rm thy}(C_{\rm thy} - C_{\rm pyr}) = \frac{C_{\rm pyr}}{C_{\rm pyr} + K^{\rm eff}_{\rm m}} + \alpha^C_{\rm str}(C_{\rm pyr} - C_{\rm str}), \tag{S66c}$$

$$\beta_{\rm CAH3}(K^{\rm eq}_{\rm thy}H^- - C_{\rm thy}) = \sigma^C_{\rm chl}(C_{\rm thy} - C_{\rm pyr}) + \alpha^C_{\rm str}(C_{\rm pyr} - C_{\rm str}), \tag{S66d}$$

where we have again grouped the parameters into the following inverse concentrations (see also Eqs. (S55a–S55d) and the surrounding text for definitions). Here, we define $\beta_{\rm CAH3} \equiv \frac{I[0,R_{\rm pyr},1,f_{\rm v}]V^C_{\rm max,CAH3}/K^C_{\rm m}}{I[0,R_{\rm pyr},1,1-f_{\rm v}]V^C_{\rm max,Rbc}} = 43.0\ {\rm mM}^{-1}$ and $\alpha^C_{\rm str} \equiv \frac{4\pi R^2_{\rm pyr}(1-f_{\rm v}|_{r=R_{\rm pyr}})D^C_{\rm eff}/L_{\rm str}}{I[0,R_{\rm pyr},1,1-f_{\rm v}]V^C_{\rm max,Rbc}}$, which is dependent on LCIB activity. Although much simplified, Eqs. (S66a – S66d) capture many key features of the full simulation. When LCIB activity is low, CAH3 is the dominant carbonic anhydrase in the chloroplast. Since CAH3 works in the thylakoid tubules at pH 6, the $HCO_3^-$ to $CO_2$ ratios in both the stroma and inner thylakoids are close to the equilibrium ratio at pH 6 (Fig. S9A). As LCIB activity increases, more $CO_2$ diffusing into the chloroplast is converted to $HCO_3^-$ by LCIB. Thus, the concentration of stromal $HCO_3^-$ increases and the $HCO_3^-$ to $CO_2$ ratio in the stroma approaches the equilibrium ratio at pH 8 (Figs. S9A and B). Consequently, more $HCO_3^-$ can diffuse into the thylakoid tubules and become available for the conversion of $CO_2$ by CAH3, leading to an increase of the $CO_2$ concentration in the thylakoid tubules inside the pyrenoid (Fig. S9B). Eq. (S61) also indicates that the transport coefficient $\alpha^C_{\rm str} \propto L^{-1}_{\rm str}$ increases with LCIB activity. Specifically, in the limit of fast LCIB, $\alpha^C_{\rm str} \sim L^{-1}_{\rm str} \approx a^{-1}_\beta \sim (V^C_{\rm max,LCIB}/K^C_{\rm m})^{1/2}$ (Sec. IV A and Fig. S9C).

##### C. Results and discussion

**Optimal LCIB activity.** To gain more analytical insight into the nonmonotonic dependence of NCFF on LCIB when thylakoid stacks are employed, we ignore the only nonlinear term $\frac{C_{\text{pyr}}}{C_{\text{pyr}}+K_{\text{m}}^{\text{eff}}}$ in Eq. (S66) and proceed with $C_{\text{str}} = C_{\text{cyt}}$. Note that this is a reasonable simplification since the neglected term is smaller than 1 while the coefficients $\alpha$, $\beta$ and $\sigma$ are much larger than 1 (Table. S3). Solving the simplified linear equations, we obtain

$$C_{\text{pyr}} = C_{\text{cyt}} + \frac{\sigma_{\text{thy}}^C}{\sigma_{\text{thy}}^C + \alpha_{\text{str}}^C}\big(C_{\text{thy}} - C_{\text{cyt}}\big), \tag{S67}$$

where $C_{\text{thy}}$ is given by

$$C_{\text{thy}} = C_{\text{cyt}} + \frac{\beta_{\text{CAH3}}}{\beta_{\text{CAH3}} + \alpha}\big(K_{\text{thy}}^{\text{eq}} H^- - C_{\text{cyt}}\big) \text{ in which } H^- = \frac{\beta_{\text{LCIB}} + \alpha^*}{K_{\text{str}}^{\text{eq}}\beta_{\text{LCIB}} + K_{\text{thy}}^{\text{eq}}\alpha^*} C_{\text{cyt}}. \tag{S68}$$

To simplify the expression, we have defined $\alpha \equiv \alpha_{\text{thy}}^C + \frac{\alpha_{\text{str}}^C \sigma_{\text{thy}}^C}{\alpha_{\text{str}}^C + \sigma_{\text{thy}}^C}$ and $\alpha^* \equiv \frac{\alpha\beta_{\text{CAH3}}}{\alpha+\beta_{\text{CAH3}}}$. The two competing effects of increasing LCIB activity on concentrating $CO_2$ are apparent in Eqs. (S67) and (S68). Since $K_{\text{str}}^{\text{eq}} \ll K_{\text{thy}}^{\text{eq}}$, Eq. (S68) indicates the benefit of increasing LCIB activity ($\beta_{\text{LCIB}}$): $HCO_3^-$ concentration in the chloroplast increases, leading to a high level of $CO_2$ ($C_{\text{thy}}$) in the thylakoid tubules inside the pyrenoid. The concentration of $CO_2$ is decreased as it diffuses from the thylakoid lumen into the pyrenoid matrix and then back out into the stroma, and the concentration drop from one compartment to another is inversely proportional to the transport coefficients $\alpha_{\text{str}}^C$ and $\sigma_{\text{thy}}^C$ (inset of Fig. S9C). Equation (S67) describes the detrimental contribution of increasing LCIB activity: LCIB converts $CO_2$ diffusing out of the pyrenoid matrix to $HCO_3^-$, leading to an increase of $\alpha_{\text{str}}^C$ while $\sigma_{\text{thy}}^C$ remains constant; thus, the concentration drop from the pyrenoid matrix to the stroma becomes smaller as LCIB activity increases, implying a lower level of $CO_2$ in the matrix. Together, the two competing effects result in the existence of an intermediate LCIB activity at which NCFF and pyrenoid matrix $CO_2$ concentration reach the maximum (Figs. S6E and S9D). This behavior is unique to the model with thylakoid stacks and no starch sheath since the path of $CO_2$ diffusing from the pyrenoid matrix to stroma is blocked when a starch sheath is present.

Introducing parameter values into Eqs. (S67) and (S68) allows us to determine quantitatively where the optimal LCIB activity occurs. For the beneficial contribution Eq. (S68), the factor $\frac{\beta_{\text{LCIB}}+\alpha^*}{K_{\text{str}}^{\text{eq}}\beta_{\text{LCIB}}+K_{\text{thy}}^{\text{eq}}\alpha^*}$ reaches half its maximum when $\beta_{\text{LCIB}} = (\frac{K_{\text{thy}}^{\text{eq}}}{K_{\text{str}}^{\text{eq}}} - 2)\alpha^* = 98\alpha^*$, which corresponds to an LCIB first-order rate constant of $k_{\frac{1}{2}} = 867\ \text{s}^{-1}$. Similarly, for the detrimental contribution Eq. (S67), setting the factor $\frac{\sigma_{\text{thy}}^C}{\sigma_{\text{thy}}^C+\alpha_{\text{str}}^C}$ to 1/2 yields an LCIB first-order rate constant of 902 $\text{s}^{-1}$. These numbers are in good agreement with the optimal LCIB rate of about $10^3\ \text{s}^{-1}$ found in the full simulation (Fig. S9D). Note also that the detrimental contribution becomes important only for high LCIB rates when the beneficial contribution saturates. Thus, in deriving the expression Eq. (S61) for $L_{\text{str}}$, which is directly related to the detrimental contribution, we have made the assumption of fast LCIB (see Sec. IV A).

**Optimal membrane permeability to $CO_2$.** If the permeability to $CO_2$ of thylakoid membranes $\kappa^C$ is the same for both the thylakoid tubules and stacks, Eq. (S67) also suggests the competing effects of increasing $\kappa^C$: when $\kappa^C$ is small, i.e., when $\sigma_{\text{thy}}^C \propto \kappa^C$ is small, it is difficult for the $CO_2$ produced by CAH3 in the thylakoid tubules to diffuse into the pyrenoid matrix, leading to a low level of $CO_2$ therein; when $\kappa^C$ is large, the diffusion coefficient of $CO_2$ in the stroma becomes large (Fig. S5), leading to a large $CO_2$ leakage flux out of the pyrenoid matrix and impairing CCM performance (Fig. S3). Thus, the theory predicts an intermediate $\kappa^C$ that maximizes NCFF or pyrenoid matrix $CO_2$ concentration. This prediction is verified by varying $\kappa^C$ in the reaction-diffusion model described in Sec. I A (Figs. S10A and B). Results obtained from the simplified model Eq. (S66) show good agreement with the full simulation (Figs. S10C – E).

**Localization of LCIB.** In deriving Eq. (S61), we have considered diffuse LCIB in the stroma. Intuitively, moving LCIB away from the boundary between the pyrenoid matrix and stroma will increase the diffusion length scale $L_{\text{str}}$ as defined in Sec. IV A. We wondered whether this would qualitatively affect the nonmonotonic dependence of NCFF on increasing LCIB activity. To address this question, we consider a scenario in which LCIB range from $R_{\text{pyr}} + \delta$ to $R_{\text{chlor}}$ where $\delta$ denotes the radial width of the LCIB-free buffer zone. Free diffusion is described by $r^{-2}\partial_r(r^2\partial_r C_{\text{str}}) = 0$ between $R_{\text{pyr}}$ and $R_{\text{pyr}} + \delta$, and the reaction-diffusion equation $a_\beta^2 r^{-2}\partial_r(r^2\partial_r C_{\text{str}}) - (C_{\text{str}} - C_0) = 0$ applies for $r > R_{\text{pyr}} + \delta$. Here, for the same reason as above, we have focused on the case where LCIB is fast (i.e., $a_\beta$ is small) and we consider boundary conditions $C_{\rm str}(r=R_{\rm pyr})=C_*$ and $C_{\rm str}(r=\infty)=C_0$.[7] Thus, the general solution of $C_{\rm str}$ is given by

$$C_{\rm str}(r)=\begin{cases}C_*+\mathcal{A}(1/r-1/R_{\rm pyr}) & \text{for } R_{\rm pyr}\leqslant r\leqslant R_{\rm pyr}+\delta,\ \text{and}\\ C_0+\mathcal{B}(a_\beta/r)\exp(-r/a_\beta) & \text{for } r\geqslant R_{\rm pyr}+\delta,\end{cases}\tag{S69}$$

where $\mathcal{A}$ and $\mathcal{B}$ are constants determined by the boundary conditions at $r=R_{\rm pyr}+\delta$, i.e., the concentration $C_{\rm str}$ and its derivative must be continuous, which yields

$$C_*-\mathcal{A}\frac{\delta}{(R_{\rm pyr}+\delta)R_{\rm pyr}}=C_0+\mathcal{B}\frac{a_\beta}{R_{\rm pyr}+\delta}\exp\Big(-\frac{R_{\rm pyr}+\delta}{a_\beta}\Big),\ \text{and}\tag{S70a}$$

$$\mathcal{A}(R_{\rm pyr}+\delta)^{-2}=\mathcal{B}(R_{\rm pyr}+\delta)^{-1}\exp\Big(-\frac{R_{\rm pyr}+\delta}{a_\beta}\Big)\Big(1+\frac{a_\beta}{R_{\rm pyr}+\delta}\Big).\tag{S70b}$$

Solving Eq. (S70), we obtain $\mathcal{A}=(C_*-C_0)\big[R_{\rm pyr}^{-1}-(R_{\rm pyr}+\delta+a_\beta)^{-1}\big]^{-1}$. The diffusive flux of $CO_2$ across $r=R_{\rm pyr}$ is given by $-D_{\rm eff}^C\partial_r C_{\rm str}|_{r=R_{\rm pyr}}=-D_{\rm eff}^C\mathcal{A}/R_{\rm pyr}^2$. Introducing the expression for $\mathcal{A}$ and rewriting the flux in terms of $L_{\rm str}$ as $D_{\rm eff}^C\frac{C_*-C_0}{L_{\rm str}}$, we obtain

$$L_{\rm str}^{-1}=R_{\rm pyr}^{-1}+(\delta+a_\beta)^{-1}.\tag{S71}$$

Recall that when LCIB is diffuse, $L_{\rm str}\approx a_\beta$ is solely determined by LCIB activity and it approaches 0 in the fast LCIB limit. In contrast, when LCIB is moved away from the pyrenoid by a distance $\delta$, $L_{\rm str}\approx\delta R_{\rm pyr}/(\delta+R_{\rm pyr})$ is determined by the geometry and it approaches some finite value in the fast LCIB limit. Denote by $a_{\frac{1}{2}}=(D_{\rm eff}^C/k_{\frac{1}{2}})^{1/2}$ where $k_{\frac{1}{2}}$ is again the first-order rate constant of LCIB above which the beneficial contribution starts to saturate (see **Optimal LCIB activity** above). Equation (S71) also indicates that when $\delta\gtrsim a_{\frac{1}{2}}$, the detrimental contribution of LCIB at high activity will be masked by the LCIB-free slow diffusion buffer zone. Thus, this analysis predicts a critical width of $\delta^*\approx a_{\frac{1}{2}}\sim(D_{\rm eff}^C)^{1/2}$, below which NCFF has nonmonotonic dependence on LCIB activity and above which NCFF increases monotonically with LCIB activity. This is indeed the case in the full simulation and the dependence of $\delta^*$ on the stroma diffusivity $D_{\rm eff}^C$ is verified (Fig. S11).

**Estimating parameter values in the full simulation.** To test whether the above calculations in the simplified model also hold true for the reaction-diffusion model (Sec. I A), we estimate the corresponding parameter values in the full simulation as follows. First, the concentrations of $CO_2$ and $HCO_3^-$ are estimated by their average in each compartment in the full simulation (Figs. S9A and B). Second, according to the definitions of the transport coefficients $\alpha_{\rm str}^C$ and $\sigma_{\rm thy}^C$, we divide the diffusive fluxes of $CO_2$ by the (average) concentration difference between compartments to estimate the values of $\alpha_{\rm str}^C$ and $\sigma_{\rm thy}^C$ in the simulation (Figs. S9 and S10). Finally, we estimate $a_{\frac{1}{2}}=(D_{\rm eff}/10^3\ {\rm s}^{-1})^{1/2}$ (Fig. S11) since $k_{\frac{1}{2}}$ is roughly $10^3\ {\rm s}^{-1}$ under various stroma diffusivity conditions.

---

[7] Using this boundary condition at a finite radius $C_{\rm str}(r=R_{\rm chlor})=C_0$ yields $L_{\rm str}=\frac{\delta R_{\rm pyr}}{\delta+R_{\rm pyr}}+\frac{a_\beta}{(1+\delta/R_{\rm pyr})^2}\frac{\sinh(x)}{\cosh(x)+\sinh(x)a_\beta/(R_{\rm pyr}+\delta)}$ where $x=(R_{\rm chlor}-R_{\rm pyr}-\delta)/a_\beta$. We recover Eq. (S71) when $a_\beta\ll R_{\rm chlor}-R_{\rm pyr}-\delta$.

### V. LOCALIZATION AND REGULATION OF KEY CCM ENZYMES

Previous experiments have shown that the localization patterns and the expression or activity levels of key CCM enzymes are regulated as Chlamydomonas cells respond to external $CO_2$ levels [16, 17, 57, 58]. One possibility is that such $CO_2$-dependent regulation contributes to the efficient functioning of the CCM under different $CO_2$ conditions. In this section, we discuss how different localization patterns and activities of key CCM enzymes impact CCM performance in the modeled chloroplast.

#### A. Localization and regulation of CAH3

Previous experiments [17] suggest that CAH3 enzyme activity is upregulated under air-level $CO_2$ and that CAH3 is localized toward the intra-pyrenoid portion of the thylakoid tubules. In our baseline model, CAH3 serves as the immediate source of $CO_2$ that elevates the local $CO_2$ concentration around Rubisco. Thus, we hypothesize that certain activity and localization of CAH3 is needed only when external $CO_2$ concentration is low.

To test this hypothesis, we hold constant the total number of CAH3 and vary its localization. As shown in Figs. S19B, G, and L, localizing CAH3 in the pyrenoid portion of the thylakoids indeed helps to concentrate more $CO_2$ in the vicinity of Rubisco under air-level $CO_2$ (10 μM cytosolic). Furthermore, such a localization pattern also reduces energy costs (Figs. S19C, H, and M) by avoiding futile cycles when CAH3 and LCIB overlap radially. In the case of such an overlap, $CO_2$ produced by CAH3 can diffuse diffuse out of thylakoid tubules in the stroma, where it is then converted to $HCO_3^-$ by LCIB. That $HCO_3^-$ can then diffuse back into the tubules, where it is again converted to $CO_2$ by CAH3. Thus, the energetically least costly CCM strategy that saturates Rubisco with $CO_2$ under air-level $CO_2$ is achieved when CAH3 is localized to the pyrenoid tubules and has a relatively high activity (Figs. S19I and N). Finally, simulations verify that, indeed, as the external $CO_2$ level decreases, high activity and pyrenoid localization of CAH3 become required for a functioning CCM, which is in line with the reported cellular regulation of CAH3 under air-level $CO_2$ [17].

#### B. Localization of LCIB

Under air-level $CO_2$ (10 μM cytosolic) when the passive $CO_2$ uptake strategy is used, for a chloroplast employing thylakoid stacks and no starch sheath, it is favorable for the CCM to localize LCIB toward the chloroplast envelope (Fig. S21; see also Sec. IV C **Localization of LCIB**). In contrast, for a chloroplast employing an impermeable starch sheath, the localization patterns of LCIB minimally impact the performance of the CCM (Fig. S21). *In vivo*, LCIB is diffusely distributed in the Chlamydomonas chloroplast under air-level $CO_2$ [58]. We speculate that the passive $CO_2$ uptake strategy and a strong barrier formed by the starch sheath are employed by Chlamydomonas cells under such conditions.

**LCIB puncta versus LCIB shell.** Under very low $CO_2$ (1 μM cytosolic), LCIB is known to form puncta around the periphery of the starch sheath [57, 58]. This localization pattern has been suggested to contribute to CCM performance [59]. Our model indicates that a punctate localization could be a strategy to avoid loss of inorganic carbon under very low $CO_2$: when LCIB is close to the envelope, $HCO_3^-$ pumped across the chloroplast envelope is converted to $CO_2$, which could diffuse back out of the chloroplast immediately (Fig. 5). Furthermore, since the major role of LCIB is to recycle outward diffusing $CO_2$ and prevent leakage, we wonder whether forming LCIB puncta around the thylakoid tubules could benefit the CCM more than forming a diffuse shell around the pyrenoid.

To estimate the difference between these two LCIB localization patterns, we consider an infinite cylindrical tubule of radius $a_{\text{tub}}$ with a $CO_2$ concentration of $C_i$ inside the tubule. We consider a case where $CO_2$ simply undergoes free diffusion outside the tubule, which is described by $\rho^{-1}\partial_\rho(\rho\partial_\rho C) = 0$ ($\rho \geqslant a_{\text{tub}}$) where $\rho$ denotes the perpendicular distance from the central axis of the tubule. We denote by $\kappa^C$ the membrane permeability to $CO_2$ on the side of the tubule. Thus, the boundary condition at $\rho = a_{\text{tub}}$ is given by $\kappa^C(C_i - C|_{\rho=a_{\text{tub}}}) = -D^C(\partial_\rho C)|_{\rho=a_{\text{tub}}}$ where $D^C$ denotes the diffusion coefficient in the direction perpendicular to the axis of the tubule. We set the boundary condition away from the tubule to be $C|_{\rho=L} = C_\infty$. Here, we choose $L \approx 1$ μm to be the typical spacing between thylakoid tubules in the stroma, and we denote by $C_\infty$ the concentration of $CO_2$ in the bulk stroma away from the thylakoid tubules. Thus, we obtain

$$C(\rho) = C_\infty - (C_i - C_\infty)\frac{\ln(\rho/L)}{\ln(L/a_{\text{tub}}) + D^C/(\kappa^C a_{\text{tub}})}. \qquad \text{(S72)}$$

Introducing numbers listed in Table S1, we find that

$$\frac{C|_{\rho=a_{\rm tub}} - C_\infty}{C_i - C_\infty} = \frac{\ln(L/a_{\rm tub})}{\ln(L/a_{\rm tub}) + D^C/(\kappa^C a_{\rm tub})} \approx 2.3\%. \tag{S73}$$

This suggests that the concentration gradient of $CO_2$ in the stroma from near to away from the thylakoid tubules is negligible. In this regard, the benefit of forming LCIB puncta around the thylakoid tubules to potentially capture $CO_2$ at a higher concentration is marginal compared to localizing LCIB in a shell around the pyrenoid.

#### VI. SUPPLEMENTARY DISCUSSION

We have focused on the reaction and transport of inorganic carbon molecules, namely, $CO_2$, $HCO_3^-$ and $H_2CO_3$, and we have not explicitly considered protons. *In vivo*, Rubisco catalyzes the addition of $CO_2$ and $H_2O$ to ribulose-1,5-bisphosphate (RuBP) to yield two molecules of 3-phosphoglycerate (3PGA) and two protons for every $CO_2$ fixed. In Sec. VI A, we will discuss where and how these protons could be transported out of the pyrenoid matrix. In Sec. VI D, we further discuss the effect of forming an LCIB-Rubisco condensate in which the protons produced during $CO_2$ fixation could participate in the reaction $HCO_3^- + H^+ \rightarrow CO_2 + H_2O$ catalyzed by LCIB, and the produced $CO_2$ could in turn feed Rubisco $CO_2$ fixation.

##### A. Proton transport

Consider a scenario where the CCM is functional and Rubisco $CO_2$ fixation flux is close to its maximum $J^C_{\text{max}} \approx 4\pi R^3_{\text{pyr}} V^C_{\text{max,Rbc}}/3$. Introducing $R_{\text{pyr}} \approx 1$ μm and $V^C_{\text{max,Rbc}} = 15$ mM/s, we obtain the total proton production flux to be $2J^C_{\text{max}} \approx 1.3 \times 10^{-16}$ mol/s. At steady state, the flux balance of protons in the pyrenoid matrix dictates that the overall proton efflux should be equal to $2J^C_{\text{max}}$. We wonder whether diffusion of free protons could account for this efflux.

One possibility is that free protons could diffuse into the thylakoid tubules in the pyrenoid where they bind to $HCO_3^-$ to produce $CO_2$. To estimate the flux across the thylakoid membranes, we consider the diffusion-absorption process around a cylindrical tubule of radius $a_{\text{tub}}$, similar to the one discussed in Sec. III D. Free diffusion of protons outside the tubule is described by $\rho^{-1}\partial_\rho(\rho\partial_\rho P) = 0$ where $\rho$ denotes the perpendicular distance from the central axis of the tubule and $P$ denotes the proton concentration. The boundary conditions are given by (1) $D^P(\partial_\rho P)|_{\rho=a_{\text{tub}}} = \kappa^P P|_{\rho=a_{\text{tub}}}$, in which $D^P$ denotes the diffusion coefficient of protons and $\kappa^P$ denotes the rate of potential proton transport across the thylakoid membrane, and (2) $P|_{\rho=L} = P_\infty$, where $L$ denotes the typical spacing between thylakoid tubules and $P_\infty$ denotes the bulk proton concentration in the pyrenoid matrix. Solving the diffusion-absorption problem yields $P(\rho) = P_\infty\big[1 + \frac{\ln(\rho/L)}{D^P/(\kappa^P a_{\text{tub}})+\ln(L/a_{\text{tub}})}\big]$. Thus, the total transport flux by diffusion is given by $\Delta S_{\text{thy}} D^P(\partial_\rho P)|_{\rho=a_{\text{tub}}} = \Delta S_{\text{thy}}\kappa^P P_\infty \frac{D^P/a_{\text{tub}}}{D^P/a_{\text{tub}}+\kappa^P\ln(L/a_{\text{tub}})}$ where $\Delta S_{\text{thy}}$ denotes the total surface area of the thylakoid membranes inside the pyrenoid. Note that when diffusion is fast, i.e., $D^P/a_{\text{tub}} \gg \kappa^P$, the absorption flux is given by $J^P = \Delta S_{\text{thy}}\kappa^P P_\infty$, and when absorption is diffusion limited, i.e., $D^P/a_{\text{tub}} \ll \kappa^P$, the maximum absorption flux is given by $J^P_{\text{max}} = \Delta S_{\text{thy}} P_\infty \frac{D^P/a_{\text{tub}}}{\ln(L/a_{\text{tub}})}$. Introducing $\Delta S_{\text{thy}} = 8.5$ μm$^2$ (Tables S1 and S3), $P_\infty = 10^{-8}$ M [2], $D^P = 10^4$ μm$^2$/s [60], $a_{\text{tub}} = 0.05$ μm (Table S1), and $L = 10\ a_{\text{tub}}$ into the expression, we obtain $J^P_{\text{max}} = 0.074 \times 10^{-16}$ mol/s, which is one order of magnitude smaller than the proton production flux. In other words, the pH of the pyrenoid matrix would have to be lower than 6.75, i.e., $P_\infty > 1.76 \times 10^{-7}$ M, in order for the protons produced during $CO_2$ fixation to enter the thylakoid tubules as free protons.

Another possibility is that free protons could diffuse into the stroma, where they are then pumped back into the thylakoid lumen by Photosystems I and II [50, 61, 62]. Assuming that the starch sheath does not serve as a barrier to proton diffusion, we can estimate the maximum efflux from the Burg-Purcell limit [63] (see also Sec. III D), which yields $J^P_{\text{max}} = 4\pi R_{\text{pyr}} D^P \Delta P$. Here, $\Delta P$ denotes the concentration difference of protons between the pyrenoid matrix and the stroma. Thus, $\Delta P$ must be greater than $\frac{J^P_{\text{max}}}{4\pi R_{\text{pyr}} D^P} = \frac{1.3\times10^{-16}\ \text{mol/s}}{4\pi\times(1\ \mu\text{m})\times10^4\ \mu\text{m}^2/\text{s}} = 1.03 \times 10^{-6}$ M if protons are removed from the pyrenoid matrix by free diffusion outward. This clearly disagrees with previous measurements showing that the pH values of both the pyrenoid matrix and stroma are about 8 [2].

The above calculations suggest that free proton concentrations are extremely low at the measured physiological pH values, so efficient transport of protons would require proton carriers; they could exist at higher concentrations in the pyrenoid matrix. A recent study has also highlighted this point and suggests that protonation of RuBP and PGA could play an important role in buffering pH [64].

Since the reactions catalyzed by CAH3 and LCIB also involve protons, similar questions arise about the transport of protons consumed/produced by the carbonic anhydrases. We speculate that proton carriers such as RuBP and 3PGA are involved in these processes. As an example, given its pKa of 6.7, RuBP will predominantly exist in the form of $\text{RuBP}^{4-}$ in the stroma where the pH is 8. If there are RuBP transporters or channels specific to $\text{RuBP}^{4-}$ on the thylakoid membranes, this form could enter the thylakoid lumen (pH 6) and pick up a proton to become $\text{RuBP}^{3-}$, which would then be trapped in the thylakoids. From there, $\text{RuBP}^{3-}$ could diffuse toward the pyrenoid carrying its proton. If this proton were then consumed in converting $HCO_3^-$ to $CO_2$ in the tubules, $\text{RuBP}^{4-}$, the substrate for Rubisco's carboxylation reaction, would be left to diffuse into the pyrenoid matrix. Understanding the mechanism underlying proton tramsport will be an important topic for future experimental studies, and will help to validate the hypotheses presented here and elsewhere.

##### B. Effect of different cytosolic Ci compositions

For simplicity, our model assumes a constant ratio of $CO_2$ to $HCO_3^-$ in the cytosol, corresponding to the equilibrium ratio at pH 7.1 (see Sec. I). However, there is no known CA in the Chlamydomonas cytosol that could catalyze the otherwise slow interconversion between $CO_2$ and $HCO_3^-$. There is a periplasmic CA (CAH1), although mutants lacking this enzyme have no significant growth defect [65]. In addition, Chlamydomonas is known to employ transporters at the cell membrane to import Ci from the external environment to the cytosol [27, 66]. Thus, to explore the effect of different cytosolic Ci compositions on the efficacy of the Chlamydomonas CCM, we varied $C_{\mathrm{cyt}}$ and $H^-_{\mathrm{cyt}}$ independently, assuming that $HCO_3^-$ and $H_2CO_3$ remain in equilibrium at the cytosolic pH 7.1, and that the pH values of the different chloroplast compartments maintain their usual values under all conditions.

As shown in Fig. S25, the efficacy of the CCM in general increases with the the total concentrations of Ci in the cytosol. Notably, for a given total cytosolic Ci concentration, the efficacy of the two Ci uptake strategies varies with the ratio $K_{C/H^-}$ of $CO_2$ to $HCO_3^-$ in the cytosol, or the equivalent cytosolic pH, $\mathrm{pH}^{\mathrm{eqv}}_{\mathrm{cyt}}$, defined by $K_{C/H^-} \equiv 10^{\mathrm{pK_{eff}} - \mathrm{pH}^{\mathrm{eqv}}_{\mathrm{cyt}}}$. In particular, the efficacy of the passive $CO_2$ uptake strategy using LCIB decreases with increasing $\mathrm{pH}^{\mathrm{eqv}}_{\mathrm{cyt}}$ while the active $HCO_3^-$ pumping strategy shows the opposite behavior. Future research could explore whether Chlamydomonas can coordinate the composition of the cytosolic pool of Ci with the Ci uptake strategies of the chloroplast.

##### C. Thylakoid morphologies

In our model of the Chlamydomonas CCM, the thylakoid tubules that traverse the pyrenoid and converge in the pyrenoid center are responsible for delivering stromal $HCO_3^-$ to the pyrenoid, where this $HCO_3^-$ can be converted to $CO_2$ by CAH3. However, this particular architecture of the thylakoid tubules is not necessary. Based on the key insights yielded from our model, we hypothesize that other thylakoid morphologies could also support the functioning of an efficient CCM, as long as (1) $HCO_3^-$ can diffuse across the thylakoid membranes, (2) the thylakoid lumen maintains a low pH, and (3) there is thylakoid CA in the vicinity of the pyrenoid matrix to convert $HCO_3^-$ to $CO_2$. To test this hypothesis, we study models with distinct thylakoid morphologies as described below. As a proof of concept, we only consider the passive $CO_2$ uptake strategy under air-level $CO_2$, i.e., $C_{\mathrm{cyt}} = 10$ μM, and we ignore the spontaneous conversion between $HCO_3^-$ and $CO_2$ for simplicity. Parameters are the same as in the baseline Chlamydomonas CCM model.

We start by considering a model with a shell of thylakoid sheet surrounding the pyrenoid matrix (see Fig. S26, model 1). Indeed, this thylakoid morphology has been found in other algae [67]. The equations describing the steady-state flux balance of $CO_2$, $HCO_3^-$, and $H_2CO_3$ are:

$$\begin{cases} r^{-2}\partial_r(r^2 D^C \partial_r C_{\mathrm{pyr}}) - j_{\mathrm{Rbc}} = 0 \\ r^{-2}\partial_r(r^2 D^H \partial_r H_{\mathrm{pyr}}) = 0 \end{cases} \qquad \text{for } r \leqslant R_{\mathrm{pyr}}, \tag{S74a}$$

$$\begin{cases} r^{-2}\partial_r(r^2 D^C \partial_r C_{\mathrm{thy}}) - j_{\mathrm{CAH3}} = 0 \\ r^{-2}\partial_r(r^2 D^H \partial_r H_{\mathrm{thy}}) + j_{\mathrm{CAH3}} = 0 = 0 \end{cases} \qquad \text{for } R_{\mathrm{pyr}} \leqslant r \leqslant R_{\mathrm{pyr}} + \Delta R_{\mathrm{thy}}, \tag{S74b}$$

$$\begin{cases} r^{-2}\partial_r(r^2 D^C \partial_r C_{\mathrm{str}}) - j_{\mathrm{LCIB}} = 0 \\ r^{-2}\partial_r(r^2 D^H \partial_r H_{\mathrm{str}}) + j_{\mathrm{LCIB}} = 0 = 0 \end{cases} \qquad \text{for } R_{\mathrm{pyr}} + \Delta R_{\mathrm{thy}} \leqslant r \leqslant R_{\mathrm{chlor}}. \tag{S74c}$$

Here, we assume that the CCM enzymes are distributed diffusely in their respective compartments and that their reaction fluxes $j_{\mathrm{X}}$ take the same form as in the Chlamydomonas CCM model (Eqs. (S10, S11) in Sec. I B). We set the radial width of the thylakoid sheet to be $\Delta R_{\mathrm{thy}} = 0.05\ R_{\mathrm{chlor}}$. The boundary conditions of Eq. (S74) are determined by the inter-compartmental transport fluxes of Ci species. Specifically, we obtain

$$\begin{cases} D^C\partial_r C_{\rm pyr} = D^C\partial_r C_{\rm thy} = \kappa^C(C_{\rm thy} - C_{\rm pyr}) \\ D^H\partial_r H_{\rm pyr} = D^H\partial_r H_{\rm thy} = \kappa^{H^0}(H^0_{\rm thy} - H^0_{\rm pyr}) + (\kappa^{H^-} + \kappa^{H^-}_{\rm thy})(H^-_{\rm thy} - H^-_{\rm pyr}) \end{cases} \quad \text{at } r = R_{\rm pyr}, \tag{S75a}$$

$$\begin{cases} D^C\partial_r C_{\rm thy} = D^C\partial_r C_{\rm str} = \kappa^C(C_{\rm str} - C_{\rm thy}) \\ D^H\partial_r H_{\rm thy} = D^H\partial_r H_{\rm str} = \kappa^{H^0}(H^0_{\rm str} - H^0_{\rm thy}) + (\kappa^{H^-} + \kappa^{H^-}_{\rm thy})(H^-_{\rm str} - H^-_{\rm thy}) \end{cases} \quad \text{at } r = R_{\rm pyr} + \Delta R_{\rm thy}, \tag{S75b}$$

$$\begin{cases} D^C\partial_r C_{\rm str} = \kappa^C(C_{\rm cyt} - C_{\rm str}) \\ D^H\partial_r H_{\rm str} = \kappa^{H^0}(H^0_{\rm cyt} - H^0_{\rm str}) + (\kappa^{H^-} + \kappa^{H^-}_{\rm chlor})(H^-_{\rm cyt} - H^-_{\rm str}) \end{cases} \quad \text{at } r = R_{\rm chlor}, \tag{S75c}$$

where $\kappa^{H^-}_{\rm thy}$ and $\kappa^{H^-}_{\rm chlor}$ denote, respectively, the additional permeability to $HCO_3^-$ due to BST channels on the thylakoid membranes and due to LCIA channels on the chloroplast envelope. We again assume that all Ci species in the cytosol are in equilibrium at pH = 7.1. As shown in Fig. S26 (model 1), this modeled thylakoid morphology could also achieve half-saturation of Rubisco, although at a higher energetic cost. In this model, the two layers of thylakoid membranes serve as diffusion barriers to prevent $CO_2$ leakage out of the pyrenoid matrix, but they are less effective than the impermeable starch sheath in the Chlamydomonas CCM model. Indeed, if we add an impermeable starch sheath with the same gaps between starch plates as the one considered in the Chlamydomonas CCM model to further surround the shell of thylakoid sheet, i.e., multiply $\kappa$ by $f_{\rm v}$ (Table S1) in Eq. S75b, we could achieve a more efficient CCM (Fig. S26, model 2). This geometry is energetically even more efficient than the native Chlamydomonas CCM, because the thylakoid membranes spanning the holes on the starch sheath could slow the escape of $CO_2$ produced in the thylakoid lumen. Note that in these models, we have assumed that low pH is always maintained in the pyrenoid-proximal thylakoids, which is presumably achieved by connecting to the surrounding thylakoids in the stroma as a source of protons.

To account this feature, we finally consider a Chlamydomonas-like configuration with a starch sheath (Fig. 2G), but with the thylakoids inside the pyrenoid simple forming a shell of width $\Delta R_{\rm thy}$ (Fig. S26, model 3). The steady-state flux-balance equations are the same as Eqs. (S74) and (S75), except that

$$\text{Eq. (S74c)} \Rightarrow \begin{cases} D^C \frac{1}{f_{\rm v} r^2}\frac{\partial}{\partial r}\left(f_{\rm v} r^2 \frac{\partial C_{\rm tub}}{\partial r}\right) - j^C_{\rm mem} f_{\rm s} = 0 \\ D^C \frac{1}{(1-f_{\rm v}) r^2}\frac{\partial}{\partial r}\left((1-f_{\rm v}) r^2 \frac{\partial C_{\rm str}}{\partial r}\right) - j_{\rm LCIB} + j^C_{\rm mem}\frac{f_{\rm s} f_{\rm v}}{1-f_{\rm v}} = 0 \\ D^H \frac{1}{f_{\rm v} r^2}\frac{\partial}{\partial r}\left(f_{\rm v} r^2 \frac{\partial H_{\rm tub}}{\partial r}\right) - j^H_{\rm mem} f_{\rm s} = 0 \\ D^H \frac{1}{(1-f_{\rm v}) r^2}\frac{\partial}{\partial r}\left((1-f_{\rm v}) r^2 \frac{\partial H_{\rm str}}{\partial r}\right) + j_{\rm LCIB} + j^H_{\rm mem}\frac{f_{\rm s} f_{\rm v}}{1-f_{\rm v}} = 0 \end{cases} \quad \text{for } R_{\rm pyr} + \Delta R_{\rm thy} \leqslant r \leqslant R_{\rm chlor} \tag{S76}$$

$$\text{Eq. (S75b)} \Rightarrow \begin{cases} \partial_r C_{\rm thy} = \partial_r C_{\rm tub}, C_{\rm thy} = C_{\rm tub}, \partial_r C_{\rm str} = 0 \\ \partial_r H_{\rm thy} = \partial_r H_{\rm tub}, H_{\rm thy} = H_{\rm tub}, \partial_r H_{\rm str} = 0 \end{cases} \quad \text{at } r = R_{\rm pyr} + \Delta R_{\rm thy}, \tag{S77}$$

describing the Ci concentrations in the stroma and in the stromal thylakoid tubules (denoted by subscript "tub"). Here, the cross-membrane fluxes $j_{\rm mem}$ are the same as in Eqs. (S13) and (S14). Indeed, the CCM model using this hypothetical thylakoid morphology yields an efficacy-cost relation that closely resembles the Chlamydomonas model (Fig. S26J). Thus, we conclude that the functioning of the CCM is largely independent of the morphologies of intra-pyrenoid thylakoids, but that thylakoid morphologies could affect $CO_2$ leakage out of the pyrenoid, which determines the energetic efficiency of the CCM.

##### D. Effect of LCIB-Rubisco complexes

Modeling the transfer of algal machinery into a plant cell requires an approximation of the starting chloroplast configuration in plants (Fig. 6). In our model, we have assumed that plant chloroplasts lack any CCM activity. Here, we offer a justification for this assumption. Previous experiments have suggested that carbonic anhydrases (CAs) and Rubisco could form protein complexes in certain plant chloroplasts [68, 69]. Based on these observations, it has been suggested that protons produced during $CO_2$ fixation could locally decrease the pH, favoring the production of $CO_2$. A recent work [64] suggests that in a modeled compartment with a diffusion barrier, Rubisco-derived protons could indeed drive the conversion of $HCO_3^-$ to $CO_2$ via co-condensed CA, thus increasing local $CO_2$ concentration and the rate of Rubisco $CO_2$ fixation. We wondered whether a similar mechanism could work for protein complexes of plant Rubisco with a stromal CA, hence rendering noticeable $CO_2$-concentrating activity to the plant chloroplasts.

Consider a condensate of Rubisco and LCIB whose radius is denoted by $R_{\rm cond}$. We explicitly consider the reactions involving $CO_2$, $HCO_3^-$, and $H^+$ inside the condensate, as well as their diffusion in and out of the condensate (Fig. S27A). Denote by $P$, $C$, and $H^-$ the concentrations of protons, $CO_2$, and $HCO_3^-$, respectively. At steady state, the flux balance of $HCO_3^-$ in the condensate requires that the total flux of $HCO_3^-$-to-$CO_2$ conversion by LCIB should be equal to the influx of $HCO_3^-$, i.e.,

$$\frac{4}{3}\pi R_{\rm cond}^3 k_{\rm LCIB}\Big(\frac{P}{P_{\rm eq}}H^- - C\Big) = 4\pi R_{\rm cond} D^H (H^-_\infty - H^-), \tag{S78}$$

where $k_{\rm LCIB}$ denotes the first-order rate constant of LCIB, $P_{\rm eq} = 10^{-{\rm pK_{eff}}} = 10^{-6.1}$ M denotes the proton concentration at which $CO_2$ and $HCO_3^-$ have equal equilibrium concentrations, and $H^-_\infty$ denotes the concentration of $HCO_3^-$ in the bulk stroma away from the condensate. Here, we have used a linear approximation for CA kinetics, and we have again used the Burg-Purcell calculation for the $HCO_3^-$ influx. At steady state, $CO_2$ converted by LCIB must either be fixed by Rubisco or diffuse out of the condensate, i.e.,

$$\frac{4}{3}\pi R_{\rm cond}^3 k_{\rm LCIB}\Big(\frac{P}{P_{\rm eq}}H^- - C\Big) = \frac{4}{3}\pi R_{\rm cond}^3 V^C_{\rm max,Rbc}\frac{C}{C + K^{\rm eff}_{\rm m}} + 4\pi R_{\rm cond} D^C (C - C_\infty), \tag{S79}$$

where $C_\infty$ denotes the concentration of $CO_2$ in the bulk stroma. Finally, the protons involved in the $HCO_3^-$-to-$CO_2$ conversion must either come from Rubisco $CO_2$ fixation or from the bulk stroma, which yields

$$\frac{4}{3}\pi R_{\rm cond}^3 k_{\rm LCIB}\Big(\frac{P}{P_{\rm eq}}H^- - C\Big) = 2\times\Big(\frac{4}{3}\pi R_{\rm cond}^3 V^C_{\rm max,Rbc}\frac{C}{C + K^{\rm eff}_{\rm m}}\Big) + 4\pi R_{\rm cond} D_P (P_\infty - P). \tag{S80}$$

Here, the factor 2 accounts for the two protons produced for each $CO_2$ fixed.

**Choice of parameters.** Consider a stroma of pH 8 under air-level $CO_2$ (10 μM cytosolic), i.e., $P_\infty = 10^{-8}$ M $= 10^{-5}$ mM, and $C_\infty = 10$ μM $= 0.01$ mM. We further assume that $HCO_3^-$ is equilibrated with $CO_2$, i.e., $H^-_\infty = 10^{2.9}$ μM $\approx 0.794$ mM (however, this assumption is not crucial for the main conclusion we will draw below). We assume that the Rubisco density is the same as that in the phase separated droplet in Chlamydomonas, and thus $V^C_{\rm max,Rbc} = 15$ mM/$s$. $K^{\rm eff}_{\rm m}$ of Rubisco is 0.076 mM (Table S1). We choose $k_{\rm LCIB} = 10^6$ s$^{-1}$ since the typical turnover number for CA is $10^6$ s$^{-1}$ and in a condensate the concentration of protein could be in the mM range, similar to the $K_{\rm m}$ of typical CA. The values of the diffusion coefficients are the same as previous sections (see Sec. I A, Sec. VI A and Table S1).

Solving Eqs. (S78 – S80) at varying $R_{\rm cond}$, we find that for small $R_{\rm cond}$, the reaction fluxes of enzymes in the condensate are negligible since they scale with $R^3_{\rm cond}$ while the absorption/leakage fluxes scale with $R_{\rm cond}$, and thus the concentration of $CO_2$ in the condensate is roughly the same as the environment (Fig. S27B). The mechanism discussed above doubles the $CO_2$ concentration in the condensate roughly when the two terms on the right-hand side of Eq. (S79) are comparable, i.e., when $R_{\rm cond} \gtrsim (3D^C C_\infty/V^C_{\rm max,Rbc})^{1/2} \approx 2$ μm, which is much larger than the size of protein complexes we consider. In the limit of large $R_{\rm cond}$, the absorption of carbon molecules become diffusion-limited, which leads to a low $CO_2$ concentration in the condensate again (Fig. S27C).

##### E. Import of bicarbonate into the cell

In our model, we have assumed constant cytosolic $CO_2$ and $HCO3^-$ concentrations representing external Ci conditions. Here, we discuss how cytosolic concentrations of inorganic carbon may be maintained at steady state. Since there is no known CA in the Chlamydomonas cytosol, cytosolic $CO_2$ and $HCO_3^-$ presumably need to be replenished separately, either by diffusion or transport. We expect that $CO_2$ can diffuse across the cell envelope, supporting the $CO_2$ assimilation flux by the chloroplast. A similar calculation to that in Sec. III D can be made to estimate the conditions under which the passive $CO_2$ uptake strategy is feasible. Below, we focus on the transport of $HCO_3^-$ into the Chlamydomonas cell. In particular, we estimate under limiting conditions of 10 μM $HCO_3^-$ outside the cell (corresponding to equilibrium with air-level $CO_2$ at pH 6), (1) how many $HCO_3^-$ channels are required on the cell membrane to support the flux used by the active $HCO_3^-$ pumping strategy, and (2) the cost of synthesizing such

$HCO_3^-$ channels.

To start with, we can consider the extreme case where the rate of $HCO_3^-$ transport into the cytosol is not limited by the turnover rate of the hypothetical $HCO_3^-$ channels on the cell envelope, but is rather limited by diffusion through the cavity of these channels. In this case, the diffusive flux through a single channel can be estimated by $J_{\text{hole}} = 2D^H r_{\text{hole}} H^-_{\text{env}}/(1+\frac{2h}{\pi r_{\text{hole}}})$ [56], where $D^H \approx 10^3$ μm$^2$/s denotes the diffusion constant of $HCO_3^-$, $r_{\text{hole}} \approx 1$ nm denotes the radius of the cavity, $h \approx 5$ nm denotes the length of the channel and $H^-_{\text{env}} = 10$ μM denotes the bulk concentration of $HCO_3^-$ in the environment. Plugging in these numbers yields a flux of $J_{\text{hole}} = 0.5 \times 10^{-20}$ mol/s. Since this estimation gives the upper limit, we choose to use a value of $J_{\text{hole}} = 0.5 \times 10^{-21}$ mol/s, i.e., an order of magnitude smaller than the upper bound, as a more realistic estimation of the single-channel flux. Our simulation results show that an active $HCO_3^-$ pumping flux of $2.4 \times 10^{-16}$mol/s across the chloroplast envelope can drive $> 70\%$ of the maximum $CO_2$ fixation flux (Fig. S12). To support this flux, a total of $4.8 \times 10^5$ channels are needed. Finally, one can estimate the fraction of the cell membrane surface area occupied by this many channels. Assuming that the outer radius of the channel is 3.5 nm (a typical value) and that the radius of a Chlamydomonas cell is 5 μm, one can obtain the surface area fraction taken by the channels to be $4.8 \times 10^5 \times (3.5\ \text{nm})^2/(5\ \mu\text{m})^2/4 \approx 6\%$.

To estimate the energetic cost of synthesizing the above number of channels, we consider the synthesis and polymerization of 1000 amino acids per channel, similar to the size of the putative $HCO_3^-$ transporters on the Chlamydomonas cell membrane. Assuming that the cost of protein synthesis is 30 ATP/aa, we obtain the cost to synthesize one $HCO_3^-$ transporter to be 30,000 ATP. The doubling time of Chlamydomonas cells across a wide range of conditions has been measured to be 10 hours [70]. Over this period of time, a Chlamydomonas cell presumably needs to synthesize $4.8 \times 10^5$ transporters as discussed above, yielding an energy consumption rate of (30,000 ATP/transporter) $\times$ ($4.8 \times 10^5$ transporters) / (10 hours) = $4 \times 10^5$ ATP/s. To compare this cost with the cost of operating the CCM, we employ the simulation results that the saturating $CO_2$ fixation flux is $6 \times 10^{-17}$ mol/s and the cost is roughly 1 ATP per $CO_2$ fixed (see main Fig. 3). Thus, an effective CCM will cost $\sim 3.6 \times 10^7$ ATP/s, which is roughly two orders of magnitude larger than the cost of synthesizing the $HCO_3^-$ transporters.

#### SUPPLEMENTARY FIGURES AND TABLES

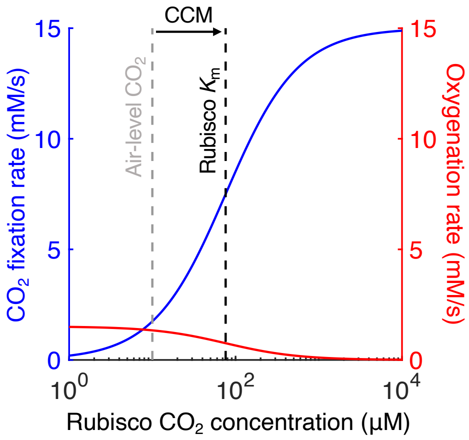


**Fig. S1.** $CO_2$ fixation by Rubisco is inhibited by oxygen at air-level $CO_2$. Rates of $CO_2$ fixation (blue, Eq. S11) and oxygenation (red, Eq. S27) by Rubisco at varying $CO_2$ concentrations. Gray dashed line denotes air-level $CO_2$ (10 µM cytosolic). Black dashed line denotes the effective $K_m$ of Rubisco (76 µM, see Sec. I B). Kinetic parameters of Rubisco are listed in Table S1.

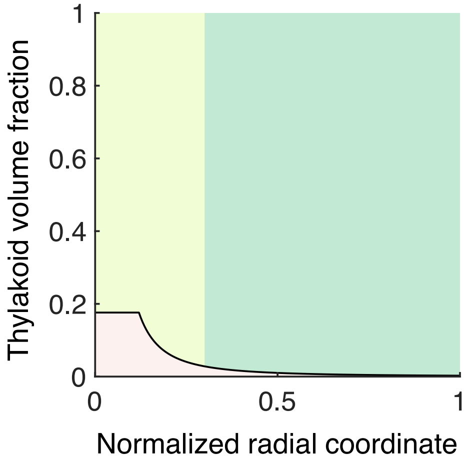


**Fig. S2.** Volume fraction of the modeled chloroplast compartments. Volume fraction of the chloroplast taken up by the thylakoid tubules (pink), pyrenoid matrix (light green), and stroma (green), versus the normalized radial coordinate. Note that our model does not consider the volume of thylakoid stacks traversing the stroma.

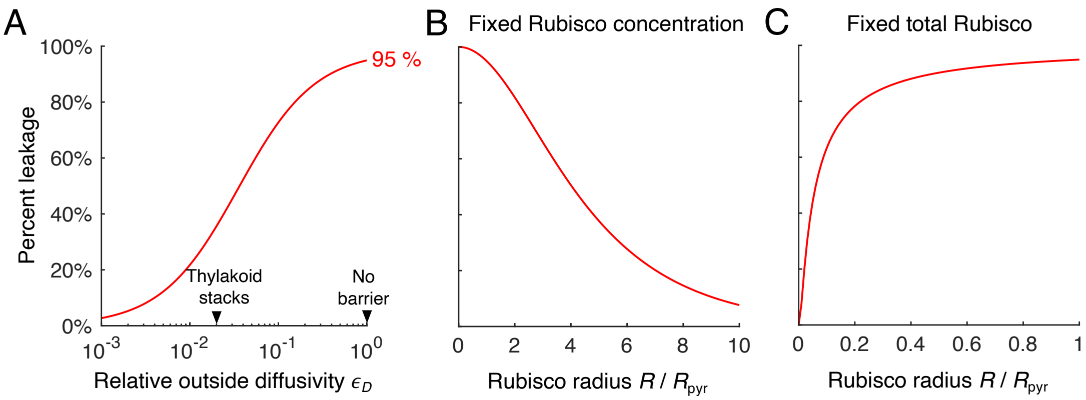


**Fig. S3.** Low fixation efficiency at air-level $CO_2$ (10 µM cytosolic) in the absence of a diffusion barrier is due to severe $CO_2$ leakage from the pyrenoid matrix. $CO_2$ percent leakage computed from a simplified reaction-diffusion model (see Sec. I H for details) for (*A*) varying relative diffusivity $\epsilon_D$ outside the sphere of Rubisco, and at varying hypothetical Rubisco radii assuming (*B*) fixed Rubisco concentration or (*C*) fixed number of Rubisco molecules. In *A*, Rubisco radius $R = R_{\text{pyr}}$, and arrowheads mark the $\epsilon_D$ values corresponding to the case of diffusion slowed by thylakoid stacks and the case of no diffusion barrier. In *B* and *C*, relative diffusivity outside the Rubisco region compared to inside the Rubisco region is $\epsilon_D = 1$.

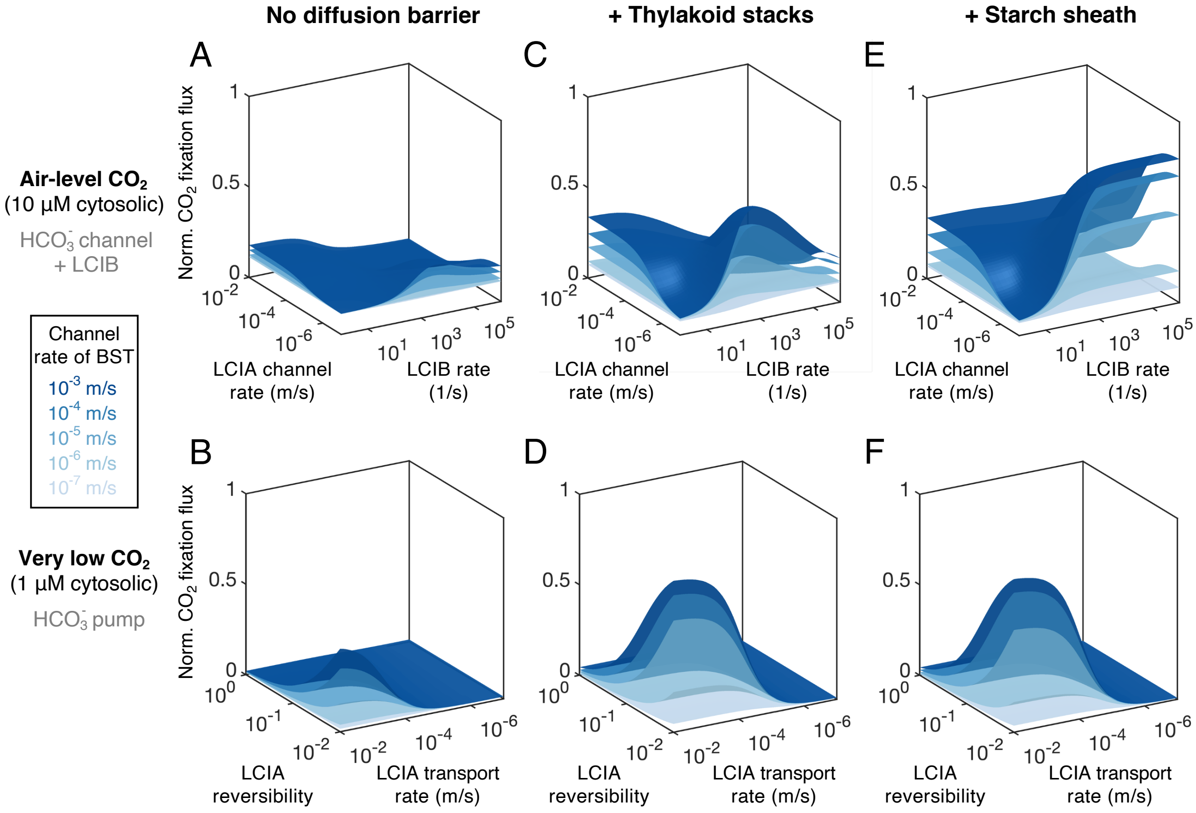


**Fig. S4.** $CO_2$ fixation flux increases with BST $HCO_3^-$ channel rate. This is verified in (*A*, *B*) a model without a diffusion barrier, (*C*, *D*) a model with thylakoid stacks slowing inorganic carbon diffusion in the stroma, and (*E*, *F*) a model with an impermeable starch sheath. (*A*, *C*, *E*) A modeled chloroplast employing LCIB and $HCO_3^-$ channel transport across the chloroplast envelope is considered under air-level $CO_2$ (10 μM cytosolic). Normalized $CO_2$ fixation fluxes at varying LCIA channel rates and LCIB rates are shown for the designated BST channel rates. (*B*, *D*, *F*) A modeled chloroplast employing active $HCO_3^-$ pumping across the chloroplast envelope and no LCIB is considered under very low $CO_2$ (1 μM cytosolic). Normalized $CO_2$ fixation fluxes at varying LCIA transport rates and reversibilities are shown for the designated BST channel rates.

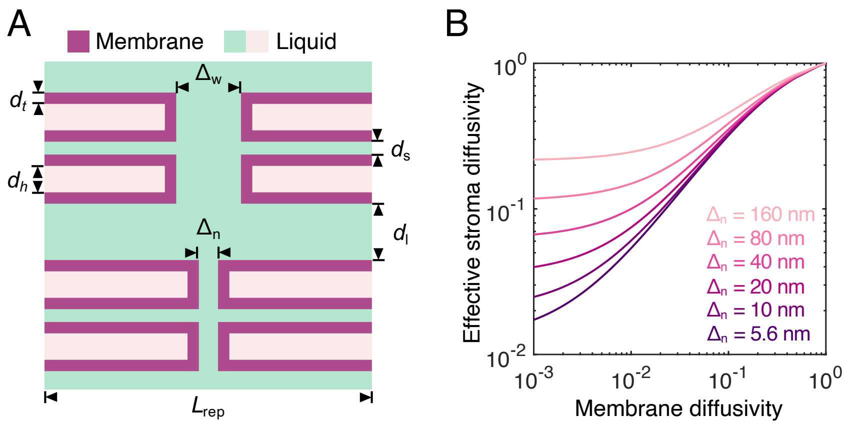


**Fig. S5.** Model for the effect of thylakoid stacks on diffusion in the stroma. (*A*) Schematic geometry of modeled thylakoid stacks. Key geometric parameters are described in Sec. I G. Color code is the same as in Fig. 1A. (*B*) Effective stroma diffusivity plotted against membrane diffusivity for the designated gap sizes $\Delta_\mathrm{n}$ of the thylakoid stacks. Diffusivities are normalized by the diffusion coefficient in liquid (see Sec. I G for details). $\Delta_\mathrm{w} - \Delta_\mathrm{n}$ is fixed when varying $\Delta_\mathrm{n}$ in the simulations. We infer from previous experiments that $\Delta_\mathrm{n} = 5.6$ nm [1], and relative membrane diffusivity is roughly $10^{-3}$ [29]. Values of other geometric parameters are provided in Sec. I G **Choice of parameters**.

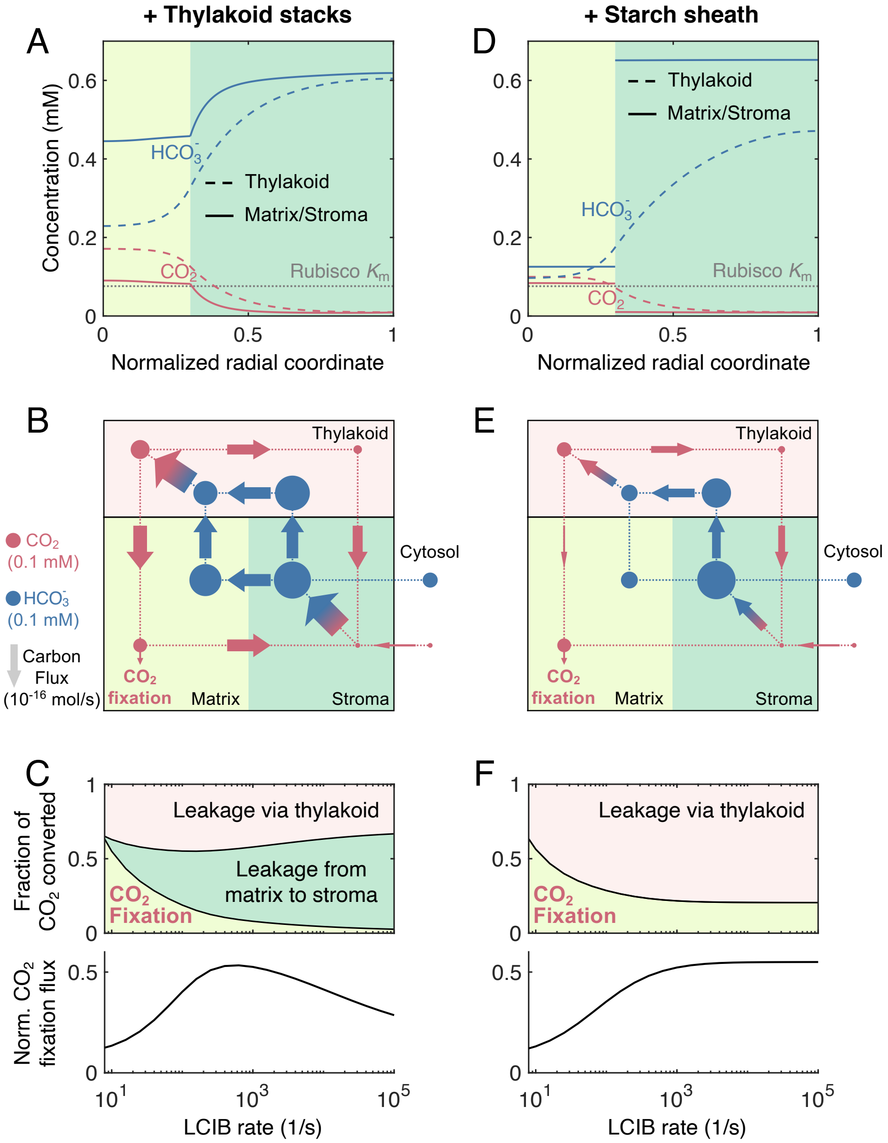


**Fig. S6.** Comparison between two types of diffusion barriers under air-level $CO_2$ (10 μM cytosolic). A model with thylakoid stacks slowing inorganic carbon diffusion in the stroma (*A*–*C*) is compared to a model with an impermeable starch sheath (*D*–*F*). (*A*, *D*) Concentration profiles of $CO_2$ and $HCO_3^-$ in the thylakoid (dashed curves) and in the matrix/stroma (solid curves). Dotted gray line indicates the effective Rubisco $K_m$ for $CO_2$ (see Sec. I B). Plots correspond to Fig. 1C. (*B*, *E*) Net fluxes of inorganic carbon between the indicated CCM compartments. Plots correspond to Fig. 1D. (*C*, *F*) Decomposition of the total $CO_2$ flux converted by CAH3 (top) and normalized $CO_2$ fixation flux (bottom, see Sec. II A for definition) at varying rates of LCIB activity. Both models have minimal $HCO_3^-$ transport across the chloroplast envelope. $HCO_3^-$ channel transport rates across the thylakoid membrane are $\kappa_{\rm thy}^{H^-} = 10^{-4}$ m/s in *A*–*C* and $\kappa_{\rm thy}^{H^-} = 10^{-4.8}$ m/s in *D*–*F*, which are chosen such that the maximum normalized $CO_2$ fixation flux is roughly 0.5. LCIB rates are $V_{\rm max,LCIB}^{C}/K_{\rm m}^{C} = 10^3$ s$^{-1}$ in *A*–*B* and *D*–*E*.

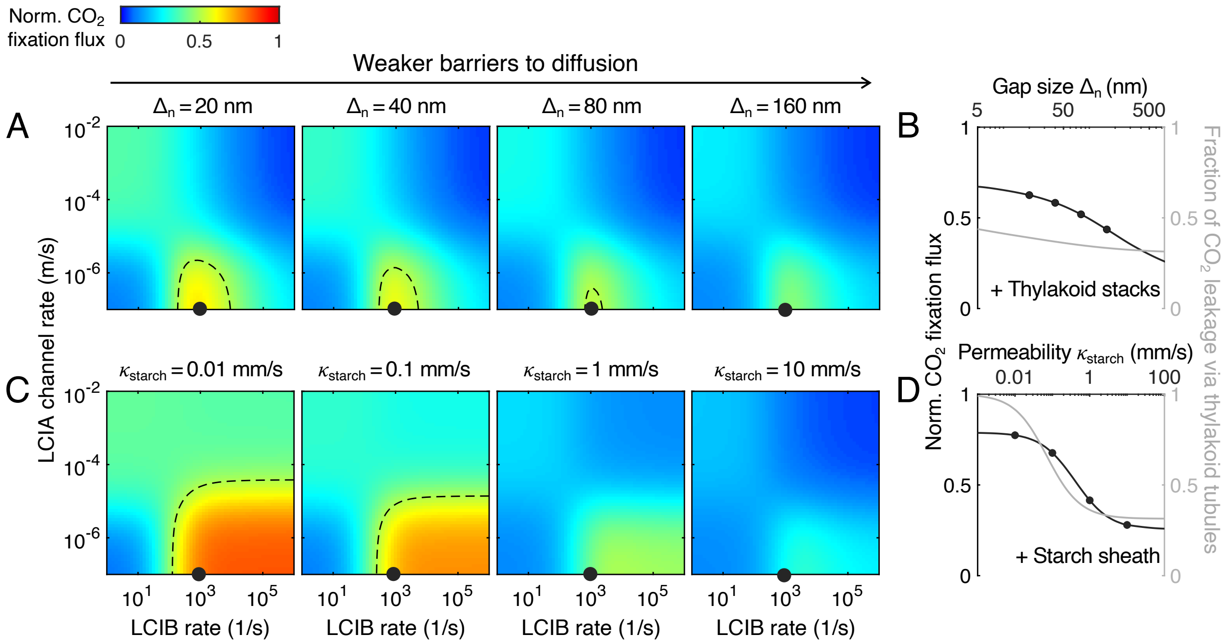


**Fig. S7.** $CO_2$-concentrating performance is compromised by weaker barriers to the diffusion of inorganic carbon. A model with thylakoid stacks slowing inorganic carbon diffusion in the stroma (*A*, *B*) is compared to a model with a starch sheath (*C*, *D*) under air-level $CO_2$ (10 μM cytosolic). (*A*, *C*) Heatmaps of normalized $CO_2$ fixation flux (see Sec. II A) at varying LCIA $HCO_3^-$ channel transport rates and LCIB activities shown for the designated parameters of the two diffusion barriers. Plots correspond to Figs. 2*E* and *H*. (*B*, *D*) Normalized $CO_2$ fixation flux (black) and fraction of $CO_2$ leakage from the pyrenoid that occurs via thylakoid tubules (and not directly from the pyrenoid matrix into the stroma) (gray) at varying strengths of the diffusion barriers. The rates of LCIA $HCO_3^-$ channel transport and LCIB activity are indicated by black dots in *A* and *C*.

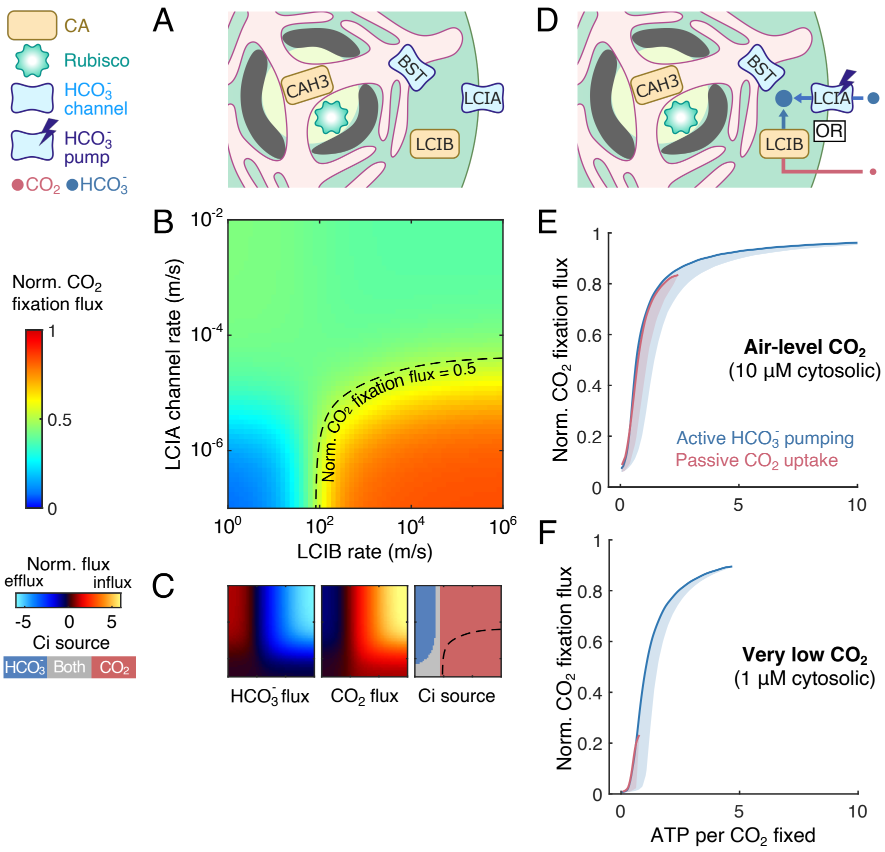


**Fig. S8.** A model including both thylakoid stacks and an impermeable starch sheath behaves similarly to a model with only the latter. (*A*) Schematic of a modeled chloroplast with both thylakoid stacks and impermeable starch sheath, employing LCIB and $HCO_3^-$ channel transport across the chloroplast envelope. (*B*) Heatmap of normalized $CO_2$ fixation flux (see Sec. II A) at varying LCIA $HCO_3^-$ channel transport rates and LCIB activities. (*C*) Heatmaps of (*left*) the total $HCO_3^-$ flux, (*middle*) the total $CO_2$ flux, and (*right*) the type(s) of inorganic carbon (Ci), taken up by the chloroplast. Fluxes are normalized by the maximum $CO_2$ fixation flux for saturated Rubisco. Negative values of flux indicate efflux. The $x$ and $y$ axes are the same as in *B*. Panels *A*-*C* correspond to Fig. 2. In *B* and *C*, parameters are the same as in Fig. 2. (*D*) Schematic of a modeled chloroplast with both thylakoid stacks and impermeable starch sheath, employing active $HCO_3^-$ pumping across the chloroplast envelope and no LCIB activity (blue), or employing LCIB for passive $CO_2$ uptake (red). (*E, F*) CCM performance under (*E*) air-level $CO_2$ (10 μM cytosolic) and (*F*) very low $CO_2$ (1 μM cytosolic), measured by normalized $CO_2$ fixation flux versus ATP spent per $CO_2$ fixed (see Sec. II B), for the two CCM strategies in *A*. Solid curves indicate the minimum energy input necessary to achieve a certain normalized $CO_2$ fixation flux. Shaded regions represent the range of possible performances found by varying $HCO_3^-$ transport rates and/or LCIB rates. Panels *D*-*F* correspond to Fig. 3.

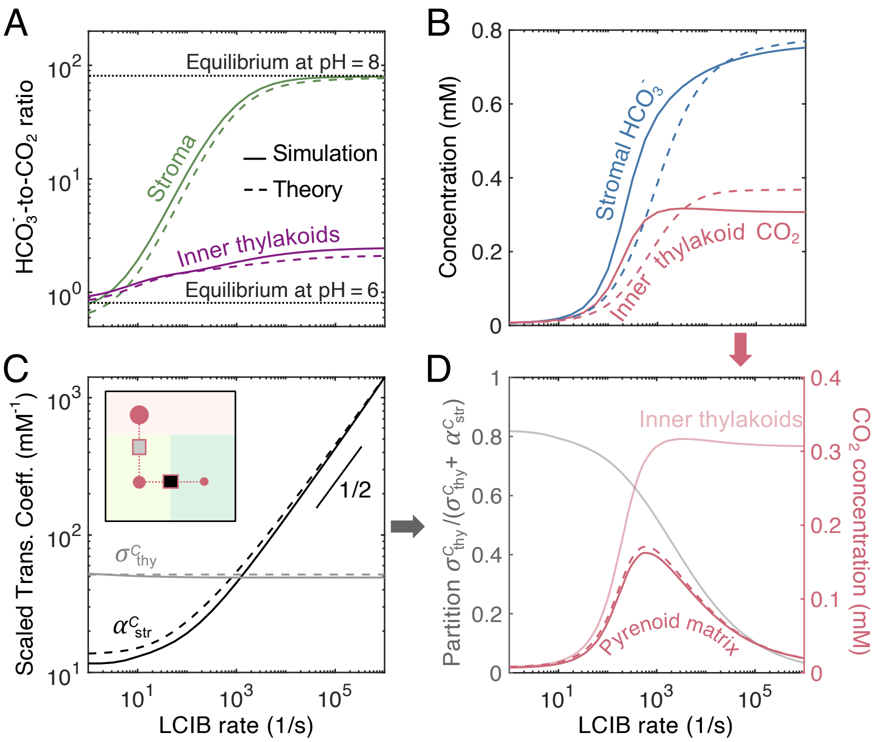


**Fig. S9.** In a modeled chloroplast employing thylakoid stacks and no starch sheath, LCIB activity increases intra-chloroplast $HCO_3^-$ concentration by converting external $CO_2$ but also effectively "steals" $CO_2$ from the pyrenoid matrix, leading to a nonmonotonic impact of LCIB rate on $CO_2$ concentration inside the pyrenoid matrix. (*A*) The concentration ratio of $HCO_3^-$ to $CO_2$ in the stroma (green) and in the thylakoid tubules inside the pyrenoid (purple; denoted by inner thylakoids) at varying LCIB rates. Black dotted lines indicate the equilibrium $HCO_3^-$-to-$CO_2$ ratio for the designated pH values. (*B*) Concentrations of $HCO_3^-$ in the stroma (blue) and $CO_2$ in the inner thylakoids increase with LCIB rates. (*C*) The linear transport coefficients of $CO_2$, defined in Sec. III C, between the thylakoid tubules and pyrenoid matrix (gray) and between the matrix and stroma (black) at varying LCIB rates. *Inset:* schematic of the transport coefficients. Color code is the same as Fig. 1D. The short black line indicates a slope of 1/2 on a log-log scale. (*D*) Decreasing partition factor $\sigma^C_{\rm thy}/(\alpha^C_{\rm str} + \sigma^C_{\rm thy})$ defined in Eq. S67 (light gray) and increasing concentration of $CO_2$ in the inner thylakoids (light red) with increasing LCIB rates together lead to the nonmonotonic dependence of pyrenoid matrix $CO_2$ level on the rate of LCIB. Solid and dashed curves denote results from the full simulation and the simplified model Eq. S66, respectively.

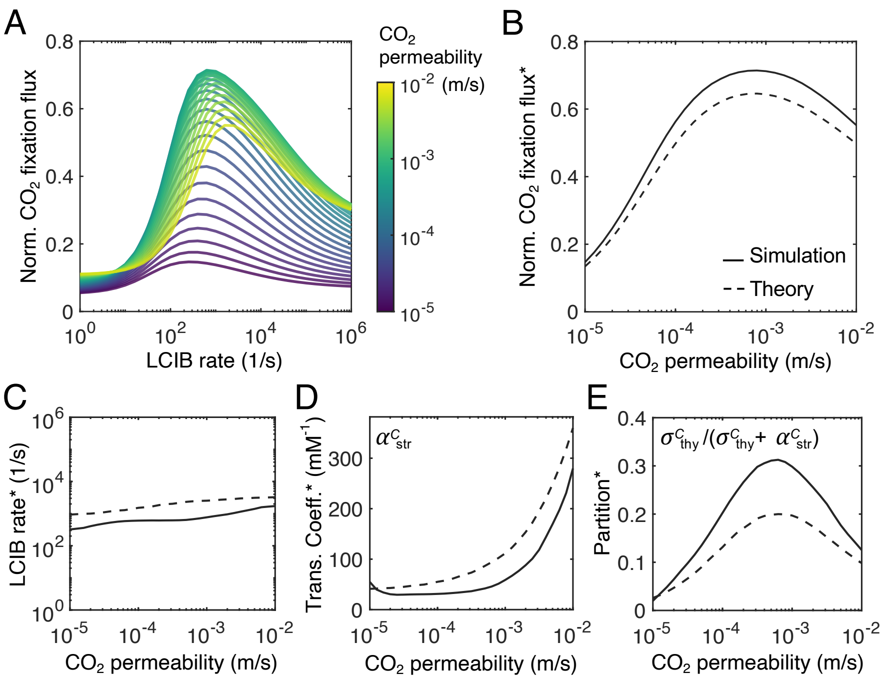


**Fig. S10.** A trade-off between $CO_2$ transport and $CO_2$ leakage leads to the existence of an intermediate membrane permeability to $CO_2$ that optimizes $CO_2$-concentrating performance. (*A*) Normalized $CO_2$ fixation flux is plotted against LCIB rate for varying membrane permeabilities to $CO_2$ ($\kappa^C$) in a modeled chloroplast with thylakoid stacks slowing the diffusion of inorganic carbon in the stroma and no starch sheath. (*B*, *C*) The maximum normalized $CO_2$ fixation flux (shown in *B*) and the correponding LCIB rate (shown in *C*), denoted by superscript *, at varying membrane permeabilities to $CO_2$. (*D*, *E*) The transport coefficient $\alpha^C_{\rm str}$ between the pyrenoid matrix and stroma (shown in *D*; see Sec. IV B) and the partition factor $\sigma^C_{\rm thy}/(\alpha^C_{\rm str} + \sigma^C_{\rm thy})$ defined in Eq. S67 (shown in *E*) at varying membrane permeabilities to $CO_2$ when normalized $CO_2$ fixation flux is maximized with respect to varying LCIB rates. In *B*–*E*, solid and dashed curves denote results from the full simulation and the simplified model Eq. S66, respectively.

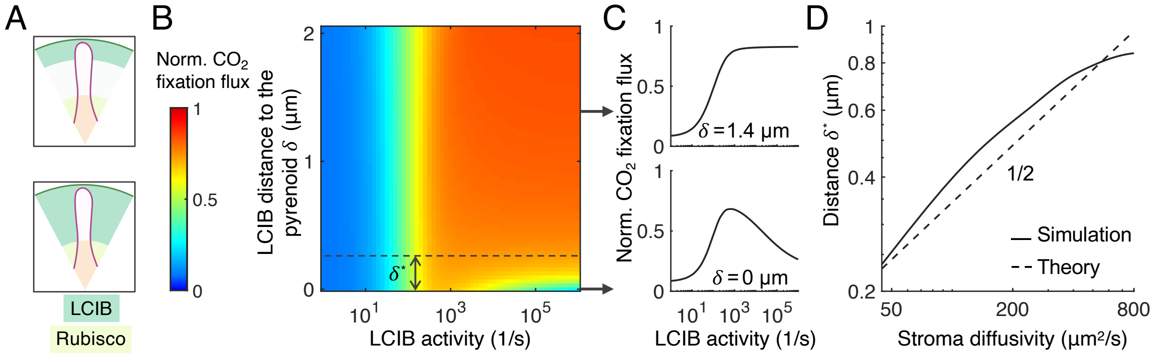


**Fig. S11.** Localizing LCIB close to the chloroplast envelope prevents excessive $CO_2$ efflux drawn by LCIB from the pyrenoid matrix in a modeled chloroplast with thylakoid stacks slowing diffusion in the stroma and no starch sheath. (*A*) Schematics of hypothetical LCIB localization patterns. (*B*) Normalized $CO_2$ fixation flux at varying LCIB activities and LCIB localization patterns. LCIB activity is characterized by the first-order rate constant when LCIB is uniformly distributed in the stroma. LCIB localization is characterized by its distance $\delta$ to the pyrenoid. For a given LCIB activity, the total number of LCIB is held constant when varying the localization pattern. (*C*) Normalized $CO_2$ fixation flux at varying LCIB activities for the designated values of $\delta$, indicated by the arrows in *B*. (*D*) The critical distance $\delta^*$ between the pyrenoid and LCIB, defined as the distance above which the normalized $CO_2$ fixation flux increases monotonically with LCIB activity (indicated in *B*), at varying $CO_2$ stroma diffusivities is plotted for the simulation result (solid) and the theoretical prediction (dashed; see Sec. IV C **Localization of LCIB**). 1/2 indicates the slope of the theory line on a log-log scale. For all panels, simulation parameters are the same as Figs. 1C and D.

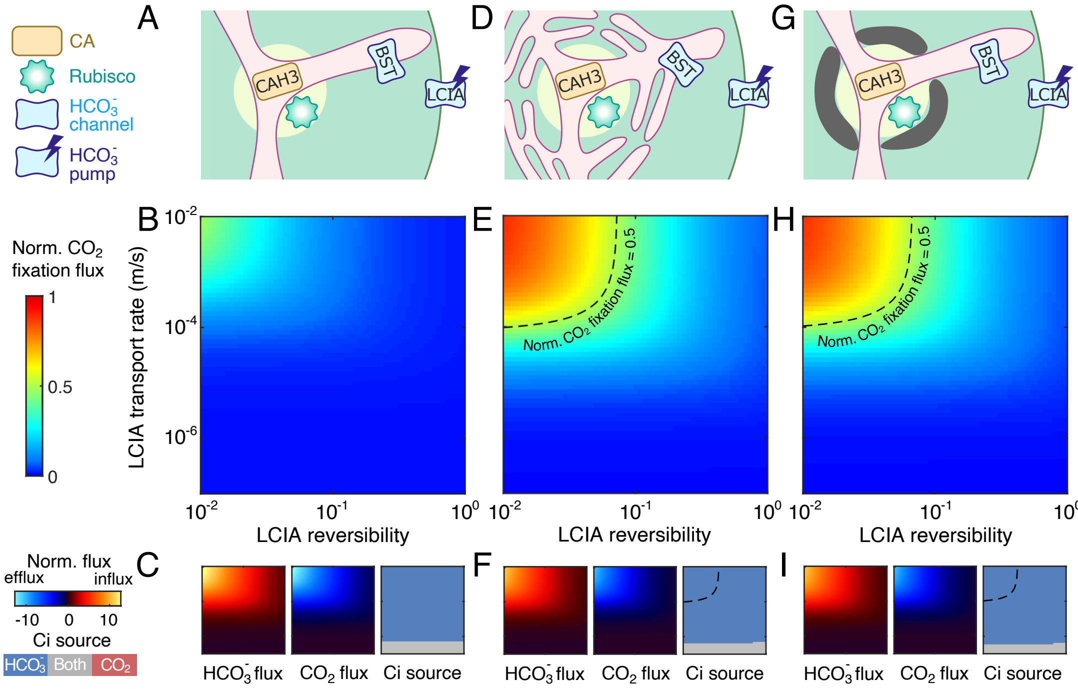


**Fig. S12.** Active $HCO_3^-$ transport across the chloroplast envelope enables an effective CCM under very low $CO_2$. A model with no barrier to $CO_2$ diffusion out of the pyrenoid matrix (*A*–*C*) is compared to a model with thylakoid stacks slowing inorganic carbon diffusion in the stroma (*D*–*F*) and a model with an impermeable starch sheath (*G*–*I*) under very low $CO_2$ (1 µM cytosolic). (*A*, *D*, and *G*) Schematics of the modeled chloroplast. (*B*, *E*, and *H*) Heatmaps of normalized $CO_2$ fixation flux at varying LCIA rates and reversibilities. The BST channel transport rate across the thylakoid membrane is the same as in Figs. 1C and D. For *E* and *H*, dashed black curves indicate a normalized $CO_2$ fixation flux of 0.5. (*C*, *F*, and *I*) Overall fluxes of $HCO_3^-$ (*Left subpanels*) and $CO_2$ (*Middle subpanels*) into the chloroplast, normalized by the maximum $CO_2$ fixation flux if Rubisco were saturated, at varying LCIA rates and reversibilities. Negative values denote efflux out of the chloroplast. The inorganic carbon (Ci) species with a positive influx is defined as the Ci source, shown in *Right subpanels*. Axes are the same as *B*, *E*, and *H*.

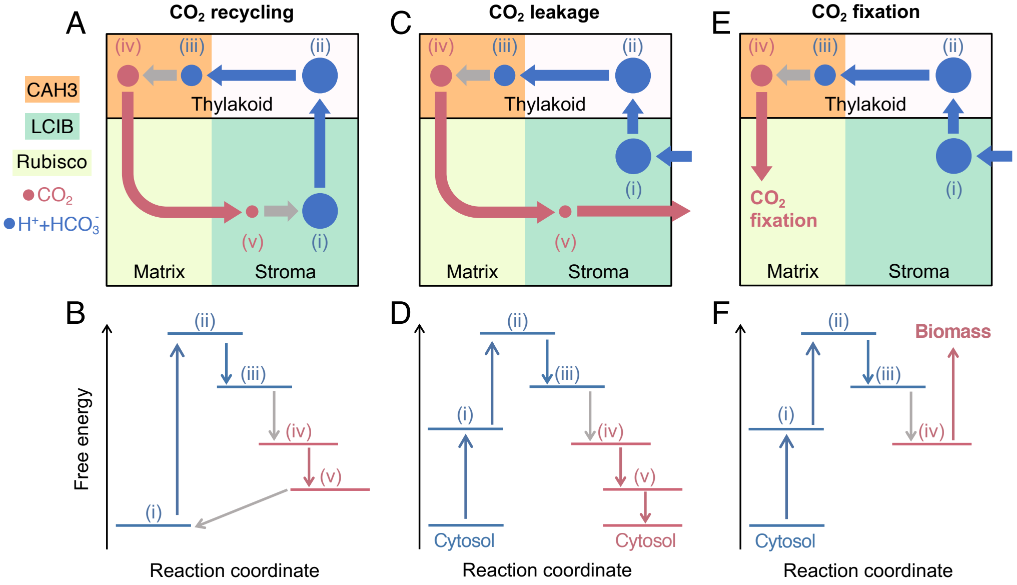


**Fig. S13.** Illustrations of the nonequilibrium processes that cost free energy. Energetic calculation Eq. S46 yields a decomposition of the nonequilibrium fluxes into three processes: (*A*, *B*) $CO_2$ recycling, (*C*, *D*) $CO_2$ leakage, and (*E*, *F*) $CO_2$ fixation (see Sec. II B for details). (*A*, *C*, *E*) Schematics of carbon fluxes. Color code is the same as Fig. 4 except that gray denotes interconversion fluxes between $CO_2$ and $HCO_3^-$. (i) $HCO_3^-$ in the stroma; (ii) $HCO_3^-$ in the thylakoids outside the pyrenoid; (iii) $HCO_3^-$ in the thylakoids inside the pyrenoid; (iv) $CO_2$ in the thylakoids inside the pyrenoid; and (v) $CO_2$ in the stroma. (*B*, *D*, *F*) Schematics of the free-energy change between the states indicated in *A*, *C*, and *E*. Upward arrows denote processes coupled to energy input, including proton pumping across the thylakoid membranes (blue, from (i) to (ii)), $HCO_3^-$ pumping across the chloroplast envelope (blue, from cytosol to (i)), and $CO_2$ fixation and biomass production (red). Downward arrows denote nonequilibrium processes that dissipate free energy, including diffusion/transport of $CO_2$ (red) and $HCO_3^-$ (blue), and reactions catalyzed by carbonic anhydrases (gray).

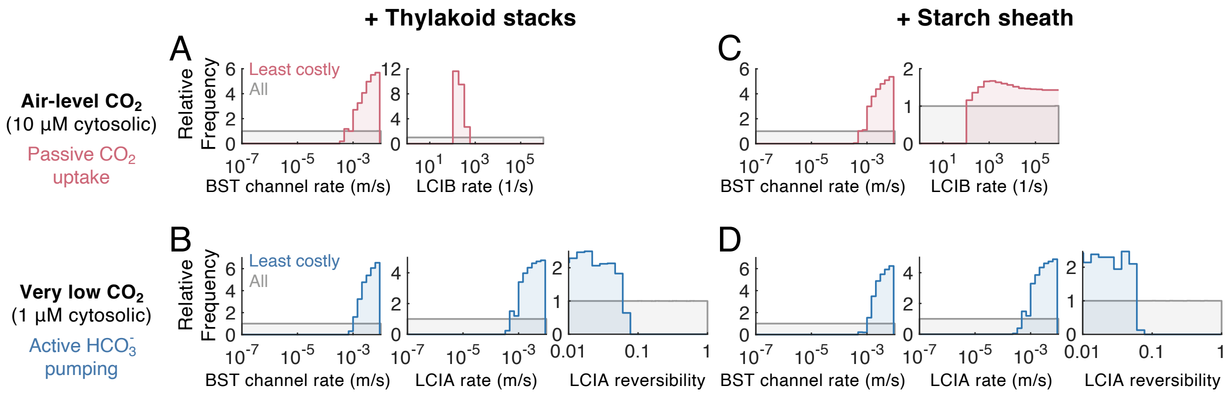


**Fig. S14.** The least energetically costly CCM strategies in (*A*–*B*) a model with thylakoid stacks slowing the diffusion of inorganic carbon in the stroma and (*C*–*D*) a model with an impermeable starch sheath. The same ensembles of strategies as in Fig. 3 are considered. (*A*, *C*) Relative frequency of (*left*) BST channel rate, and (*middle*) LCIB rate for the CCM strategy employing LCIB for passive $CO_2$ uptake and no $HCO_3^-$ transport across the chloroplast envelope under air-level $CO_2$ (10 µM cytosolic). (*B*,*D*) Relative frequency of (*left*) BST channel rate, and (*middle*) the rate and (*right*) reversibility of $HCO_3^-$ transport across the chloroplast envelope by LCIA, for the CCM strategy employing no LCIB and active $HCO_3^-$ pumping across the chloroplast envelope under very low $CO_2$ (1 µM cytosolic). In all panels, red or blue denotes parameters underlying the least costly CCM strategies (Fig. 3 solid curves) that achieve normalized $CO_2$ fixation flux larger than 0.5. Gray denotes all tested parameters.

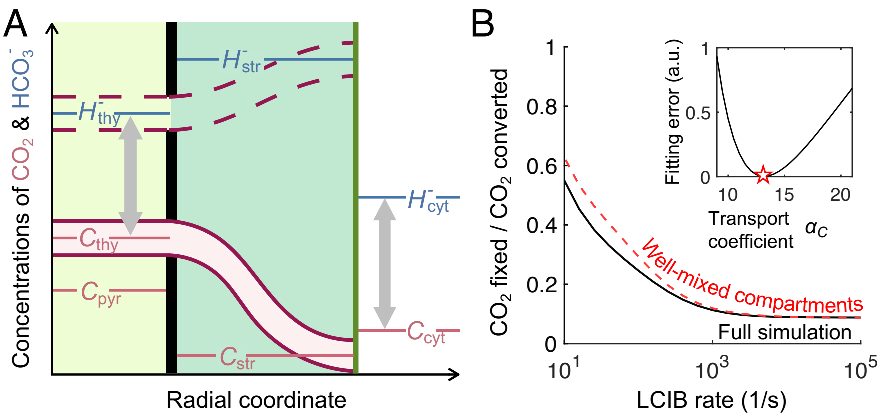


**Fig. S15.** A well-mixed compartment model of the CCM. (*A*) Schematic of the model. Diffusion is assumed to be fast within each compartment (see Sec. III for details). Thus, concentrations of $CO_2$ (red) and $HCO_3^-$ (blue) are represented by horizontal lines, i.e., no gradients. Dashed purple curves indicate that $HCO_3^-$ transport across the thylakoid membrane is fast. Gray arrows indicate species that are in fast equilibrium. Color code is the same as Fig. 1. (*B*) $CO_2$ fixed per $CO_2$ converted from $HCO_3^-$ by CAH3 at varying LCIB rates, computed from the full simulation (black solid curve) and from the well-mixed compartment model (red dashed curve). In both calculations, $HCO_3^-$ transport across the chloroplast envelope is negligible. For the full simulation, we set $\kappa_{\mathrm{thy}}^{H^-} = 10^{-2}$ m/s to approximate the fast $HCO_3^-$ transport across the thylakoid membrane. For the well-mixed compartment model, the $CO_2$ transport coefficient is $\alpha_{\mathrm{thy}}^{C} = 13.0$. *Inset:* Fitting error, defined as the rescaled mean squared difference between the two curves in the main panel when LCIB rate is larger than $10^3$ s$^{-1}$, is plotted against the transport coefficient $\alpha_{\mathrm{thy}}^{C}$ (see Sec. III A). The red star indicates a value of $\alpha_{\mathrm{thy}}^{C} = 13.0$ minimizes the fitting error.

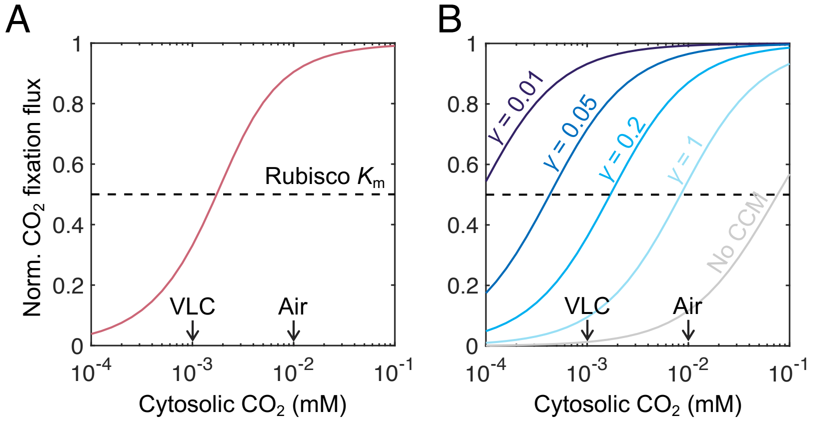


**Fig. S16.** Two extreme CCM strategies are compared under various $CO_2$ conditions for a model employing an impermeable starch sheath. Normalized $CO_2$ fixation flux under different $CO_2$ conditions (described by cytosolic $CO_2$ concentration $C_{\mathrm{cyt}}$ in the model) for (*A*) a CCM strategy that employs fast LCIB for passive $CO_2$ uptake but no $HCO_3^-$ transport across the chloroplast envelope, and (*B*) a CCM strategy that employs no LCIB and fast $HCO_3^-$ transport of the designated reversibilties $\gamma$ across the chloroplast envelope. Black dashed line denotes Rubisco $K_{\mathrm{m}}$ (see Sec. I B). Black arrows indicate air-level $CO_2$ (Air; 10 μM cytosolic) and very low $CO_2$ (VLC; 1 μM cytosolic) conditions. In *B*, the gray curve indicates when there is no CCM, i.e., when the pyrenoid matrix $CO_2$ concentration is the same as that in the environment.

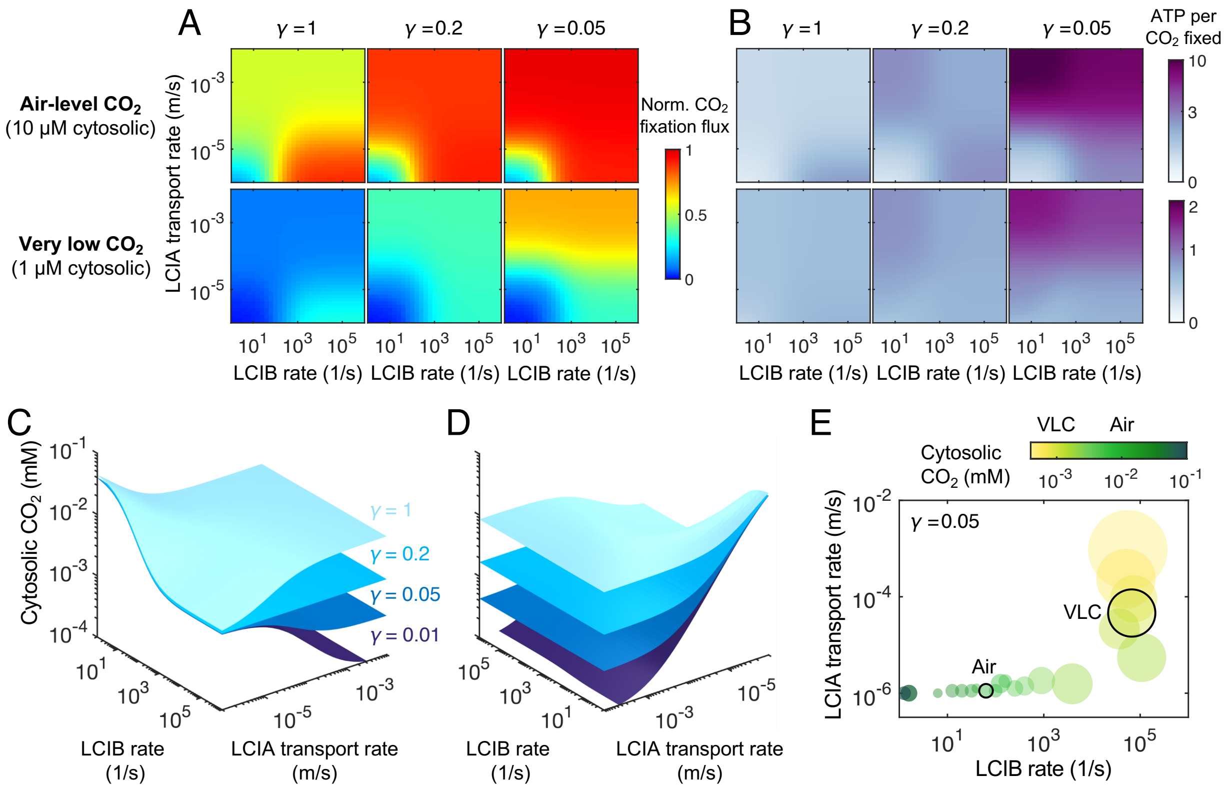


**Fig. S17.** Feasible CCM strategies depend on external $CO_2$ levels in the well-mixed compartment model. (*A*) Normalized $CO_2$ fixation flux and (*B*) ATP spent per $CO_2$ fixed at varying LCIB rates and LCIA transport rates for the designated LCIA reversibilities $\gamma$ under air-level $CO_2$ (top, 10 μM cytosolic) and under very low $CO_2$ (bottom, 1 μM cytosolic). (*C*) Surfaces of normalized $CO_2$ fixation flux of 0.5 for the designated values of $\gamma$. (*D*) Same as *C* but rotated 180 degrees about the $z$ axis. (*E*) The energetically most efficient CCM strategies under varying external $CO_2$ conditions. Circles indicate the regions of parameters in which normalized $CO_2$ fixation flux is larger than 0.5 and the energetic cost is no more that 5% larger than the minimal cost. The center of each circle is chosen as the regional mean on log scale, and the radius of each circle is proportional to the regional area on log scale. Black outlines denote 10 μM (Air) and 1 μM (VLC) cytosolic $CO_2$, respectively.

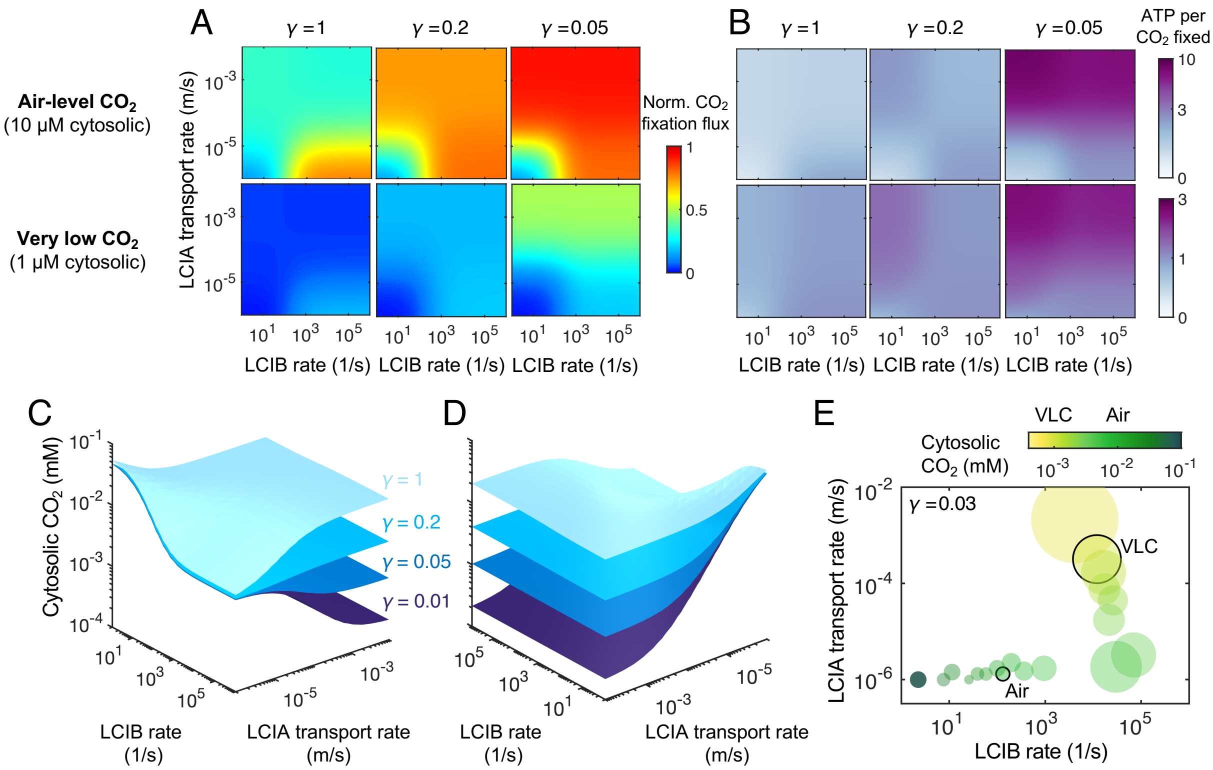


**Fig. S18.** Feasible CCM strategies depend on external $CO_2$ levels in the full reaction-diffusion model. (*A*) Normalized $CO_2$ fixation flux and (*B*) ATP spent per $CO_2$ fixed at varying LCIB rates and LCIA transport rates for the designated LCIA reversibilities $\gamma$ under air-level $CO_2$ (top, 10 µM cytosolic) and under very low $CO_2$ (bottom, 1 µM cytosolic). (*C*) Surfaces of normalized $CO_2$ fixation flux of 0.5 for the designated values of $\gamma$. (*D*) Same as *C* but rotated 180 degrees about the $z$ axis. (*E*) The energetically most efficient CCM strategies under varying external $CO_2$ conditions. Circles indicate the regions of parameters in which normalized $CO_2$ fixation flux is larger than 0.5 and the energetic cost is no more that 5% larger than the minimal cost. The center of each circle is chosen as the regional mean on log scale, and the radius of each circle is proportional to the regional area on log scale. Black outline denotes 10 µM (Air) and 1 µM (VLC) cytosolic $CO_2$. All plots shown are for the full model with an impermeable starch sheath around the pyrenoid matrix.

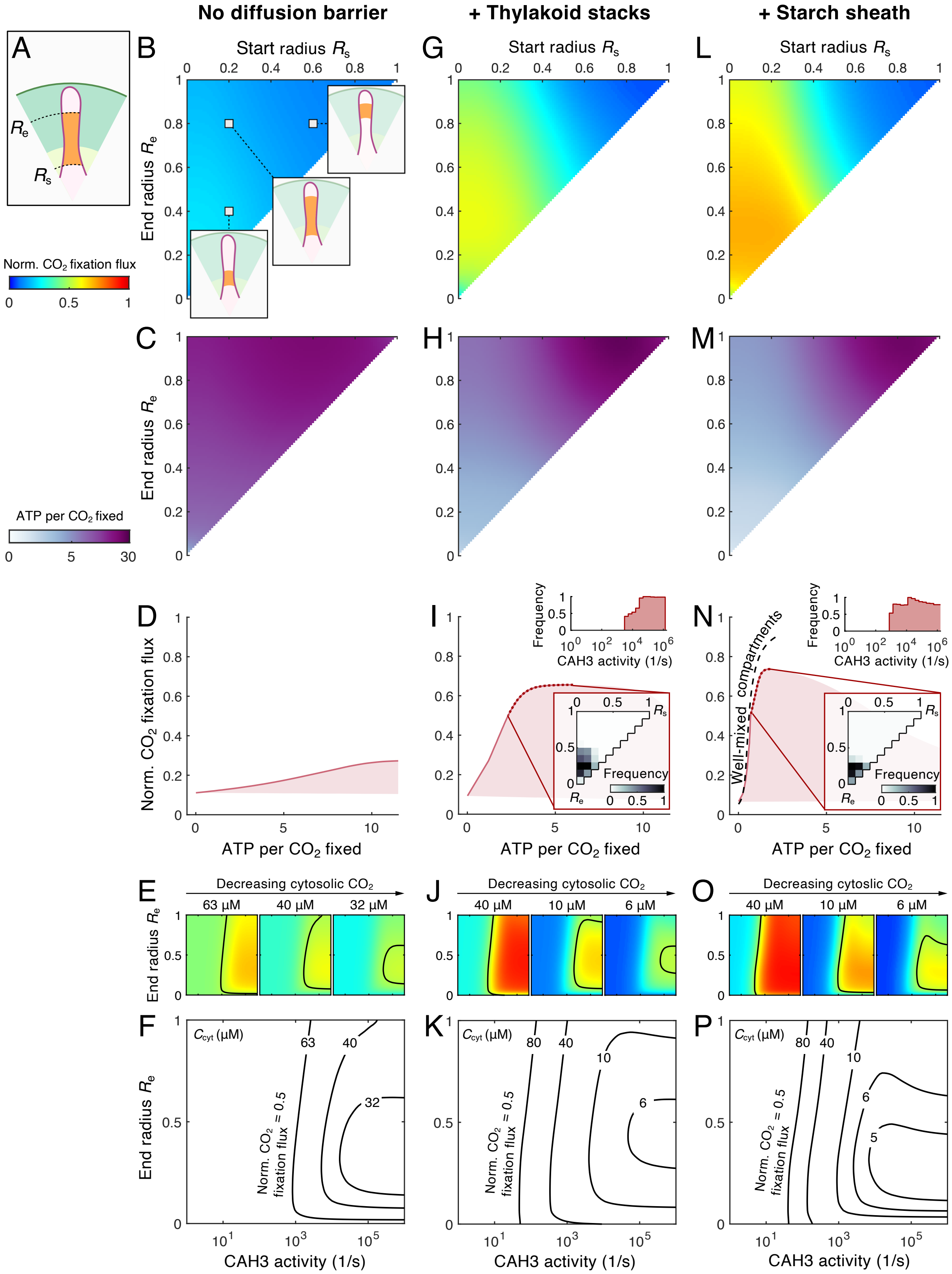
No diffusion barrier
+ Thylakoid stacks
+ Starch sheath
A
B
C
D
E
F
G
H
I
J
K
L
M
N
O
P
Start radius $R_s$
End radius $R_e$
Norm. $CO_2$ fixation flux
ATP per $CO_2$ fixed
Frequency
CAH3 activity (1/s)
Well-mixed compartments
Decreasing cytosolic $CO_2$
Decreasing cytoslic $CO_2$
63 μM
40 μM
32 μM
10 μM
6 μM
$C_{cyt}$ (μM)
Norm. $CO_2$ fixation flux = 0.5

**Fig. S19.** CAH3 localization and activity that enable an effective CCM depend on environmental $CO_2$ levels. (*A*) Schematic of CAH3 (orange) whose localization ranges from start radius $R_s$ to end radius $R_e$. The varying localization patterns are compared in (*B*–*F*) a model without diffusion barriers, (*G*–*K*) a model with thylakoid stacks slowing diffusion of inorganic carbon in the stroma, and (*L*–*P*) a model with an impermeable starch sheath. (*B*,*G*,*L*) Normalized $CO_2$ fixation flux (NCFF) and (*C*,*H*,*M*) ATP spent per $CO_2$ fixed at varying $R_s$ and $R_e$ under air-level $CO_2$ (10 μM cytosolic). Insets in *B* illustrate the localization patterns for the designated parameter values. CAH3 activity, defined by the first-order rate constant in the baseline model ($R_s = 0$, $R_e = R_{pyr}$), is set to be $10^4$ s$^{-1}$. The total number of CAH3 is held constant for a fixed CAH3 activity. (*D*,*I*,*N*) NCFF versus ATP cost per $CO_2$ fixed under air-level $CO_2$ (10 μM cytosolic). Shaded regions show the possible performances by varying CAH3 localization patterns and activities. Solid curves indicate the lowest energy cost for a particular NCFF. Dotted curves indicate the points where NCFF $\geqslant 0.5$, which can be achieved by the activities (*top*) and localization patterns (*bottom*) shown in the insets. Dashed black curve in *N* indicates the optimal CCM performance of a well-mixed compartment model (Fig. 3 and Sec. III A). (*E*,*J*,*O*) NCFF at varying activities and end radii of CAH3 under the designated cytosolic $CO_2$ concentrations. Black curves indicate NCFF = 0.5. (*F*,*K*,*P*) Contours of NCFF = 0.5 under the designated cytosolic $CO_2$ concentrations. In all panels, the passive $CO_2$ uptake strategy (Sec. III B) is considered, i.e., $\kappa_{\mathrm{chlor}}^{H^-} = 10^{-8}$ m/s, LCIB first-order rate constant is $10^3$ s$^{-1}$, $\kappa_{\mathrm{thy}}^{H^-} = 3 \times 10^{-4}$ m/s.

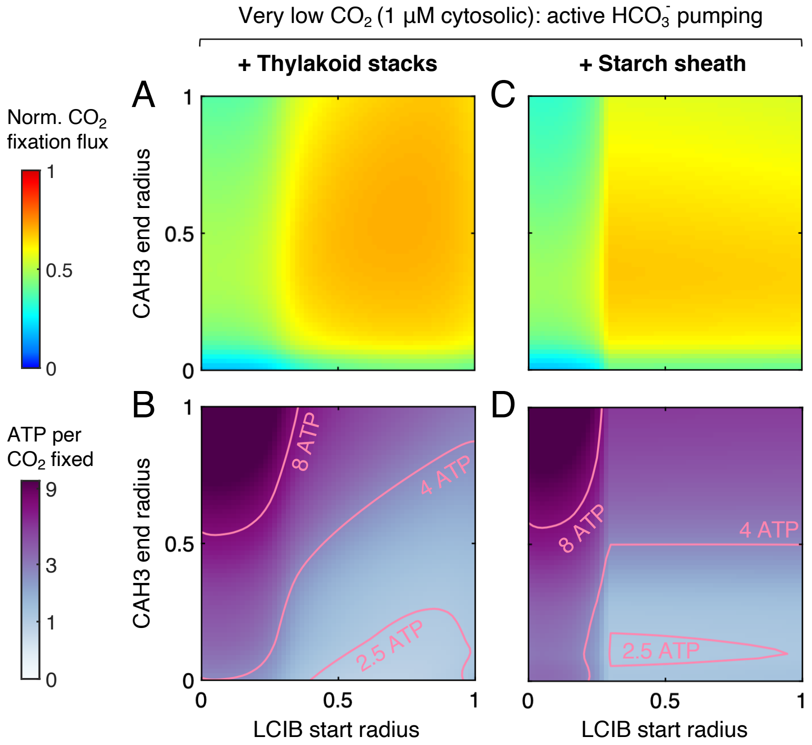


**Fig. S20.** Localization of carbonic anhydrases enhances CCM performance under very low $CO_2$. CAH3 end radius and LCIB start radius are varied in a modeled chloroplast employing the active $HCO_3^-$ pumping strategy under very low $CO_2$, (*A*–*B*) with thylakoid stacks slowing inorganic carbon diffusion in the stroma or (*C*–*D*) with an impermeable starch sheath. (*A*, *C*) Normalized $CO_2$ fixation flux and (*B*, *D*) ATP spent per $CO_2$ fixed when the localizations of carbonic anhydrases are varied. Plots correspond to Figs. 4B–E. The rate and reversibility of $HCO_3^-$ pumping across the chloroplast envelope by LCIA are $\kappa_{\mathrm{chlor}}^{H^-} = 3 \times 10^{-4}$ m/s and $\gamma = 0.01$, respectively. The parameters are chosen such that $CO_2$ fixation flux reaches a level similar to that in Fig. 4.

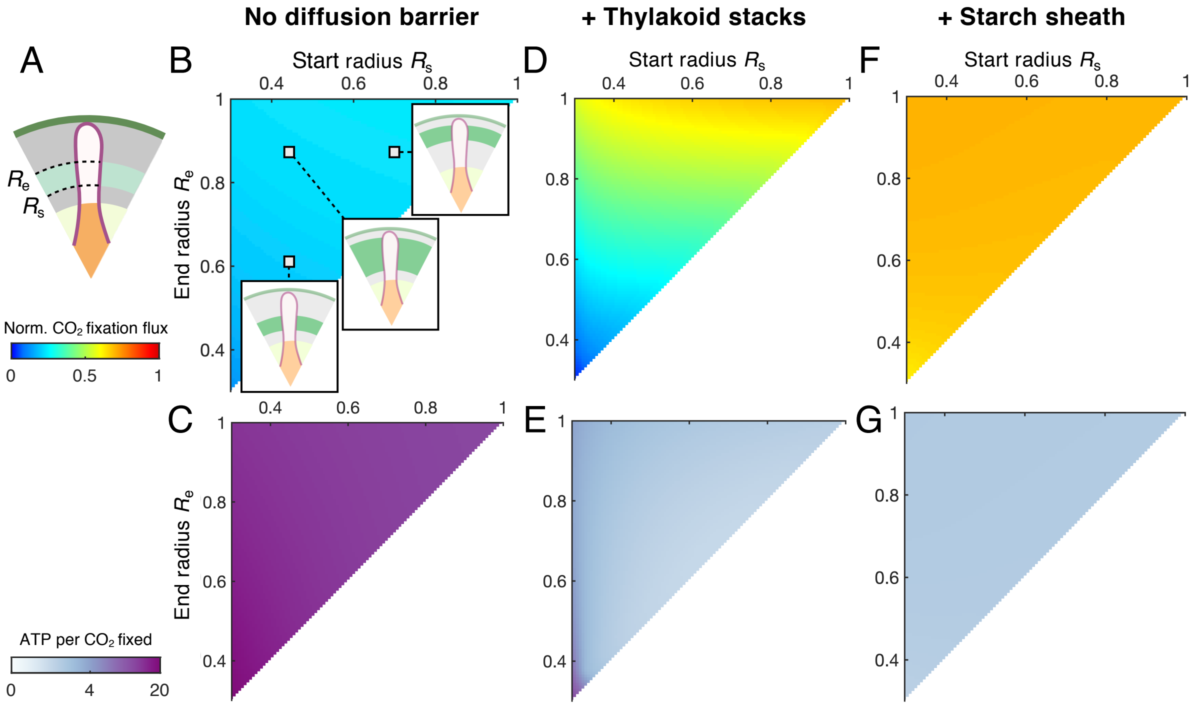


**Fig. S21.** A strong diffusion barrier limits the dependence of CCM activity on LCIB localization for a passive $CO_2$ uptake strategy. (*A*) Schematic of LCIB localization (green) ranging from start radius $R_s$ to end radius $R_e$. Varying localization patterns are compared in (*B–C*) a model without diffusion barriers, (*D–E*) a model with thylakoid stacks slowing diffusion of inorganic carbon in the stroma, and (*F–G*) a model with an impermeable starch sheath. (*B*,*D*,*F*) Normalized $CO_2$ fixation flux and (*C*,*E*,*G*) ATP spent per $CO_2$ fixed at varying $R_s$ and $R_e$ under air-level $CO_2$ (10 μM cytosolic). Insets in *B* illustrate localization patterns for the designated parameter values. LCIB activity, defined by the first-order rate constant in the baseline model ($R_s = R_{pyr}$, $R_e = R_{chlor}$), is $10^3$ $s^{-1}$. The total number of LCIB is held constant for this fixed LCIB activity. In all panels, the $CO_2$-uptake-dominant CCM strategy (Sec. III B) is considered, i.e., $\kappa^{H^-}_{chlor} = 10^{-7}$ m/s, and $\kappa^{H^-}_{thy} = 3 \times 10^{-4}$ m/s.

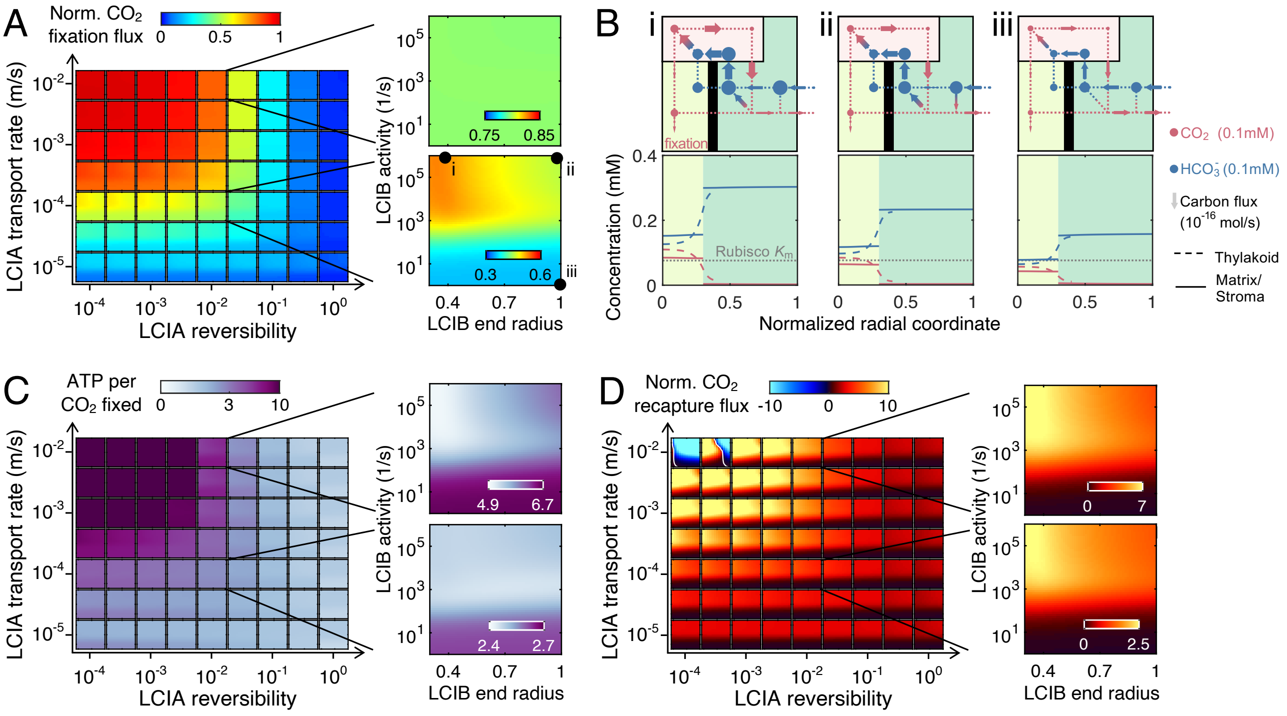


**Fig. S22.** Localization of LCIB around the pyrenoid in a model with a starch sheath allows for $CO_2$ recycling without inducing carbon leakage from the chloroplast when the active $HCO_3^-$ pumping strategy is employed. (*A*) Heatmaps of normalized $CO_2$ fixation flux at varying LCIB end radii and LCIB activities are shown for different rates $\kappa_{\text{chlor}}^{H^-}$ and reversibilities $\gamma$ of $HCO_3^-$ transport across the chloroplast envelope by LCIA. *Right:* Close-up heatmaps for (*top*) $\kappa_{\text{chlor}}^{H^-} = 10^{-2}$ m/s, $\gamma = 10^{-2}$, and for (*bottom*) $\kappa_{\text{chlor}}^{H^-} = 10^{-4}$ m/s, $\gamma = 10^{-2}$. (*B*) Flux diagrams (*top*) and concentration profiles (*bottom*) of inorganic carbon molecules for the parameters indicated in *A*. Color code is the same as Figs. 1C and D. (*C*, *D*) Same plots as *A* for (*C*) ATP per $CO_2$ fixed, and (*D*) normalized $CO_2$ capture flux, defined as the conversion flux of $CO_2$ to $HCO_3^-$ by LCIB normalized by the maximum $CO_2$ fixation if Rubisco were saturated. In *D*, a negative value of normalized $CO_2$ capture flux denotes conversion from $HCO_3^-$ to $CO_2$. For all panels, parameters are the same as in Fig. 5.

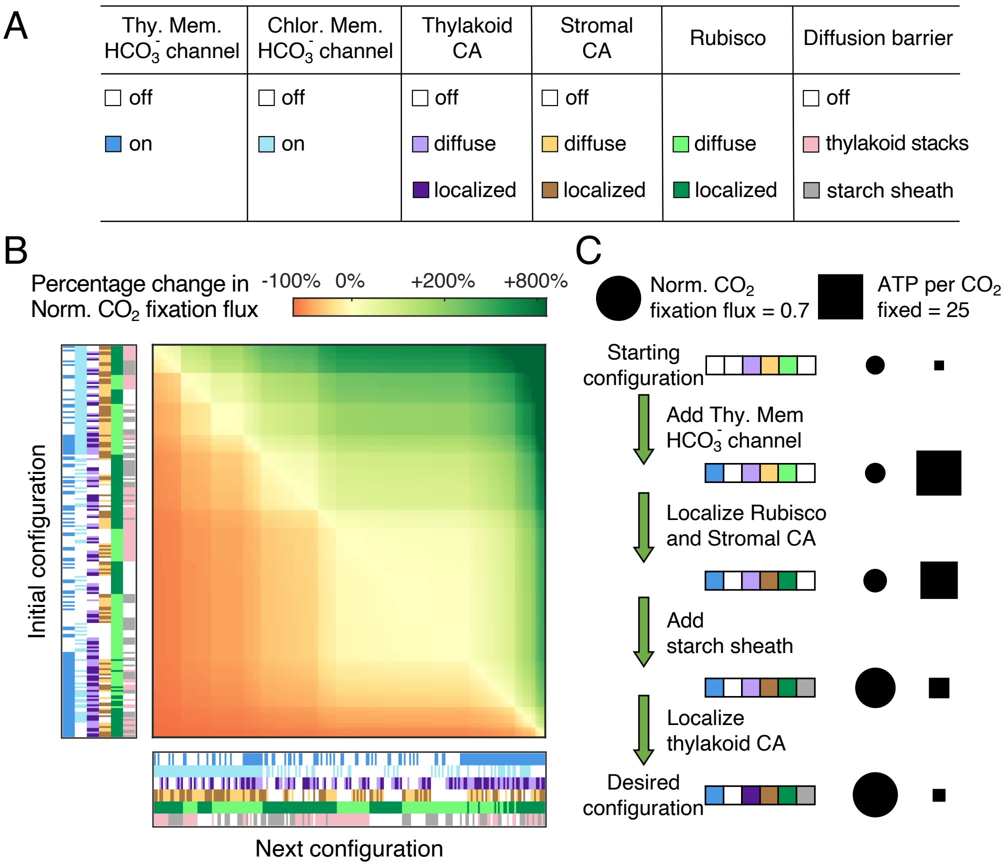


**Fig. S23.** Characterizing various CCM configurations yields feasible engineering paths for installing a CCM into plant chloroplasts. (*A*) CCM components varied in the model. The "off" state denotes rate constants of zero for the $HCO_3^-$ channels and carbonic anhydrases (CAs), and $\kappa_{\text{starch}} = 10$ m/s for the starch diffusion barrier (see Sec. I F). The "on" state denotes $\kappa_{\text{thy}}^{H^-} = \kappa_{\text{chlor}}^{H^-} = 3 \times 10^{-4}$ m/s for the $HCO_3^-$ channels. The thylakoid and stromal CAs, as well as Rubisco, take the baseline activity (Table S4), i.e., total number of enzyme molecules, in both the "diffuse" and "localized" states. The "diffuse" state of the enzymes denotes a start radius $R_s = 0$ and an end radius $R_e = R_{\text{chlor}}$ in their respective compartments (see Sec. I B). The "localized" state of the enzymes denotes the same localization as in the baseline Chlamydomonas model (see Sec. I B). We use the Chlamydomonas thylakoid stacks (Sec. I G), and an impermeable starch at $r = R_{\text{pyr}}$, to model the potential diffusion barriers, regardless of whether Rubisco is localized in a pyrenoid matrix. This yields a combination of 216 CCM configurations. (*B*) Percentage change in the normalized $CO_2$ fixation flux between every pair of configurations described in *A*. Each configuration is indicated by a six-element color code corresponding to *A*. (*C*) Normalized $CO_2$ fixation flux (circle, area in proportion to magnitude) and ATP spent per $CO_2$ fixed (square, area in proportion to magnitude) of all the configurations (indicated by the colored barcode) along one shortest engineering path from the starting configuration representing a typical plant chloroplast to the desired configuration representing a Chlamydomonas chloroplast (see also main Fig. 6). The path above yields the fastest increase in $CO_2$ fixation flux.

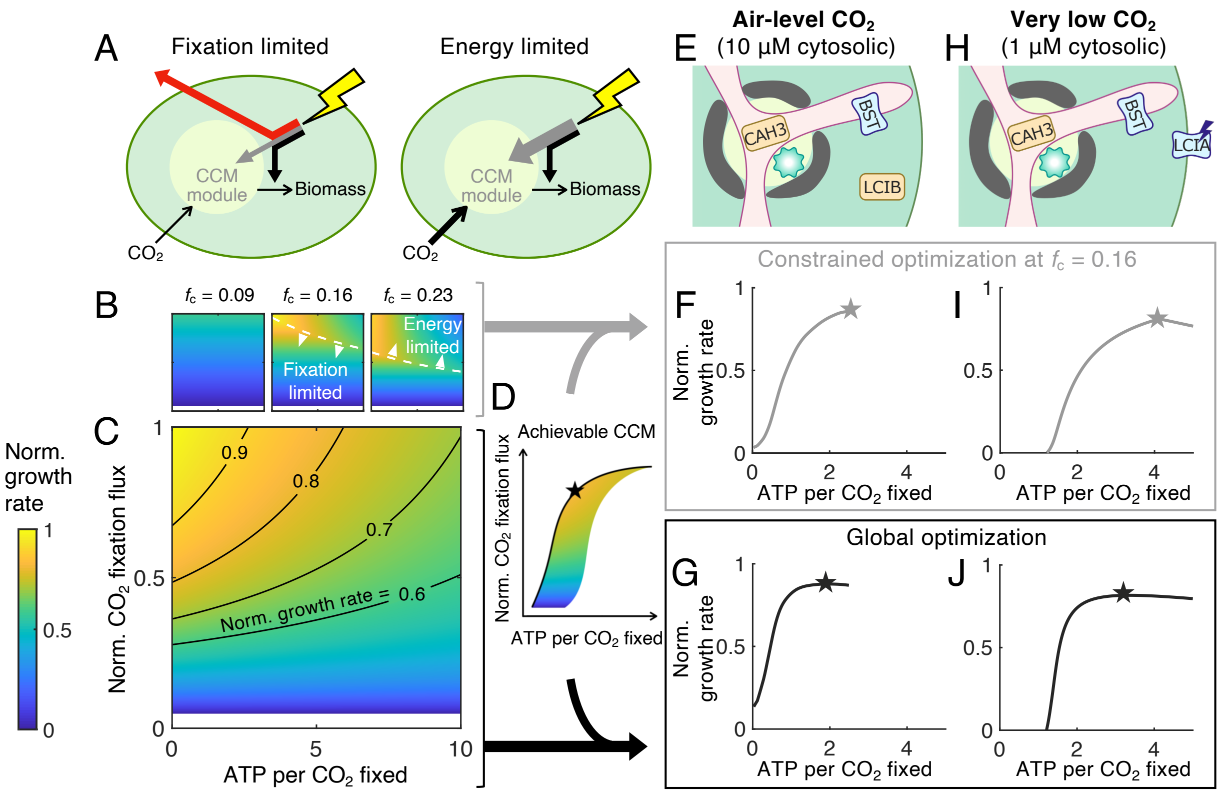


**Fig. S24.** Modeled cell growth rate is determined jointly by the efficacy and energy cost of the CCM. (*A*) Schematics of two scenarios under which cell growth rate is limited. The total energy input from light (lightning symbol) is either used to concentrate and fix $CO_2$ (gray), used to convert $CO_2$ to biomass (black), or dissipated (red). In *left*, the biomass growth rate is limited by $CO_2$ fixation. In *right*, the biomass growth rate is limited by energy. See Sec. II C for details. (*B*, *C*) Normalized growth rate, defined as the growth rate (Eqs. S48 and S50) compared to its global maximum, at varying normalized $CO_2$ fixation flux and ATP cost of the CCM. *B* shows heatmap at the designated CCM biomass fraction $f_c$. White dashed curves indicate the boundary above which growth is energy-limited and below which growth is $CO_2$ fixation-limited. *C* shows the maximum projection over all values $f_c$. (*D*) Schematic of the optimal CCM strategy (denoted by star) which maximizes the growth rate. Note that the shape of the colored region, which indicates all achievable CCM strategies, is schematic, and does not represent any data. The optimal CCM strategies are found by filtering *B* and *C* with the achievable CCM strategies (denoted by the arrows coming from *B*, *C* and *D*) for (*E* – *G*) a modeled chloroplast employing LCIB for passive $CO_2$ uptake under air-level $CO_2$, and (*H* – *J*) a modeled chloroplast employing active $HCO_3^-$ pumping across the chloroplast envelope and no LCIB under very low $CO_2$ (Fig. 3). (*E*, *H*) Schematics of the modeled chloroplast. (*F*, *I*, and *G*, *J*) Normalized growth rates are plotted against ATP spent by the CCM per $CO_2$ fixed along the solid curve indicated in *D* for (*F*, *I*) the designated $f_c = 0.16$ (constrained optimization, gray), and for (*G*, *J*) optimal $f_c$ (global optimization, black).

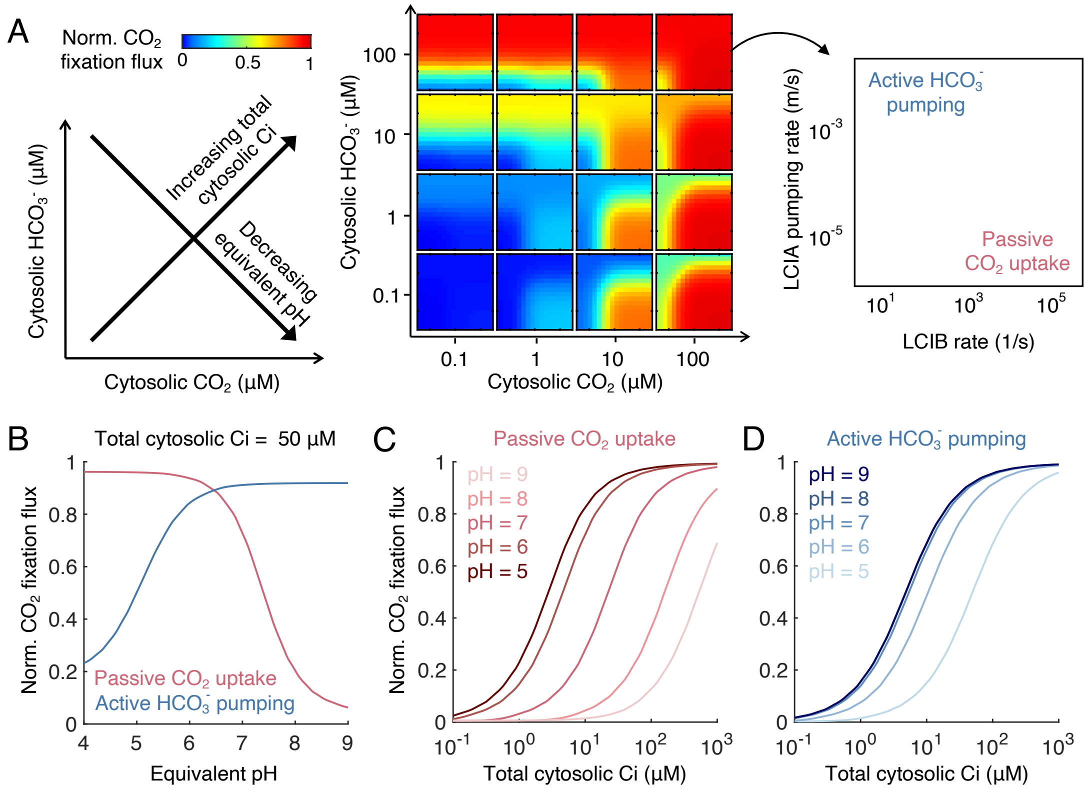


**Fig. S25.** The composition of cytosolic Ci impacts the efficacy of different Ci import strategies. (*A*) Heatmaps of normalized $CO_2$ fixation flux at varying LCIB rate and LCIA pumping rate for different cytosolic $CO_2$ and $HCO_3^-$ concentrations. *Left:* Schematic of the varying cytosolic Ci compositions. Equivalent pH is defined as the pH value for which the $CO_2$ to $HCO_3^-$ ratio corresponds to equilibrium. *Right:* Schematic of one heatmap panel, showing regions where the active $HCO_3^-$ import strategy is employed and where the passive $CO_2$ uptake strategy is employed. (Note that in some cases the activity of LCIA $HCO_3^-$ pumps may result in a net export of $HCO_3^-$ from the chloroplast; this occurs when the external $CO_2$ concentration is high, the LCIB rate is high, and the LCIA pump is fast, since then the $HCO_3^-$ concentration inside the cell becomes much higher than in the cytosol.) (*B*) Normalized $CO_2$ fixation flux under the two Ci import strategies versus equivalent pH of the cytosol. Total cytosolic Ci fixed at 50 μM. (*C*, *D*) Normalized $CO_2$ fixation flux plotted against total cytosolic Ci concentrations at the indicated equivalent pH values, using (*C*) passive $CO_2$ uptake, and (*D*) active $HCO_3^-$ pumping. For all panels, the reversibility of the $HCO_3^-$ pumps is $\gamma = 10^{-1.5}$.

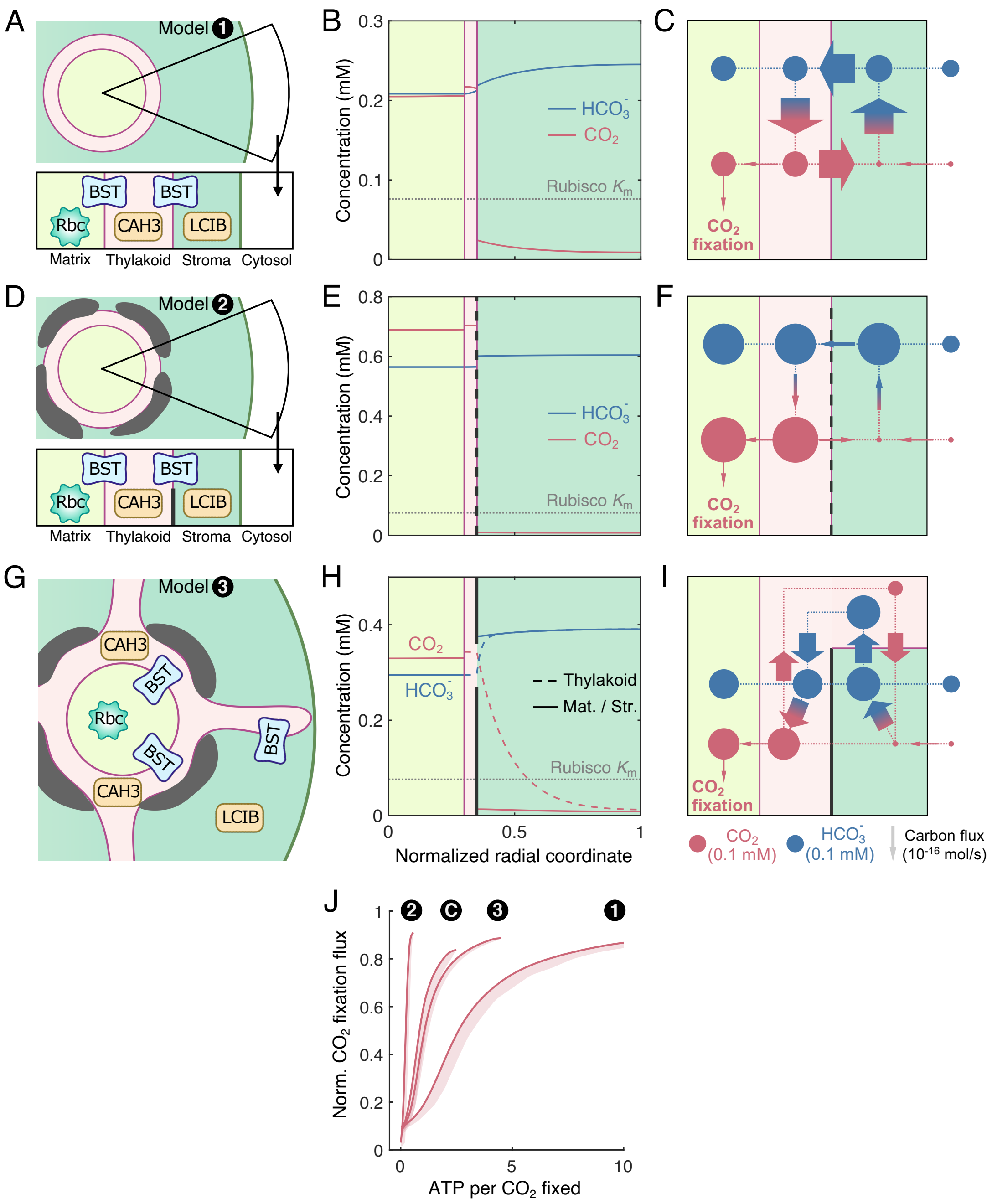


**Fig. S26.** Alternative thylakoid morphologies also support an effective CCM. We consider a pyrenoid matrix localized at the center of the chloroplast, surrounded by a thylakoid sheet, which is (*A*-*C*) surrounded directly by a stroma (model 1), or (*D*-*F*) surrounded by a gapped starch sheath and a stroma (model 2), or (*G*-*I*) surrounded by a gapped starch sheath and extended by cylindrical tubules into the stroma (model 3). See Sec. VI C for details. (*A*, *D*, *G*) Schematics of the modeled chloroplast. Color code as in Fig. 1A. (*B*, *E*, *H*) Concentration profiles of $CO_2$ (red) and $HCO_3^-$ (blue) in all compartments. Vertical gray lines denote the starch sheath. Horizontal dotted gray lines indicate the effective $K_m$ of Rubisco (Sec. I B). Color code as in Fig. 1C. (*C*, *F*, *I*) Net fluxes of inorganic carbon between compartments. The width of arrows is proportional to flux; the area of circles is proportional to the average molecular concentration in the compartment. Color code as in Fig. 1D. (*J*) CCM performance of the passive $CO_2$ uptake strategy under air-level $CO_2$ (10 μM cytosolic), measured by normalized $CO_2$ fixation flux versus ATP spent per CO2 fixed, is compared among models with Chlamydomonas thylakoids (C) and distinct thylakoid morphologies described above (models 1-3). Plots correspond to Fig. 3E. For all panels, parameters are the same as Fig. 1D (Table S1, baseline).

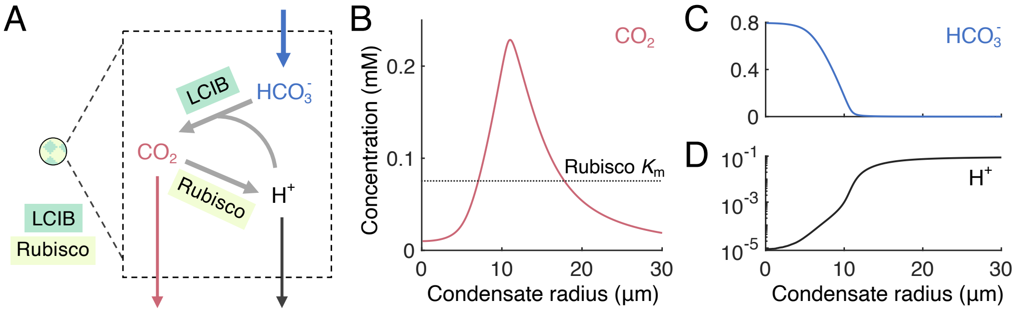


**Fig. S27.** Protons derived from Rubisco $CO_2$ fixation lead to a notable increase in local pH and $CO_2$ availability only in 10 μm CA-Rubisco condensates. (*A*) Schematic of the modeled condensate. $CO_2$ fixation by Rubisco produces protons which in turn drive the conversion of $HCO_3^-$ to $CO_2$ by LCIB. Color code is the same as in Figs. 1 and S11. Black denotes $H^+$. (*B*–*D*) Concentrations of (*B*) $CO_2$, (*C*) $HCO_3^-$, and (*D*) $H^+$ in the condensate at varying condensate radius.

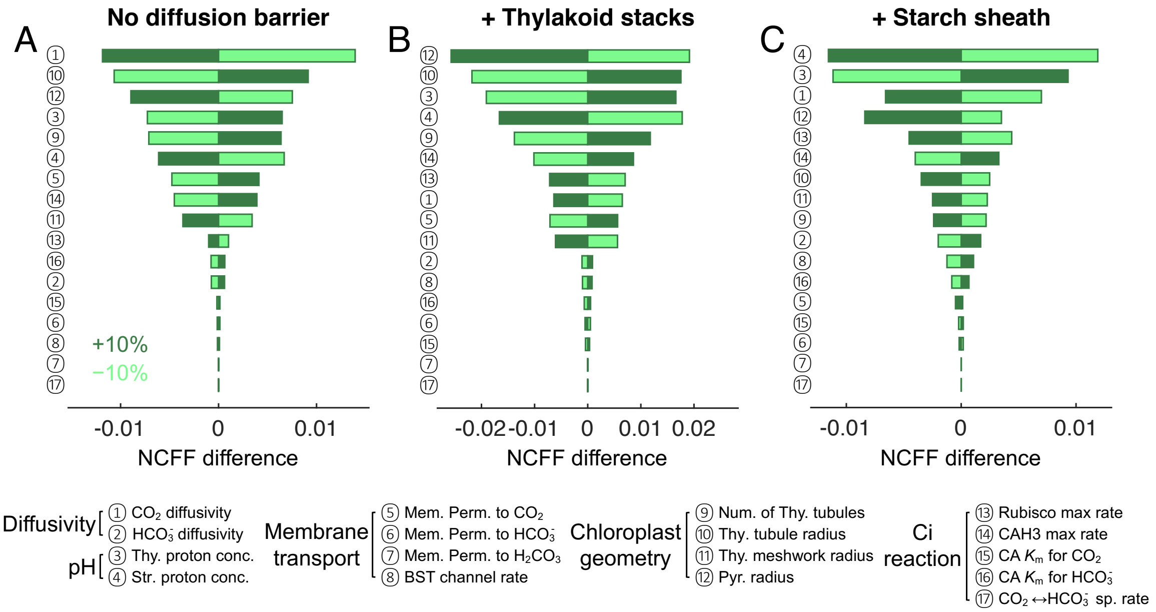


**Fig. S28.** Parameter sensitivity analysis of the multi-compartment reaction-diffusion model. Tornado graphs show the changes in normalized $CO_2$ fixation flux (NCFF, Sec. II A) when the designated parameters (see Table S1) are varied individually by 10% from their baseline values, in (*A*) a model with no diffusion barrier, (*B*) a model with thylakoid stacks slowing inorganic carbon diffusion in the stroma, and (*C*) a model with an impermeable starch sheath. Parameters that impact normalized $CO_2$ fixation flux the most are shown on the top. Sensitivity analyses are done for the CCM strategy employing minimal $HCO_3^-$ transport across the chloroplast envelope ($\kappa_{\text{chlor}}^{H^-} = 10^{-8}$ m/s) and intermediate LCIB activity (first-order rate constant $10^3$ s$^{-1}$) under air-level $CO_2$ (10 μM cytosolic).

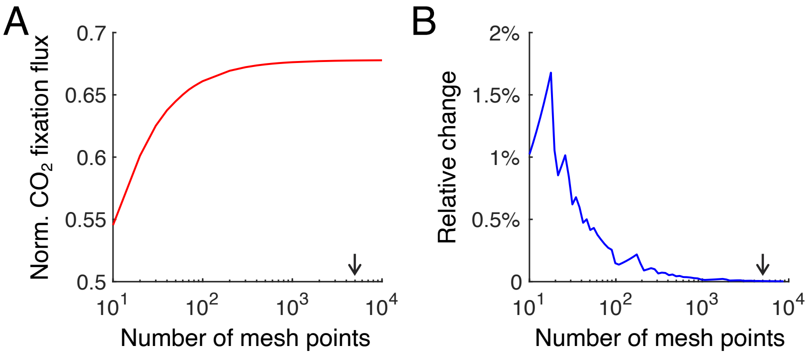


**Fig. S29.** Mesh convergence of the finite-element simulations. The full reaction-diffusion model is simulated for a chloroplast with thylakoid stacks under air-level $CO_2$ (10 μM cytosolic). Parameters are the same as in Fig. 1D. (*A*) Normalized $CO_2$ fixation flux, and (*B*) its relative change upon 10% increase of the number of mesh points, plotted against the number of mesh points. Black arrows indicate simulation results with 5000 mesh points, which we use for all other figures.

**Table S1:** Summary of the notations and parameter values used in the reaction-diffusion model of the CCM.

| Symbols | Descriptions | Values | Ref. |
|---|---|---|---|
| **Compartments** | | | |
| pyr, str, thy | The pyrenoid matrix, stroma, and thylakiod | subscripts | |
| chlor | The chloroplast | subscripts | |
| cyt | The cytosol | subscripts | |
| **Inorganic carbon concentration** | | | |
| $C, H^-, H^0$ | Concentrations of $CO_2$, $HCO_3^-$, and $H_2CO_3$ | | |
| $H$ | Total concentration of $HCO_3^-$ and $H_2CO_3$ | $H^0 + H^-$ | |
| **pH and pKa** | | | |
| $\mathrm{pH}_{\mathrm{pyr}}$ | pH of the pyrenoid in light | 8.0 | [2] |
| $\mathrm{pH}_{\mathrm{str}}$ | pH of the stroma in light | 8.0 | [2, 71] |
| $\mathrm{pH}_{\mathrm{thy}}$ | pH of the thylakoid lumen in light | 6.0 | [72] |
| $\mathrm{pH}_{\mathrm{cyt}}$ | pH of the cytosol | 7.1 | [33] |
| $\mathrm{pKa}_1$ | First pKa of $H_2CO_3$ | 3.4 | [73] |
| $\mathrm{pK}_{\mathrm{eff}}$ | pH at which $C$ and $H^-$ are equal | 6.1 | [74] |
| $\eta$ | Equilibrium $HCO_3^-$ to $H_2CO_3$ concentration ratio | $10^{\mathrm{pH}-\mathrm{pKa}_1}$ | |
| $K^{\mathrm{eq}}$ | Equilibrium $CO_2$ to $HCO_3^-$ concentration ratio | $10^{\mathrm{pK}_{\mathrm{eff}}-\mathrm{pH}}$ | |
| **Chloroplast geometry** | | | |
| $R_{\mathrm{chlor}}$ | Radius of the chloroplast | 3.14 μm | [1, 6] |
| $R_{\mathrm{pyr}}$ | Radius of the pyrenoid | $0.3\,R_{\mathrm{chlor}}$ | [1, 6] |
| $N_{\mathrm{tub}}$ | Number of thylakoid tubules | 40 | [1] |
| $R_{\mathrm{mesh}}$ | Radius of the tubule meshwork | 0.4 μm | [1] |
| $a_{\mathrm{tub}}$ | Cylindrical radius of the thylakoid tubules | 50 nm | [1] |
| $f_{\mathrm{v}}(r)$ | Volume fraction of the thylakoids at radius $r$ | | Sec. I C |
| $f_{\mathrm{s}}(r)$ | Thylakoid surface-to-volume ratio at radius $r$ | | Sec. I C |
| **Diffusion coefficient and permeability** | | | |
| $D^C$ | Diffusion coefficient of $CO_2$ in aqueous solution | $1.88 \times 10^3$ μm²/s | [7] |
| $D^{H^-}$ | Diffusion coefficient of $HCO_3^-$ in aqueous solution | $1.15 \times 10^3$ μm²/s | [8] |
| $D_{\mathrm{eff}}$ | Effective diffusion coefficient in the stroma | | Sec. I G |
| $\kappa^C$ | Membrane permeability to $CO_2$ | 300 μm/s | [29] |
| $\kappa^{H^-}$ | Membrane permeability to $HCO_3^-$ | 0.05 μm/s | [29] |
| $\kappa^{H^0}$ | Membrane permeability to $H_2CO_3$ | 30 μm/s | [4, 30] |
| $\kappa_{\mathrm{starch}}$ | Starch sheath permeability to inorganic carbon | | Sec. I F |
| **Enzyme kinetics of carbonic anhydrases** | | | |
| $V^C_{\mathrm{max}}/K^C_{\mathrm{m}}$ | First-order rate of $CO_2 + H_2O \to HCO_3^- + H^+$ | $10^4$ s$^{-1}$ for CAH3<br>$10^3$ s$^{-1}$ for LCIB | *<br>* |
| $K^C_{\mathrm{m}}, K^{H^-}_{\mathrm{m}}$ | $K_{\mathrm{m}}$ for substrates $CO_2$ and $HCO_3^-$ | 5 mM | [20] |
| $k^C_{\mathrm{sp}}$ | Spontaneous rate of $CO_2 + H_2O \to HCO_3^- + H^+$ | 0.036 s$^{-1}$ | [75] |
| **Enzyme kinetics of Rubisco** | | | |
| $V^C_{\mathrm{max,Rbc}}$ | Maximum rate of $CO_2$ fixation by Rubisco | 15 mM/s | [23, 26] |
| $K^C_{\mathrm{m,Rbc}}$ | $K_{\mathrm{m}}$ of $CO_2$ fixation | 30 μM | [23] |
| $K^O_{\mathrm{m,Rbc}}$ | $K_{\mathrm{m}}$ of oxygenation | 150 μM | [24] |
| $O$ | Dissolved oxygen concentration | 230 μM | [25] |
| $K^{\mathrm{eff}}_{\mathrm{m}}$ | Effective $K_{\mathrm{m}}$ of $CO_2$ fixation at $O = 230$ μM | 76 μM | Sec. I B |
| **Kinetics of $HCO_3^-$ transport** | | | |
| $\kappa^{H^-}_{\mathrm{thy}}$ | Rate of $HCO_3^-$ channels at the thylakoid membranes | $10^{-2}$ m/s | * |
| $\kappa^{H^-}_{\mathrm{chlor}}$ | Rate of chloroplast envelope $HCO_3^-$ transporters | $10^{-8}$ m/s | * |
| $\gamma$ | Reversibility of chloroplast envelope $HCO_3^-$ transport | $0 < \gamma \leqslant 1$ | Sec. I E |

* Values of unknown parameters in the baseline model. See Fig. 2 and Fig. S4 for a thorough parameter scan.

**Table S2:** Summary of the notations and parameter values used in the combined metric calculation (Sec. II C).

| Symbols | Descriptions | Values | Ref. |
|---|---|---|---|
| **Biomass partitioning** | | | |
| $\nu$ | Number of maintenance photons per biomass | 1.67 μmol/g biomass/s | [45] |
| $\chi_C$ | Fraction of carbon in biomass | 0.04 mol C/g biomass | [48] |
| $B_{\mathrm{tot}}$ | Total cellular protein biomass | 20 pg | [55] |
| $B_{\mathrm{c}}$ | Total CCM protein biomass | 1.76 pg | [55] |
| $f_{\mathrm{c}}^0$ | Fraction of biomass in the CCM | 0.086 | [55] |
| $f_{\mathrm{c}}^*$ | Fraction of biomass in the CCM, excess light, $CO_2$ | 0.15 | Sec. II C |
| **Chlamydomonas characteristics** | | | |
| $N_{\mathrm{Chl}}$ | Amount of chlorophyll per cell | 1.5 pg | [54] |
| $a_{\mathrm{Chl}}^*$ | Absorption coefficient of chlorophyll | 10 $\mathrm{m}^2$/(g Chl) | [54] |
| $\mu$ | Turnover number per CCM biomass | 20.3 μmol/g biomass/s | Sec. II C |
| **Fluxes** | | | |
| $\Gamma_C$ | Flux of $CO_2$ fixation by Rubisco | $\mu B_{\mathrm{c}} \Phi_C$ | |
| $\Gamma_O$ | Flux of $CO_2$ lost to oxygenation | $\mu B_{\mathrm{c}} \Phi_O$ | |
| $\Gamma_{\mathrm{b}}$ | Flux of carbon biomass produced | | Sec. II C |
| $\Gamma_{\mathrm{ph}}$ | Photons absorbed per unit time | | |
| $\alpha$ | Proportionality constant, depends on light intensity | $\frac{\Gamma_{\mathrm{ph}}}{B_{\mathrm{tot}}-B_{\mathrm{c}}}$ | |
| $\Gamma_{\mathrm{m}}$ | Absorbed photons used for cellular maintenance | $\nu B_{\mathrm{tot}}$ | |
| $\Phi_C$ | Normalized $CO_2$ fixation flux | | Sec. II A |
| $\Phi_O$ | Normalized $CO_2$ loss from oxygenation | | Sec. II A |
| $\Psi_C$ | Normalized net $CO_2$ fixation flux | $\Phi_C - \Phi_O$ | Sec. II A |
| **Energetic costs and yields** | | | |
| $\epsilon_{\mathrm{b}}$ | Energetic cost of biomass synthesis from $CO_2$ | 20 ATP | [46] |
| $\epsilon_O$ | Energetic cost of oxygenation per $CO_2$ lost | 26.4 ATP | [4, 47] |
| $\epsilon_{\mathrm{ph}}$ | Yield of chemical energy per absorbed photon | 1 ATP | Sec. II C |
| $\epsilon_C$ | Energetic cost of the CCM | | Sec. II B |
| $P_{\mathrm{tot}}$ | Total energy input to the cell | $\epsilon_{\mathrm{ph}}(\Gamma_{\mathrm{ph}} - \Gamma_{\mathrm{m}})$ | Sec. II C |
| **Variables under saturating light** | | | |
| $\Gamma_{c,\mathrm{max}}$ | Flux of $CO_2$ fixation at saturating light | 7.5 fmol C/cell/s | [16, 26] |
| $\Gamma_{\mathrm{ph,S}}$ | Photons absorbed under saturating light | 3 pmol photon/cell/s | Sec. II C |
| $\alpha_S$ | Proportionality constant at saturating light | 76 μmol/g biomass/s | Sec. II C |
| $I_S$ | Saturating light intensity | 200 μmol photon/$\mathrm{m}^2$/s | [53] |

**Table S3:** Summary of the parameters and their values used in the well-mixed compartment model.

| Symbols | Descriptions | Values |
|---|---|---|
| **Starch sheath model (Sec. III A)** | | |
| $H_i, H_o$ | $HCO_3^-$ concentrations inside and outside the pyrenoid | |
| $L_{\text{thy}}$ | Diffusion length scale in the thylakoid tubules | $\approx 0.9$ µm (Fig. S15) |
| $\Delta S_{\text{thy}}$ | Area of thylakoid membranes in the pyrenoid | 8.49 µm$^2$ |
| $\alpha_{\text{thy}}^{H^-}$ | $\frac{2D^H a_{\text{tub}} N_{\text{tub}}}{I[0,R_{\text{pyr}},1,1-f_{\text{v}}]V_{\text{max,Rbc}}^C}$ | 93.2 mM$^{-1}$ |
| $I[0, R_{\text{pyr}}, 1, 1-f_{\text{v}}]$ | Pyrenoid matrix volume | 3.29 µm$^3$ |
| $I[R_{\text{pyr}}, R_{\text{chlor}}, 1, 1-f_{\text{v}}]$ | Stroma volume | 125.5 µm$^3$ |
| $\alpha_{\text{thy}}^{C}$ | $\frac{D^C(\pi a_{\text{tub}}^2/L_{\text{thy}})N_{\text{tub}}}{I[0,R_{\text{pyr}},1,1-f_{\text{v}}]V_{\text{max,Rbc}}^C}$ | 13.0 mM$^{-1}$ (Fig. S15) |
| $\sigma_{\text{thy}}^{C}$ | $\frac{\Delta S_{\text{thy}}\kappa^C}{I[0,R_{\text{pyr}},1,1-f_{\text{v}}]V_{\text{max,Rbc}}^C}$ | 51.6 mM$^{-1}$ |
| $\sigma_{\text{chl}}^{C}$ | $\frac{4\pi R_{\text{chlor}}^2\kappa^C}{I[0,R_{\text{pyr}},1,1-f_{\text{v}}]V_{\text{max,Rbc}}^C}$ | 753 mM$^{-1}$ |
| $\sigma_{\text{chl}}^{H^-}$ | $\frac{4\pi R_{\text{chlor}}^2\kappa_{\text{chlor}}^{H^-}}{I[0,R_{\text{pyr}},1,1-f_{\text{v}}]V_{\text{max,Rbc}}^C}$ | |
| $\beta_{\text{LCIB}}$ | $\frac{I[R_{\text{pyr}},R_{\text{chlor}},1,1-f_{\text{v}}]V_{\text{max,LCIB}}^C/K_{\text{m}}^C}{I[0,R_{\text{pyr}},1,1-f_{\text{v}}]V_{\text{max,Rbc}}^C}$ | |
| **Thylakoid stacks model (Sec. IV B)** | | |
| $\beta_{\text{CAH3}}$ | $\frac{I[0,R_{\text{pyr}},1,f_{\text{v}}]V_{\text{max,CAH3}}^C/K_{\text{m}}^C}{I[0,R_{\text{pyr}},1,1-f_{\text{v}}]V_{\text{max,Rbc}}^C}$ | 43.0 mM$^{-1}$ |
| $L_{\text{str}}$ | diffusion length scale in the stroma | Eq. (S61) |
| $\alpha_{\text{str}}^{C}$ | $\frac{4\pi R_{\text{pyr}}^2(1-f_{\text{v}}\vert_{r=R_{\text{pyr}}})/L_{\text{str}}}{I[0,R_{\text{pyr}},1,1-f_{\text{v}}]V_{\text{max,Rbc}}^C}$ | |

**Table S4:** Efficacy and energetic efficiency of the 216 engineering configurations varying the presence and localization of enzymes, $HCO_3^-$ channels, and diffusion barriers.
